# NgBR controls hepatic adiponectin signaling competence through KAT7-dependent chromatin regulation

**DOI:** 10.64898/2026.05.20.726643

**Authors:** Mohammad Sarif Mohiuddin, Munichandra Babu Tirumalasetty, Wenquan Hu, Rashu Barua, Md. Wahiduzzaman, Mayank Choubey, Gary J. Schwartz, Qing Robert Miao

## Abstract

Adiponectin signaling is essential for hepatic glucose homeostasis, yet the molecular basis of adiponectin receptor responsiveness remains incompletely understood. Here, we identify the Nogo-B receptor (NgBR; NUS1) as a regulator of hepatic adiponectin sensitivity. Across human, cynomolgus monkey, and mouse datasets, hepatic NgBR expression is consistently reduced in obesity-associated diabetes, indicating a conserved metabolic signature. Hepatocyte-specific NgBR deletion abolishes the metabolic effects of the adiponectin agonist AdipoRon, resulting in impaired AMPK activation, persistent gluconeogenesis, and ceramide accumulation. Mechanistically, NgBR loss suppresses KAT7 expression and reduces histone acetylation at AdipoR1 and AdipoR2 promoters, thereby limiting receptor expression. Adeno-associated virus (AAV)-mediated restoration of hepatic NgBR reinstates KAT7-dependent chromatin activation, adiponectin receptor expression, and glucose homeostasis. These findings support a hepatocellular mechanism in which NgBR maintains adiponectin receptor competence and suggest a potential therapeutic strategy for restoring adiponectin responsiveness in metabolic disease.

**Graphical Abstract:** 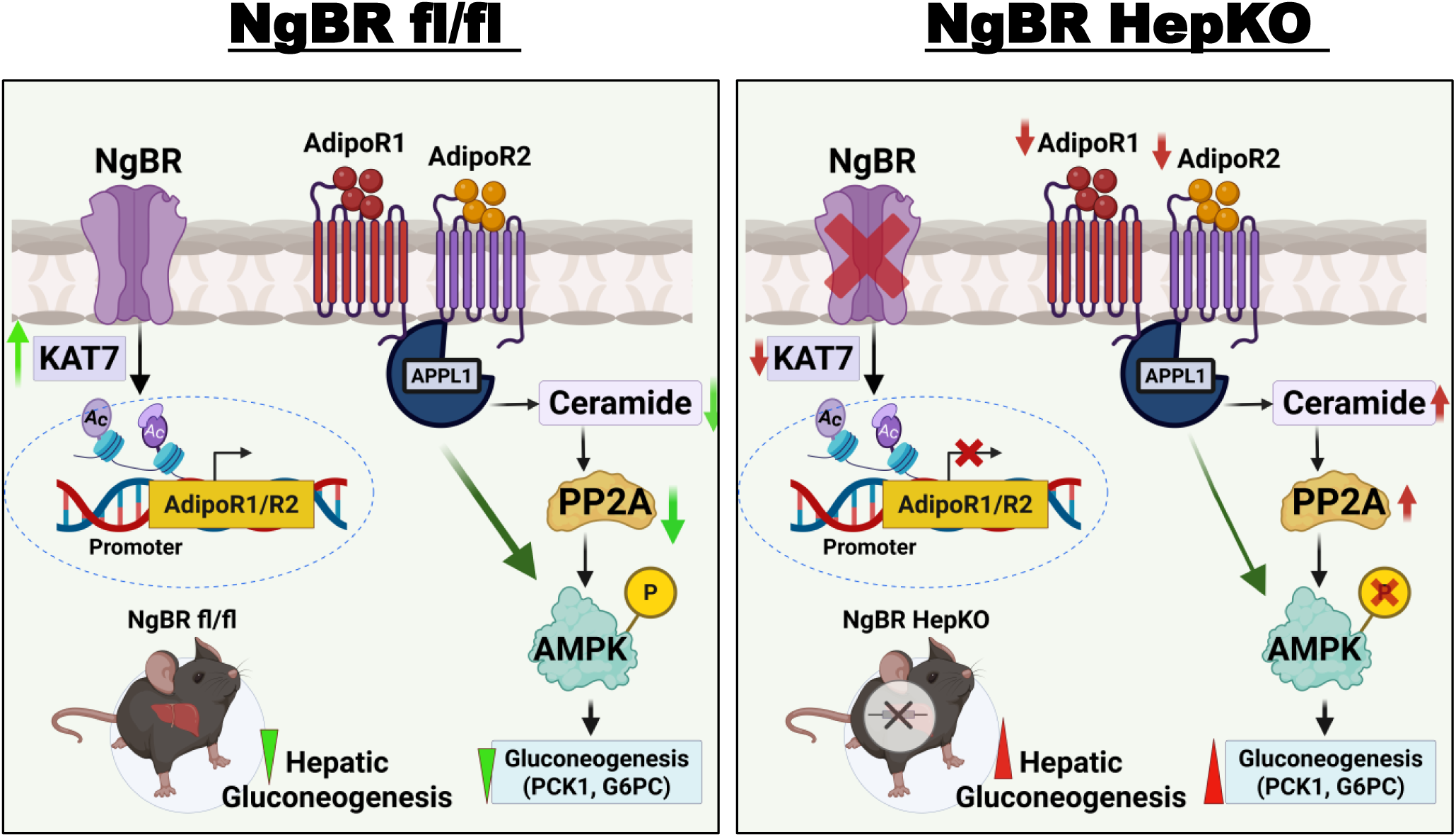

## Main

Obesity and type 2 diabetes (T2D) are among the most prevalent metabolic disorders worldwide, affecting over 460 million individuals and projected to exceed 1.31 billion by 2050^1–3^. These conditions reflect progressive disruption of nutrient and hormonal communication across adipose tissue, liver, and skeletal muscle^4,5^. Adiponectin, an adipocyte-derived hormone, plays an important role in this network by enhancing insulin sensitivity and regulating lipid metabolism.^6–9^ Through its receptors AdipoR1 and AdipoR2, adiponectin activates AMPK and PPARα signaling to promote fatty acid oxidation, suppress hepatic gluconeogenesis, and maintain metabolic flexibility.^10–12^

Despite its abundance, adiponectin signaling is impaired in obesity and T2D, a state termed adiponectin resistance.^13,14^ This condition is characterized by reduced receptor expression and impaired signaling, leading to persistent hepatic glucose production despite preserved or elevated circulating adiponectin.^15,16^ While insulin resistance has well-defined molecular drivers^17,18^, the upstream determinants of adiponectin resistance remain unclear. Pharmacological agonists such as AdipoRon activate adiponectin receptors in experimental models yet exhibit inconsistent metabolic efficacy, suggesting that receptor competence, rather than ligand availability, may limit adiponectin actio^19–21^.

Emerging evidence identifies the Nogo-B receptor (NgBR; encoded by *NUS1*) as a multifunctional regulator of cellular homeostasis. Initially characterized as a receptor for the reticulon family protein Nogo-B (RTN4)^22^, NgBR has since been implicated in diverse processes, including dolichol biosynthesis, intracellular cholesterol trafficking, and protein N-glycosylation.^23^ Despite these established roles, the function of NgBR in metabolic regulation, particularly in the context of endocrine signaling, remains poorly defined. Recent work from our group demonstrated that NgBR can act as a transcriptional regulator by stabilizing the histone acetyltransferase KAT7 (HBO1) and maintaining histone H3/H4 acetylation, thereby preserving chromatin accessibility^24^. These findings raise the possibility that NgBR may regulate metabolic adaptation through epigenetic mechanisms; however, whether this function extends to adiponectin signaling has not been determined.

Here, we identify NgBR as a regulator of adiponectin receptor competence and provide evidence for an epigenetic mechanism underlying hepatic adiponectin resistance. We show that NgBR sustains adiponectin signaling by promoting KAT7-dependent histone acetylation at the *AdipoR1* and *AdipoR2* promoters, thereby maintaining receptor expression and downstream signaling capacity. Loss of hepatic NgBR in metabolic disease suppresses adiponectin receptor transcription, impairs AMPK activation, and drives excessive gluconeogenesis, recapitulating a state of adiponectin resistance. These findings uncover an NgBR–KAT7 regulatory axis that links epigenetic control to adiponectin receptor function and support a model in which this pathway contributes to hepatic adiponectin resistance in T2D.

## Results

### 2.1 Hepatic NgBR loss defines a conserved driver of glucose dysregulation and adiponectin resistance in diabetes

To determine whether NgBR is altered in human metabolic disease, we analyzed liver transcriptomic data from lean, obese, and obese individuals with type 2 diabetes (T2D) (GSE15653).^25^ Hepatic *NUS1* expression was significantly reduced in obese individuals with T2D, whereas *RTN4* expression showed minimal changes across groups (**Fig. 1A**). Analysis of cynomolgus monkey liver RNA-seq datasets (GEO: GSE188418)^26^ confirmed reduced NUS1 expression in obesity-associated diabetes, while RTN4 remained largely unchanged. (**Fig. 1B**). Integration of human single-cell RNA-seq data (GSE202379)^27^ further confirmed reduced hepatocyte-specific expression of *NUS1* in T2D (**Fig. 1C**).

**Fig. 1:**
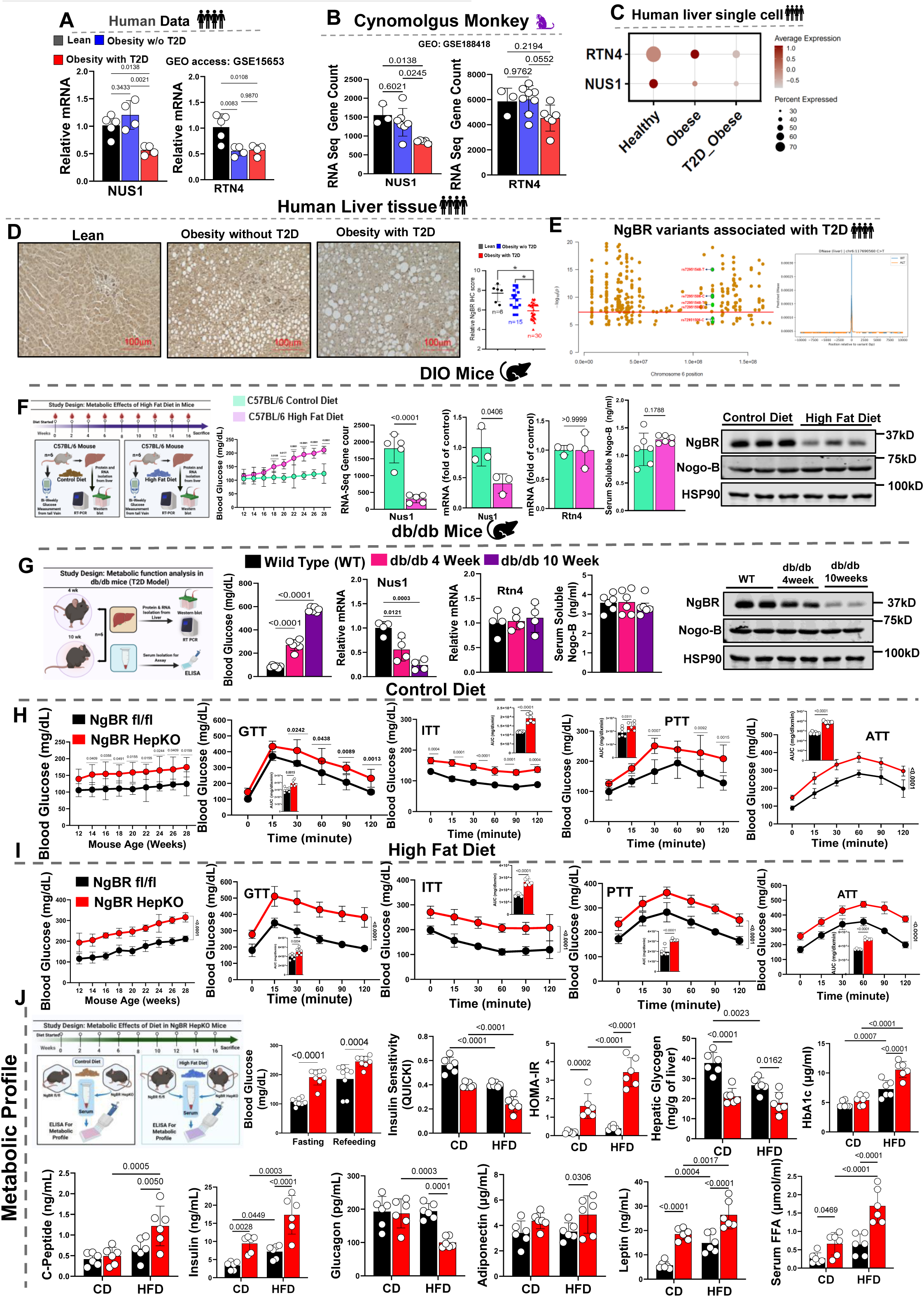
Hepatic NgBR Expression Is Decreased in Obesity-Associated Diabetes and Correlates with Impaired Glucose Homeostasis. **(A)** Human liver transcriptomic data (GEO: GSE15653) showing reduced *NUS1* (NgBR) mRNA in obese individuals with type 2 diabetes (T2D) compared with lean and obese non-diabetic controls, whereas *RTN4* (Nogo-B) shows minimal change (n = 16–20 individuals per group). **(B)** Cynomolgus monkey liver RNA-seq dataset (GSE188418) demonstrated reduced *NUS1* transcripts in obesity-associated diabetes, with no significant alteration in *RTN4* (n = 6 animals per group). **(C)** Human liver single-cell RNA-seq analysis showing hepatocyte-specific reduction of *NUS1* expression in individuals with T2D, whereas *RTN4* remains broadly expressed across cell types. **(D)** Representative human liver immunohistochemistry images showing reduced hepatocellular NgBR protein in T2D livers with steatosis compared with lean and obese non-diabetic controls; quantification shown at right (n = 5–7 individuals per group). **(E)** Regional association plot of the chromosome 6 *NUS1* locus showing T2D-associated variants, including rs72951548 and rs72951506, within a non-coding regulatory region. Predicted liver DNase I hypersensitivity centered on rs72951548 (chr6:117690560 C>T) indicates allele-dependent chromatin accessibility. **(F)** Diet-induced obesity (DIO) model showing reduced hepatic *Nus1* mRNA and protein in high-fat diet (HFD)-fed mice, whereas *Rtn4* expression and circulating soluble Nogo-B remain unchanged (n = 5–7 mice per group). **(G)** db/db mouse model demonstrates progressive reduction of hepatic *Nus1* expression from 4 to 10 weeks of age, with stable *Rtn4* expression. **(H–I)** Longitudinal metabolic testing in NgBR fl/fl and NgBR HepKO mice under control diet (CD; H) or HFD (I), showing fasting hyperglycemia, impaired glucose tolerance (GTT), reduced insulin sensitivity (ITT), and increased pyruvate- (PTT) and alanine-stimulated (ATT) glucose production in NgBR HepKO mice (n = 6–8 mice per group). **(J)** Metabolic profiling showing increased fasting and refeeding glucose, insulin, C-peptide, glucagon, HOMA-IR, HbA1c, leptin, and serum free fatty acids (FFAs), and reduced QUICKI and hepatic glycogen in NgBR HepKO mice (n = 6–8 mice per group). Data are presented as mean ± SD denotes biological replicates (individual humans or animals) as indicated. Statistical tests are described in Methods. Exact *P* values are provided in the figures.

At the protein level, immunohistochemical analysis of human liver sections demonstrated a marked reduction in hepatocellular NgBR expression in T2D, accompanied by steatotic changes (**Fig. 1D**). Genetic analysis of the *NUS1* locus identified T2D-associated variants within non-coding regulatory regions linked to reduced chromatin accessibility (**Fig. 1E**), suggesting that both transcriptional and genetic mechanisms contribute to NgBR downregulation in metabolic disease.^28,29^ These datasets demonstrate that hepatic NgBR downregulation is associated with metabolic disease and suggest that NgBR may regulate hepatic glucose homeostasis.

To validate these findings in experimental systems, we examined hepatic NgBR expression in diet-induced obesity (DIO)^30^ and db/db genetic diabetes models.^31,32^ In DIO mice, chronic high-fat diet feeding led to progressive hyperglycemia accompanied by a significant reduction in hepatic *Nus1* expression, while *Rtn4* expression and circulating Nogo-B levels remained unchanged (**Fig. 1F**). Similarly, db/db mice exhibited marked hyperglycemia and a progressive decline in hepatic NgBR mRNA and protein levels with disease progression (**Fig. 1G**). These findings validate human observations and demonstrate that NgBR downregulation occurs independently of changes in its ligand.

To determine the functional consequences of hepatic NgBR loss, we performed longitudinal metabolic phenotyping in hepatocyte-specific NgBR knockout (NgBR HepKO) mice. Under control diet conditions, NgBR HepKO mice developed progressive fasting hyperglycemia despite no differences in body weight or food intake **(Extended Data Fig.1B)**. Dynamic metabolic testing revealed significant impairments in glucose tolerance, insulin sensitivity, and substrate-driven gluconeogenesis, as assessed by glucose tolerance (GTT), insulin tolerance (ITT), pyruvate tolerance (PTT), and alanine tolerance (ATT) tests (Fig. 1H). These data reveal that hepatic NgBR is required for maintaining glucose homeostasis under physiological conditions.

When challenged with a high-fat diet, the metabolic phenotype of NgBR HepKO mice was markedly exacerbated. Although both genotypes exhibited comparable body weight gain and food intake **(Extended Data Fig.1C),** NgBR HepKO mice displayed severe hyperglycemia, pronounced glucose intolerance, blunted insulin responsiveness, and increased hepatic glucose production (**Fig. 1I**). These results demonstrate that NgBR deficiency renders the liver unable to adapt to nutrient excess.

Consistent with these phenotypes, NgBR HepKO mice exhibited significantly elevated fasting and refeeding blood glucose levels, increased HOMA-IR^33^, reduced insulin sensitivity (QUICKI)^34^, and elevated HbA1c (**Fig. 1J**). Hyperinsulinemia and increased C-peptide levels indicated compensatory β-cell activity, whereas persistent hyperglycemia confirmed severe insulin resistance. Notably, circulating adiponectin levels were preserved or modestly increased, yet failed to improve metabolic outcomes, suggesting impaired hepatic adiponectin responsiveness. In contrast, leptin and free fatty acid levels were markedly elevated, consistent with systemic metabolic dysregulation (**Fig. 1J**).

NgBR deficiency was further associated with widespread metabolic dysfunction, including hypertriglyceridemia, elevated hepatic and serum cholesterol, and increased markers of liver, kidney, and pancreatic injury **(Extended Data Fig. D–G).** Importantly, these abnormalities occurred independently of changes in body composition, as dual-energy X-ray absorptiometry (DEXA) analysis revealed no significant differences in fat mass, lean mass, or bone parameters between genotypes under either dietary condition **(Extended Data Fig. 2A–B).** Similarly, Comprehensive Lab Animal Monitoring System (CLAMS) analysis showed no major differences in energy expenditure, locomotor activity, or respiratory exchange ratio, indicating that altered energy balance does not account for the observed phenotype **(Extended Data Fig. 3A–B).**

Collectively, these findings show that hepatic NgBR downregulation is associated with metabolic disease and that its loss is sufficient to drive hyperglycemia, insulin resistance, and increased gluconeogenesis independent of adiposity or energy expenditure. These data support a role for NgBR in regulating hepatic glucose homeostasis and suggest that its deficiency contributes to adiponectin resistance in type 2 diabetes.

### 2.2 Hepatic NgBR deficiency drives a conserved gluconeogenic transcriptional program and increases hepatic glucose output

To determine whether NgBR downregulation associates with activation of gluconeogenic programs in human disease, we analyzed human single-cell liver datasets. Hepatocytes from obese and type 2 diabetic individuals exhibited increased expression of key gluconeogenic genes, including *PCK1* and *G6PC*, accompanied by reduced *NUS1* expression (**Fig. 2A**)^27^, suggesting a link between NgBR loss and enhanced hepatic glucose production.

**Fig. 2:**
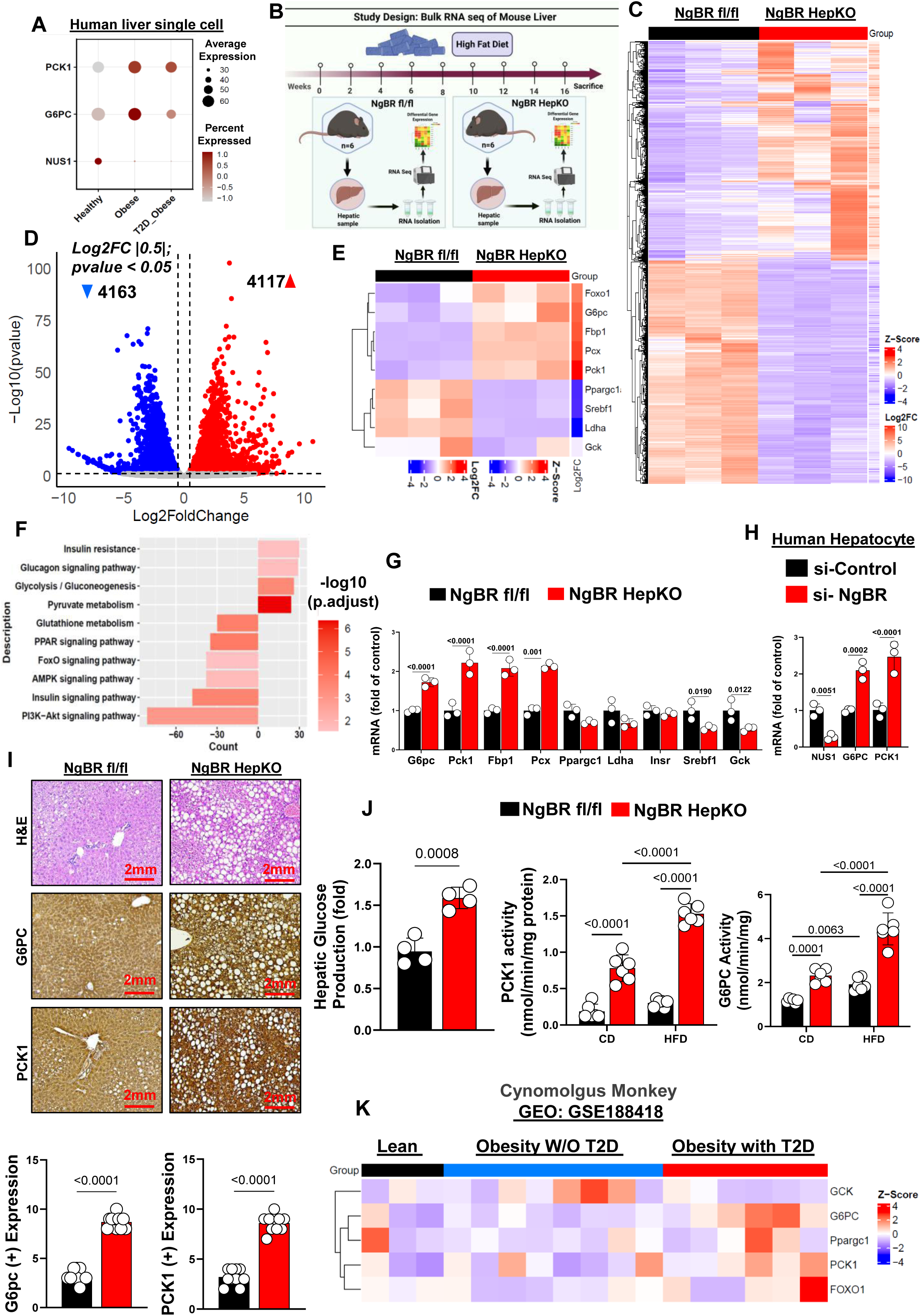
RNAseq analysis of liver from NgBR HepKO mice reveals a higher gluconeogenic gene expression. (**A**) Human liver single-cell RNA-seq analysis showing increased expression of *PCK1* and *G6PC* in hepatocytes from individuals with type 2 diabetes (T2D), alongside reduced *NUS1* (NgBR), compared with healthy and obese non-diabetic individuals. (**B**) Schematic of bulk RNA-seq experimental design in NgBR fl/fl and NgBR HepKO mice fed a high-fat diet (HFD). Livers were collected after 16 weeks for transcriptomic profiling. (**C**) Heatmap of differentially expressed genes showing clear segregation and widespread transcriptional remodeling between NgBR fl/fl and NgBR HepKO livers (Z-score normalized). (**D**) Volcano plot showing significantly altered genes in NgBR HepKO mice (|log_₂_FC| ≥ 0.5; *P* < 0.05), with 4,117 upregulated and 4,163 downregulated transcripts. (**E**) Focused heatmap of gluconeogenic and metabolic regulators showing increased expression of *Foxo1*, *G6pc*, *Fbp1*, *Pcx*, and *Pck1* in NgBR HepKO livers. (**F**) KEGG pathway enrichment analysis showing activation of insulin resistance, glucagon signaling, glycolysis/gluconeogenesis, pyruvate metabolism, and lipid-related pathways in NgBR-deficient livers. (**G**) qPCR validation showing increased expression of *G6pc*, *Pck1*, *Fbp1*, *Pcx*, and *Ppargc1a*, with modest changes in *Ldha*, *Insr*, *Srebf1*, and *Gck* in NgBR HepKO livers (n = 6 mice per group; exact *P* values shown). (**H**) Primary human hepatocytes transfected with si-NgBR showing increased *G6PC* and *PCK1* expression compared with si-control (n = 3 independent experiments; exact *P* values shown). (**I**) Representative H&E staining showing hepatocellular ballooning and lipid accumulation in NgBR HepKO livers. Immunohistochemistry for G6PC and PCK1 shows increased protein abundance; quantification shown below (n = 5–6 mice per group; exact *P* values shown). (**J**) Functional assays showing increased hepatic glucose production, elevated PCK1 enzymatic activity, and increased G6PC activity in NgBR HepKO mice under both control diet (CD) and HFD conditions (n = 6–8 mice per group; exact *P* values shown). (**K**) Cynomolgus monkey liver RNA-seq dataset (GSE188418) showing increased expression of *GCK*, *PCK1*, *FOXO1*, and *G6PC* in obesity-associated diabetes compared with lean controls. Data are presented as mean ± SD denotes biological replicates (animals or independent experiments) as indicated. RNA-seq differential expression analysis was performed using DESeq2, with significance defined as *P* < 0.05 (or adjusted *P* < 0.05 where applicable). Statistical tests are described in Methods. Exact *P* values are provided in the figures.

To define the transcriptional consequences of NgBR deficiency, we performed bulk RNA sequencing of liver tissue from hepatocyte-specific NgBR knockout (NgBR HepKO) and littermate control (NgBR fl/fl) mice following high-fat diet feeding (**Fig. 2B**).^30^ Unsupervised clustering revealed clear segregation between genotypes, indicating extensive transcriptional remodeling in NgBR-deficient livers (**Fig. 2C**). Differential expression analysis revealed widespread transcriptional changes, with prominent enrichment of metabolic pathways (**Fig. 2D**).

Gluconeogenic genes, including *G6pc*, *Pck1*, *Fbp1*, and *Pcx*, were robustly upregulated in NgBR HepKO livers (**Fig. 2E–G**).^33^ In contrast, genes involved in insulin signaling and broader metabolic regulation, such as *Insr* and *Ppargc1a*, remained largely unchanged, indicating a selective activation of gluconeogenic pathways rather than global metabolic suppression. Pathway enrichment analysis further supported this shift, revealing activation of gluconeogenesis, glucagon signaling, and pyruvate metabolism, alongside suppression of AMPK, PPAR, and PI3K–Akt signaling pathways (**Fig. 2F**). These data demonstrate a shift toward a gluconeogenic, insulin-resistant state.

Quantitative PCR analysis confirmed the upregulation of gluconeogenic genes in NgBR HepKO mice (**Fig. 2G**). Consistently, silencing NgBR in primary human hepatocytes increased *G6PC* and *PCK1* expression (**Fig. 2H**), demonstrating that this transcriptional program is hepatocyte-intrinsic. At the tissue level, NgBR-deficient livers exhibited hepatocellular ballooning and lipid accumulation, accompanied by increased G6PC and PCK1 protein expression (**Fig. 2I**). Functionally, hepatic glucose production and enzymatic activities of PCK1 and G6PC were significantly elevated in NgBR HepKO mice under both control and high-fat diet conditions (**Fig. 2J**)^33^, indicating that transcriptional changes translate into increased gluconeogenic flux. Analysis of liver transcriptomic data from cynomolgus monkeys further supported these findings, revealing coordinated upregulation of *G6PC*, *PCK1*, and *FOXO1* in obesity-associated diabetes without consistent changes in *INSR* or *PPARGC1A* expression (**Fig. 2K**)^26^. Together, these results demonstrate that hepatic NgBR deficiency drives a selective gluconeogenic transcriptional program, establishing a metabolic state characterized by increased hepatic glucose production and impaired metabolic flexibility.

### 2.3 Hepatic NgBR is required for adiponectin responsiveness and its loss induces intrinsic adiponectin resistance in vivo

Adiponectin receptor agonists improve metabolic homeostasis by suppressing hepatic glucose production through AdipoR1/2-dependent signaling. To determine whether hepatic NgBR is required for adiponectin responsiveness in vivo, we treated NgBR fl/fl and hepatocyte-specific NgBR knockout (NgBR HepKO) mice with the adiponectin receptor agonist AdipoRon under both control diet (CD) and high-fat diet (HFD) conditions (**Fig. 3A, H**).

**Fig. 3:**
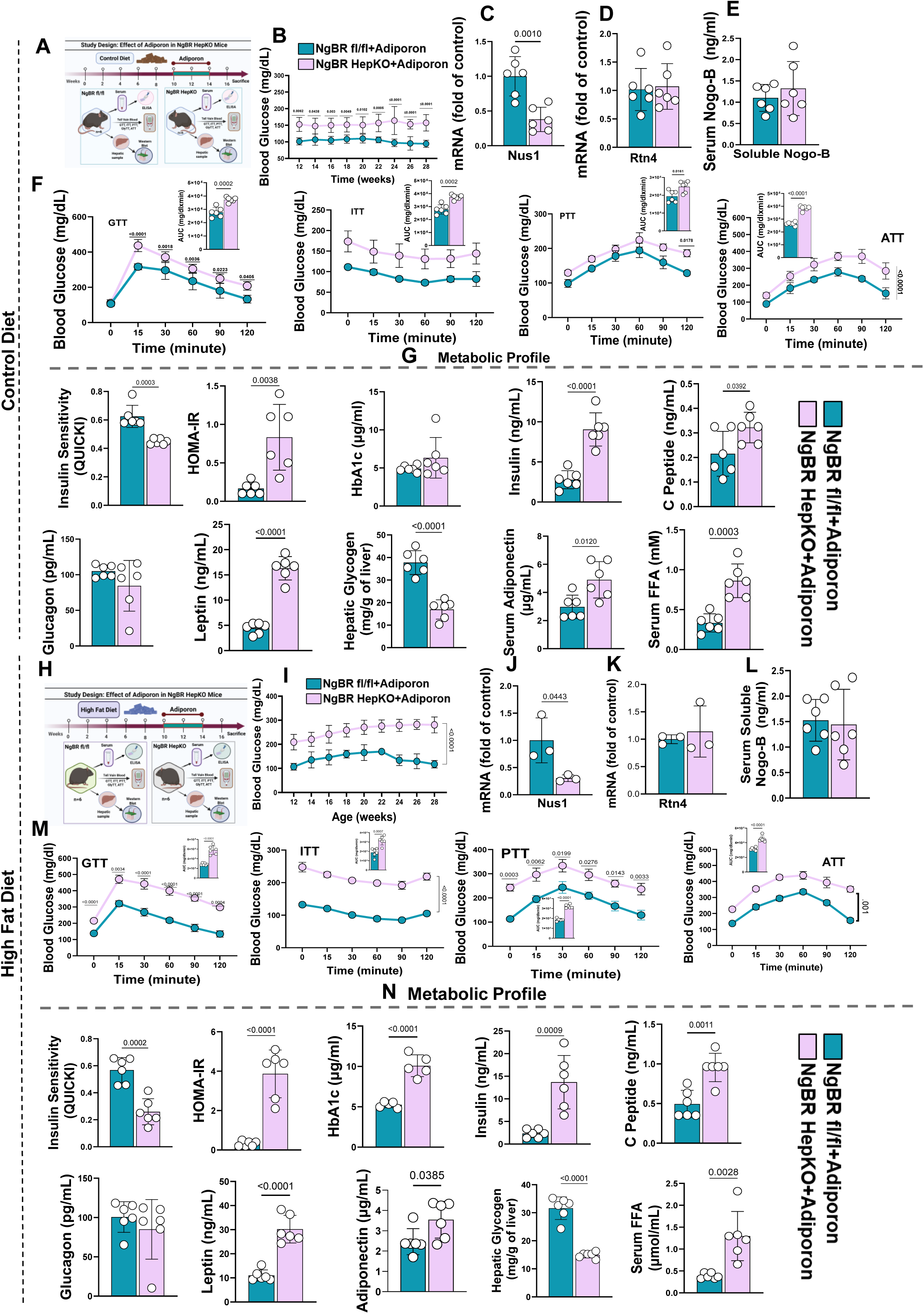
AdipoRon treatment enhances glycemic control in NgBR fl/fl mice but has no effect in NgBR HepKO mice. **(A)** Schematic of the adiponectin receptor agonist (AdipoRon) intervention in NgBR fl/fl and NgBR HepKO mice under control diet (CD). Mice were treated for 4 weeks followed by metabolic phenotyping. **(B)** Longitudinal fasting blood glucose measurements show sustained glucose lowering in AdipoRon-treated NgBR fl/fl mice, with no improvement in NgBR HepKO mice. **(C–D)** Hepatic mRNA expression showing absence of *Nus1* in NgBR HepKO mice and no change in *Rtn4* expression following AdipoRon treatment (n = 5–6 mice per group; exact *P* values shown). **(E)** Circulating soluble Nogo-B levels showing no difference between genotypes or treatment groups (n = 5–6 mice per group; exact *P* values shown). **(F)** Dynamic metabolic testing under CD conditions. AdipoRon improved glucose tolerance (GTT), insulin sensitivity (ITT), and suppressed pyruvate-(PTT) and alanine-stimulated (ATT) glucose production in NgBR fl/fl mice, whereas NgBR HepKO mice showed no response. Insets show area under the curve (AUC) quantification (n = 6–8 mice per group; exact *P* values shown). **(G)** Metabolic profiling under CD showing reduced insulin resistance (HOMA-IR), HbA1c, fasting insulin, C-peptide, and serum free fatty acids (FFAs), together with increased insulin sensitivity (QUICKI) and hepatic glycogen in AdipoRon-treated NgBR fl/fl mice. NgBR HepKO mice showed no improvement (n = 6–8 mice per group; exact *P* values shown). **(H)** Schematic of AdipoRon intervention under high-fat diet (HFD) conditions. **(I)** Longitudinal fasting glucose measurements showing progressive hyperglycemia in NgBR HepKO mice despite AdipoRon treatment, whereas NgBR fl/fl mice exhibited improved glycemic control. **(J–K)** Hepatic mRNA expression showing persistent absence of *Nus1* in NgBR HepKO mice and unchanged *Rtn4* expression across groups (n = 5–6 mice per group; exact *P* values shown). **(L)** Circulating soluble Nogo-B levels showing no effect of AdipoRon treatment (n = 5–6 mice per group; exact *P* values shown). **(M)** Dynamic metabolic testing under HFD conditions. AdipoRon improved GTT, ITT, PTT, and ATT responses in NgBR fl/fl mice but not in NgBR HepKO mice. Insets show AUC quantification (n = 6–8 mice per group; exact *P* values shown). **(N)** Metabolic profiling under HFD showing improvements in HbA1c, fasting insulin, C-peptide, serum FFAs, glucagon, leptin, and hepatic glycogen in AdipoRon-treated NgBR fl/fl mice, with no improvement in NgBR HepKO mice (n = 6–8 mice per group; exact *P* values shown). Data are presented as mean ± SD denotes biological replicates (mice) as indicated. Statistical tests are described in Methods. For time-course experiments (GTT, ITT, PTT, ATT), two-way repeated-measures ANOVA were used. Exact *P* values are provided in the figures.

AdipoRon treatment reduced fasting blood glucose and improved glycemic control in NgBR fl/fl mice, whereas NgBR HepKO mice showed no response under either dietary condition (**Fig. 3B, I**). Hepatic *Nus1* expression remained absent in NgBR HepKO mice and was not altered by treatment, and neither hepatic *Rtn4* expression nor circulating soluble Nogo-B levels changed following AdipoRon administration (**Fig. 3C–E, J–L**), indicating that differences in ligand availability do not account for the observed phenotype. AdipoRon improved glucose tolerance, insulin sensitivity, and suppressed substrate-driven gluconeogenesis in NgBR fl/fl mice under both CD and HFD conditions (**Fig. 3F, M**). In contrast, NgBR HepKO mice exhibited no improvement across glucose (GTT), insulin (ITT), pyruvate (PTT), or alanine (ATT) tolerance tests, indicating failure to respond to pharmacological activation of adiponectin receptors (**Fig. 3F, M; Fig. Extended Data Fig. 4A–B, D–E).**

At the systemic level, AdipoRon markedly improved metabolic parameters in NgBR fl/fl mice, including reduced insulin resistance (HOMA-IR)^35^, increased insulin sensitivity (QUICKI)^34^, and decreased HbA1c, with more pronounced effects under HFD conditions (**Fig. 3G, N; Extended Data Fig. 4C).** These changes were accompanied by normalization of circulating insulin and C-peptide levels, reduced glucagon and leptin, decreased serum free fatty acids, and restoration of hepatic glycogen content, consistent with effective suppression of hepatic glucose output (**Fig. 3G, N**). In contrast, NgBR HepKO mice remained refractory to AdipoRon under both dietary conditions. Hyperglycemia, hyperinsulinemia, elevated HbA1c, and increased circulating lipids persisted, together with failure to restore hepatic glycogen stores (**Fig. 3G, N; Extended Data Fig. 4F)**. Hepatic adiponectin resistance is defined here as the inability of AdipoRon to suppress hepatic glucose production and restore glycogen content despite systemic agonist exposure. Consistent with this definition, NgBR HepKO mice exhibited a complete absence of metabolic correction.

AdipoRon also improved lipid metabolism and organ function selectively in NgBR fl/fl mice, reducing hepatic and circulating triglycerides, lowering serum cholesterol, and improving markers of liver injury. In contrast, NgBR HepKO mice displayed persistent dyslipidemia and elevated liver enzymes, with additional evidence of renal and pancreatic dysfunction under HFD conditions **(Extended Data Fig. 5).** Importantly, these genotype-dependent differences were not attributable to alterations in body composition or energy balance. AdipoRon treatment did not significantly affect body weight or body composition, and no differences in energy expenditure, locomotor activity, or food intake were observed between groups **(Extended Data Fig.6–7)**.

These findings reveal that hepatic NgBR is required for adiponectin-mediated metabolic regulation. While pharmacological activation of adiponectin receptors restores glucose and lipid homeostasis in control mice, NgBR-deficient livers remain unresponsive, establishing a state of intrinsic hepatic adiponectin resistance and identifying NgBR as a regulator of adiponectin efficacy in vivo.

### 2.4 Loss of NgBR disrupts adiponectin receptor expression and abolishes AMPK-mediated suppression of gluconeogenesis

Adiponectin signaling suppresses hepatic gluconeogenesis through coordinated transcriptional and signaling mechanisms. To determine whether NgBR is required for this response, we performed bulk RNA sequencing of liver tissue from NgBR fl/fl and hepatocyte-specific NgBR knockout (NgBR HepKO) mice following AdipoRon treatment (**Fig. 4A).** In NgBR fl/fl mice, AdipoRon induced coordinated transcriptional remodeling of metabolic pathways (**Fig. 4B–D**). In contrast, NgBR HepKO livers exhibited a markedly blunted transcriptional response, indicating failure to engage adiponectin-dependent gene regulation. Genes normally suppressed by adiponectin signaling, including *G6pc*, *Pck1*, *Fbp1*, and *Foxo1*, were effectively downregulated in control livers but remained unchanged or elevated in NgBR-deficient livers (**Fig. 4C**). Pathway analysis further demonstrated suppression of gluconeogenesis, pyruvate metabolism, and inflammatory signaling pathways in NgBR fl/fl mice, whereas these pathways remained active in NgBR HepKO livers (**Fig. 4E**). Quantitative PCR confirmed persistent elevation of gluconeogenic genes despite AdipoRon treatment in NgBR-deficient mice (**Fig. 4F**).

**Fig. 4:**
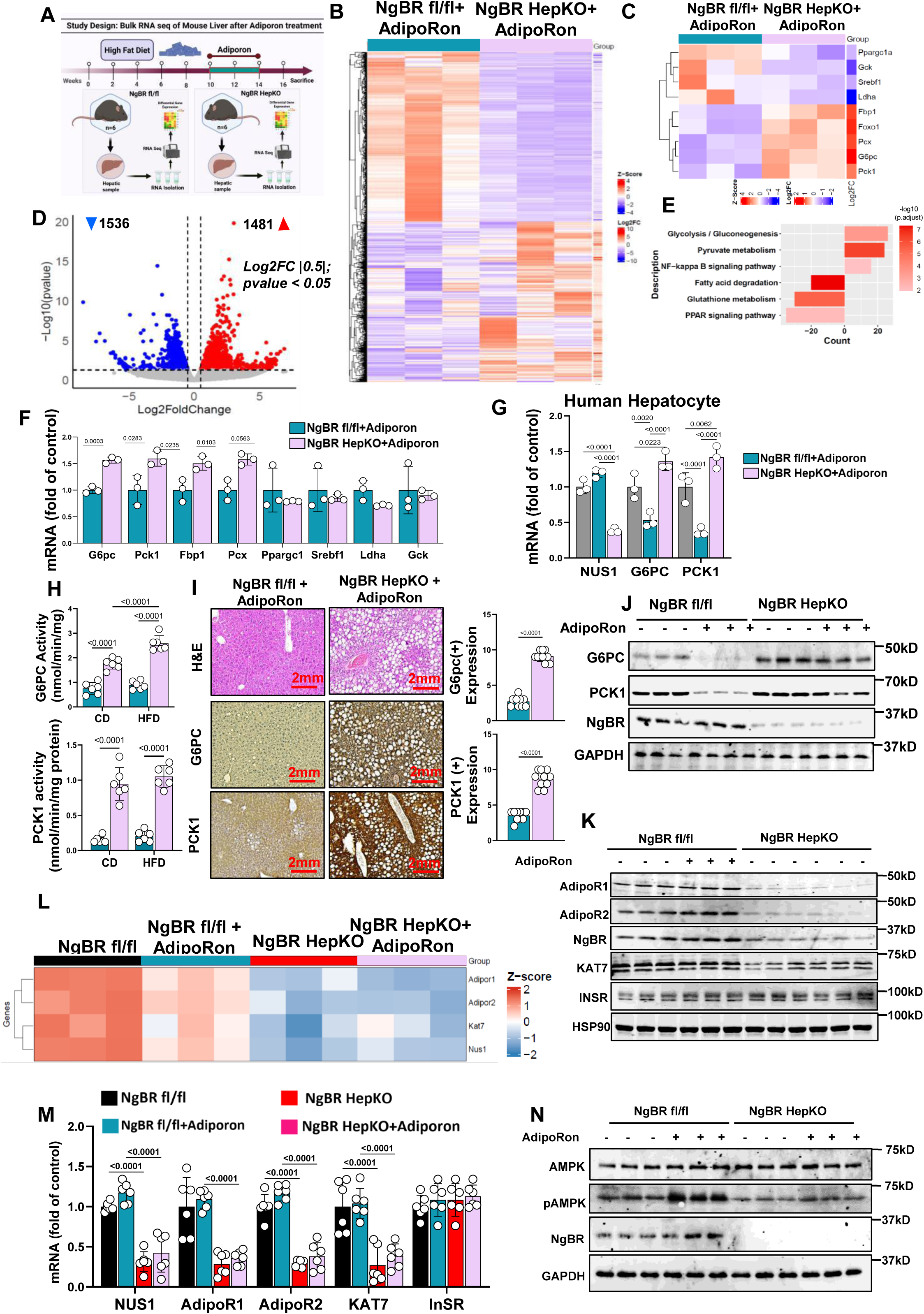
NgBR Deficiency Disrupts AdipoRon-Mediated Repression of Hepatic Gluconeogenesis. **(A)** Schematic of the bulk RNA-seq experimental design in NgBR fl/fl and NgBR HepKO mice following 4 weeks of AdipoRon treatment under high-fat diet (HFD). **(B)** Heatmap of global transcriptomic changes showing robust AdipoRon-induced remodeling in NgBR fl/fl livers, with markedly attenuated responses in NgBR HepKO mice (Z-score normalized). **(C)** Focused heatmap of gluconeogenic and metabolic regulators showing suppression of *Ppargc1a*, *Fbp1*, *Foxo1*, *Pcx*, *G6pc*, and *Pck1* in AdipoRon-treated NgBR fl/fl mice, with minimal or absent changes in NgBR HepKO livers. **(D)** Volcano plot showing differential gene expression following AdipoRon treatment (|log_₂_FC| ≥ 0.5; *P* < 0.05), indicating substantial transcriptional remodeling in NgBR fl/fl livers compared with a blunted response in NgBR HepKO mice. **(E)** KEGG pathway enrichment analysis showing suppression of gluconeogenesis, pyruvate metabolism, fatty acid metabolism, and inflammatory signaling pathways in NgBR fl/fl livers, with persistent pathway activity in NgBR HepKO mice. **(F)** qPCR analysis showing that AdipoRon suppresses *G6pc*, *Pck1*, *Fbp1*, *Pcx*, and *Ppargc1a* expression in NgBR fl/fl livers, whereas NgBR HepKO mice fail to respond (n = 5–6 mice per group; exact *P* values shown). **(G)** Primary human hepatocytes showing that NgBR deficiency abolishes AdipoRon-mediated suppression of *G6PC* and *PCK1* expression (n = 3 independent experiments; exact *P* values shown). **(H)** Enzymatic activity assays showing that AdipoRon reduces G6PC and PCK1 activity in NgBR fl/fl mice under control diet (CD) and HFD conditions, with no effect in NgBR HepKO mice (n = 5–6 mice per group; exact *P* values shown). **(I)** Representative histology (H&E) and immunohistochemistry showing improved hepatic architecture and reduced G6PC and PCK1 protein expression in AdipoRon-treated NgBR fl/fl mice, with persistent steatosis and elevated protein levels in NgBR HepKO livers; quantification shown at right (n = 5–6 mice per group; exact *P* values shown). **(J)** Immunoblot analysis showing that AdipoRon reduces G6PC and PCK1 protein abundance in NgBR fl/fl livers but not in NgBR HepKO mice. **(K)** Immunoblot analysis showing reduced AdipoR1, AdipoR2, and KAT7 expression in NgBR HepKO livers, with no restoration following AdipoRon treatment. **(L)** Heatmap of adiponectin signaling components showing failure of AdipoRon to induce *AdipoR1*, *AdipoR2*, *KAT7*, and *Nus1* expression in NgBR HepKO mice. **(M)** qPCR validation showing reduced expression of *Nus1*, *AdipoR1*, *AdipoR2*, *KAT7*, and *Insr* in NgBR HepKO livers independent of AdipoRon treatment (n = 5–6 mice per group; exact *P* values shown). **(N)** Immunoblot analysis showing that AdipoRon induces AMPK phosphorylation in NgBR fl/fl livers but fails to activate AMPK in NgBR HepKO mice. Data are presented as mean ± SD denotes biological replicates (mice or independent experiments) as indicated. RNA-seq differential expression analysis was performed using DESeq2, with significance defined as *P* < 0.05 (or adjusted *P* < 0.05 where applicable). Statistical tests are described in Methods. Exact *P* values are provided in the figures.

To assess whether this defect is hepatocyte-intrinsic, we silenced NgBR in primary human hepatocytes. NgBR knockdown abolished AdipoRon-mediated suppression of *G6PC* and *PCK1* expression (**Fig. 4G**), indicating a cell-autonomous requirement for NgBR in adiponectin signaling. At the functional level, AdipoRon reduced hepatic glucose production and gluconeogenic enzyme activity in NgBR fl/fl mice, whereas NgBR HepKO mice exhibited persistently elevated glucose output and enzyme activity despite treatment (**Fig. 4H).** Histological analysis showed improved hepatic architecture and reduced lipid accumulation in control mice, while NgBR-deficient livers remained steatotic and retained high G6PC and PCK1 expression (**Fig. 4I–J).** We next examined proximal components of the adiponectin signaling pathway. Expression of both *AdipoR1* and *AdipoR2* was markedly reduced in NgBR HepKO livers, accompanied by decreased expression of the histone acetyltransferase *KAT7* (**Fig. 4L, M).** In contrast, *Insr* expression remained largely unchanged, indicating that the defect is selective to the adiponectin signaling axis. Consistent with this, AdipoRon robustly induced AMPK phosphorylation in NgBR fl/fl livers but failed to activate AMPK in NgBR-deficient mice (**Fig. 4N).**

These data reveal that NgBR is required for adiponectin receptor signaling competence in hepatocytes and downstream signaling. Loss of NgBR disrupts receptor expression, impairs AMPK activation, and prevents suppression of hepatic gluconeogenesis, supporting a state of intrinsic adiponectin resistance in which receptor agonism fails to restore metabolic control.

### 2.5 NgBR–KAT7 epigenetic signaling governs adiponectin receptor expression and metabolic responsiveness

To define the mechanism linking NgBR to adiponectin receptor expression, we investigated the histone acetyltransferase KAT7 as a candidate chromatin effector. In primary hepatocytes, AdipoRon stimulation did not significantly alter expression of *AdipoR1*, *AdipoR2*, or *KAT7* (**Fig. 5A–B**), indicating that receptor abundance is not acutely regulated by ligand stimulation. In contrast, KAT7 silencing markedly reduced *AdipoR1* and *AdipoR2* expression at both mRNA and protein levels, while *Insr* remained unchanged (**Fig. 5A–B**), establishing KAT7 as a selective regulator of adiponectin receptor transcription.

**Fig. 5:**
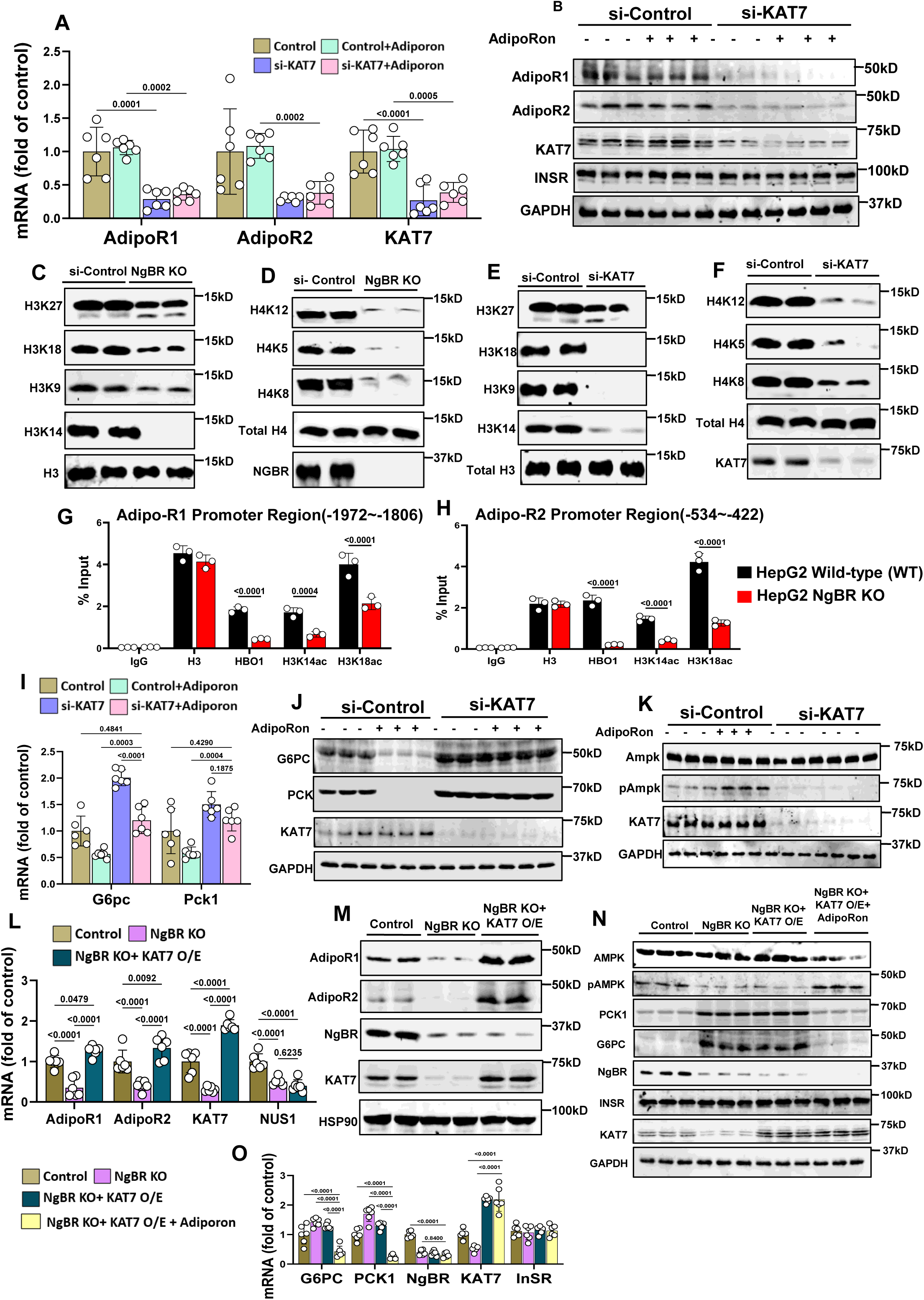
KAT7 Mediates Adiponectin–NgBR Crosstalk to Regulate Hepatic Gluconeogenesis via Histone Acetylation and AMPK Activation. **(A)** qPCR analysis in hepatocytes showing that KAT7 knockdown (si-KAT7) reduces *AdipoR1*, *AdipoR2*, and *KAT7* expression and blunts AdipoRon-induced transcriptional responses, whereas *Insr* remains largely unchanged (n = 3–4 independent experiments; exact *P* values shown). **(B)** Immunoblot analysis confirming that si-KAT7 reduces AdipoR1 and AdipoR2 protein levels without affecting INSR abundance. **(C–D)** Immunoblot analysis of histone modifications in NgBR-deficient cells showing reduced activating histone H3 acetylation marks (H3K27ac, H3K18ac, H3K9ac, H3K14ac) and H4 acetylation marks (H4K12ac, H4K5ac, H4K8ac), with no change in total H3 or H4, indicating impaired chromatin activation in the absence of NgBR. **(E–F)** si-KAT7 hepatocytes showing a similar global reduction in histone acetylation marks, recapitulating the NgBR-deficient phenotype. **(G–H)** ChIP–qPCR analysis showing reduced occupancy of activating histone acetylation marks at *AdipoR1* (−1972 to −1806) and *AdipoR2* (−534 to −422) promoter regions in NgBR-deficient cells, indicating impaired chromatin accessibility (n = 3 independent experiments; exact *P* values shown). **(I)** qPCR analysis showing that KAT7 knockdown prevents AdipoRon-mediated suppression of *G6PC* and *PCK1* expression (n = 3–4 independent experiments; exact *P* values shown). **(J)** Immunoblot analysis showing that si-KAT7 blocks AdipoRon-induced reduction of G6PC and PCK1 protein levels. **(K)** Immunoblot analysis showing that KAT7 knockdown abolishes AdipoRon-induced AMPK phosphorylation without affecting total AMPK levels. **(L)** qPCR analysis showing that KAT7 overexpression restores *AdipoR1*, *AdipoR2*, and *KAT7* expression in NgBR-deficient hepatocytes (n = 3–4 independent experiments; exact *P* values shown). **(M)** Immunoblot analysis confirming that KAT7 overexpression rescues AdipoR1 and AdipoR2 protein levels in NgBR-deficient cells. **(N)** Combined KAT7 overexpression and AdipoRon treatment restores AMPK phosphorylation and suppresses PCK1 and G6PC protein expression in NgBR-deficient hepatocytes. **(O)** qPCR validation showing restoration of *G6PC* and *PCK1* repression and recovery of *KAT7* and *Insr* expression following KAT7 overexpression (n = 3–4 independent experiments; exact *P* values shown). Data are presented as mean ± SD denotes biological replicates (independent experiments) as indicated. Statistical tests are described in Methods. Exact *P* values are provided in the figures.

Consistent with this, NgBR deficiency resulted in a broad reduction in activating histone acetylation marks. NgBR-deficient hepatocytes exhibited decreased levels of H3K14ac, H3K18ac, and H3K9ac, together with reduced H4K5ac, without changes in total histone H3 or H4 (**Fig. 5C–D**). A comparable loss of histone acetylation was observed following KAT7 knockdown (**Fig. 5E–F**), indicating that NgBR deficiency phenocopies KAT7 loss at the chromatin level and that NgBR functions upstream of KAT7-dependent epigenetic regulation.

To determine whether these changes occur at adiponectin receptor loci, we performed chromatin immunoprecipitation analyses. NgBR deficiency significantly reduced H3K14ac and H3K18ac occupancy at regulatory regions within the *AdipoR1* (−1972 to −1806) and *AdipoR2* (−534 to −422) promoters (**Fig. 5G–H).** Expanded profiling across the full promoter regions revealed that total H3 and KAT7 occupancy were largely unchanged, whereas activating histone acetylation marks were consistently reduced at multiple regulatory elements **(Extended Data Fig. 9–10).** These findings demonstrate that NgBR does not affect chromatin loading but instead regulates site-specific histone acetylation at functionally relevant loci controlling adiponectin receptor transcription.

Functionally, KAT7 was required for adiponectin-mediated suppression of gluconeogenesis. KAT7 knockdown increased *G6PC* and *PCK1* expression and abolished their repression by AdipoRon (**Fig. 5I–J**). In parallel, loss of KAT7 impaired AdipoRon-induced AMPK phosphorylation (**Fig. 5K**), linking chromatin-level regulation of receptor expression to downstream signaling output.

To establish causality, we restored KAT7 expression in NgBR-deficient hepatocytes. KAT7 overexpression rescued *AdipoR1* and *AdipoR2* expression at both transcriptional and protein levels (**Fig. 5L–M**). This restoration reinstated AdipoRon-induced AMPK activation and suppressed *G6PC* and *PCK1* expression (**Fig. 5N–O**), demonstrating that KAT7 is sufficient to restore adiponectin signaling in the absence of NgBR. These findings were further supported in HepG2 models, where NgBR deletion or knockdown reduced *AdipoR1*, *AdipoR2*, and *KAT7* expression and prevented AdipoRon-mediated suppression of gluconeogenic genes **(Extended Data Fig. 8A–D).** NgBR deficiency also impaired AMPK phosphorylation in response to AdipoRon or recombinant Nogo-B **(Extended Data Fig. 8E–F)** and increasing AdipoRon concentrations failed to restore suppression of *G6PC* and *PCK1* expression **(Extended Data Fig. 8G)**, indicating a primary defect in signaling competence.

Collectively, these data support a role for KAT7 as a chromatin effector that links NgBR to transcription of the adiponectin receptor. NgBR maintains site-specific histone acetylation at *AdipoR1* and *AdipoR2* regulatory regions, thereby preserving receptor expression and enabling downstream AMPK activation. Loss of NgBR disrupts this epigenetic program, resulting in impaired signaling and failure to suppress hepatic gluconeogenesis, whereas restoration of KAT7 reverses these defects, supporting a mechanistic NgBR–KAT7–adiponectin axis that regulates hepatic metabolic homeostasis.

### 2.6 NgBR deficiency sustains ceramide–PP2A signaling and prevents adiponectin-mediated metabolic correction

Ceramide accumulation is a common feature of metabolic dysfunction and has been linked to adiponectin resistance and excessive hepatic glucose production^36,37^. To establish clinical relevance, we analyzed lipidomic data from the All of Us cohort, which revealed elevated plasma ceramide species, including Cer16:0 and Cer22:0, in individuals with type 2 diabetes compared to metabolically healthy controls (**Fig. 6A**). These data are consistent with an association between ceramide accumulation and impaired metabolic homeostasis in humans.

**Fig. 6:**
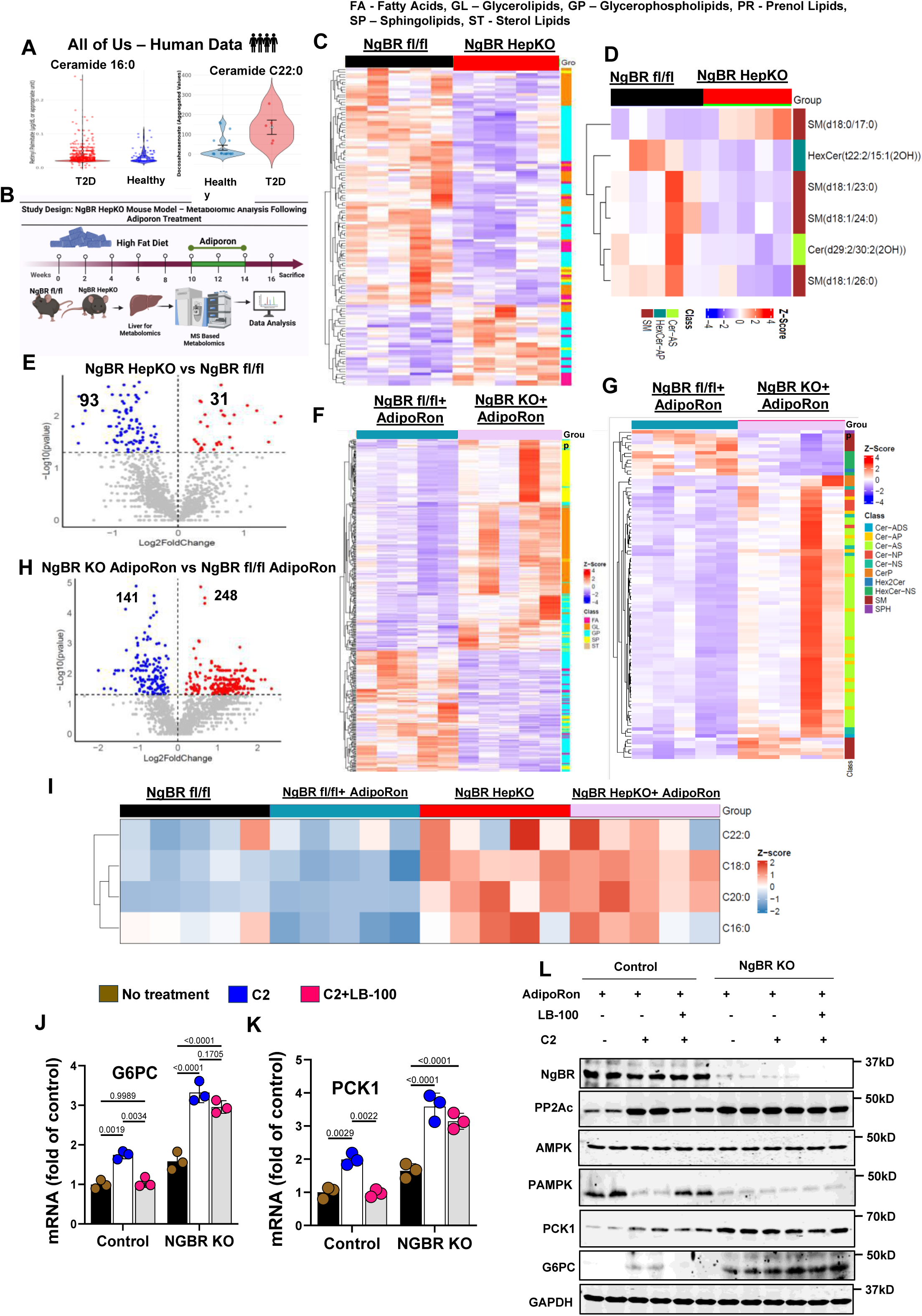
Hepatic lipidomics data reveal a reduction in ceramide levels in NgBR fl/fl mice treated with AdipoRon, while NgBR HepKO mice exhibit either unchanged or elevated ceramide levels in response to the treatment. **(A)** Human plasma lipidomics from the All of Us cohort showing increased circulating ceramide species (Cer16:0 and Cer22:0) in individuals with type 2 diabetes (T2D) compared with metabolically healthy controls. **(B)** Schematic of the NgBR HepKO metabolic study design, including high-fat diet (HFD) feeding, AdipoRon treatment, hepatic lipid extraction, and mass spectrometry–based lipidomic analysis. **(C–D)** Heatmaps of hepatic lipid profiles showing enrichment of ceramide and sphingolipid species in NgBR HepKO livers compared with NgBR fl/fl controls at baseline. **(E)** Volcano plot comparing NgBR HepKO and NgBR fl/fl livers showing differential lipid abundance, with enrichment of ceramide and sphingolipid subclasses among significantly altered species. **(F)** Heatmap showing AdipoRon-induced lipid remodeling in NgBR fl/fl livers, characterized by reductions across multiple ceramide and sphingolipid classes. **(G–H)** In contrast, NgBR HepKO livers show a blunted or altered lipid remodeling response following AdipoRon treatment, with persistence of ceramide-enriched lipid profiles. **(I)** Focused heatmap of long-chain ceramide species showing reduction following AdipoRon treatment in NgBR fl/fl livers, whereas NgBR HepKO mice maintain elevated ceramide levels. **(J–K)** qPCR analysis in primary hepatocytes showing that C2-ceramide induces *G6PC* and *PCK1* expression, with enhanced responses in NgBR-deficient cells. Co-treatment with the PP2A inhibitor LB-100 attenuates ceramide-induced gene expression (n = 3–4 independent experiments; exact *P* values shown). **(L)** Immunoblot analysis showing that C2-ceramide reduces AMPK phosphorylation and increases G6PC and PCK1 protein levels, with stronger effects in NgBR-deficient cells. LB-100 partially restores AMPK phosphorylation and reduces gluconeogenic protein expression. Data is presented as mean ± SD denotes biological replicates (individuals, mice, or independent experiments) as indicated. Lipidomic data were analyzed with multiple testing corrections where applicable. Statistical tests are described in Methods. Exact *P* values are provided in the figures.

To determine whether NgBR regulates adiponectin-dependent ceramide remodeling, we performed mass spectrometry–based lipidomic profiling of liver tissue from NgBR fl/fl and hepatocyte-specific NgBR knockout (NgBR HepKO) mice following AdipoRon treatment under high-fat diet conditions (**Fig. 6B**). Untargeted lipidomic analysis revealed marked differences in sphingolipid composition between genotypes. NgBR HepKO livers exhibited enrichment of ceramide and sphingolipid subclasses relative to controls (**Fig. 6C–E**), indicating a basal shift toward lipotoxic lipid accumulation.

AdipoRon treatment induced coordinated lipid remodeling in NgBR fl/fl mice, characterized by reductions across multiple ceramide and sphingolipid species (**Fig. 6F**). In contrast, NgBR HepKO mice failed to exhibit comparable lipid suppression and instead showed persistent or increased abundance of ceramide species following treatment (**Fig. 6G–H**). Focused analysis of long-chain ceramides further demonstrated that AdipoRon effectively reduced these species in NgBR fl/fl livers but failed to suppress their accumulation in NgBR-deficient mice (**Fig. 6I**). Extended lipidomic profiling confirmed these findings. NgBR fl/fl livers exhibited coordinated suppression of lipid species following AdipoRon treatment, whereas NgBR HepKO livers displayed a blunted and reversed remodeling pattern, characterized by persistent enrichment of ceramide and sphingolipid classes **(Extended Data Fig. 11)**. These data demonstrate that NgBR is required for adiponectin-driven lipid remodeling.

To determine whether ceramide accumulation directly contributes to the metabolic phenotype, we treated hepatocytes with the membrane-permeable ceramide analog C2. NgBR-deficient cells exhibited enhanced induction of *G6PC* and *PCK1* compared to controls (**Fig. 6J–K**), indicating increased sensitivity to ceramide-driven gluconeogenic signaling. Because ceramides suppress AMPK activity in part through PP2A activation^38^, we next assessed the impact of PP2A inhibition. Co-treatment with C2 and the PP2A inhibitor LB-100 suppressed gluconeogenic gene induction in control hepatocytes but showed only partial efficacy in NgBR-deficient cells (**Fig. 6J–K**), suggesting amplification of ceramide–PP2A signaling in the absence of NgBR.

Immunoblot analysis further supported these findings. Ceramide treatment reduced AMPK phosphorylation and increased G6PC and PCK1 protein levels in NgBR-deficient cells, whereas PP2A inhibition partially restored AMPK activation and attenuated gluconeogenic protein expression (**Fig. 6L**). These effects were observed within physiologically relevant conditions, as confirmed by cell viability assays **(Extended Data Fig. 11G–H)**.

Collectively, these results demonstrate that NgBR is required for adiponectin-driven ceramide clearance and suppression of ceramide-induced gluconeogenesis. In the presence of NgBR, adiponectin signaling promotes lipid remodeling, activates AMPK, and restrains hepatic glucose production. In contrast, NgBR deficiency results in persistent ceramide accumulation, enhanced ceramide–PP2A signaling, impaired AMPK activation, and sustained gluconeogenic activity, supporting a lipid-associated mechanism contributing to adiponectin resistance.

### 2.7 Restoration of hepatic NgBR re-establishes KAT7-dependent adiponectin signaling and glucose homeostasis

To determine whether restoring hepatic NgBR is sufficient to rescue adiponectin responsiveness in vivo, we re-expressed NgBR in hepatocytes of NgBR HepKO mice using an AAV8–TBG–NUS1 system followed by AdipoRon treatment (**Fig. 7A, H**). AAV-mediated NgBR reconstitution led to progressive normalization of fasting glycemia under both control diet (CD) and high-fat diet (HFD) conditions, whereas AAV-control–treated NgBR HepKO mice remained persistently hyperglycemic despite identical AdipoRon exposure (**Fig. 7B, I**). Quantitative PCR confirmed robust hepatic re-expression of *Nus1* together with restoration of key adiponectin signaling components, including *KAT7*, *AdipoR1*, and *AdipoR2* (**Fig. 7C, J**), indicating recovery of receptor competence.

**Figure 7.**
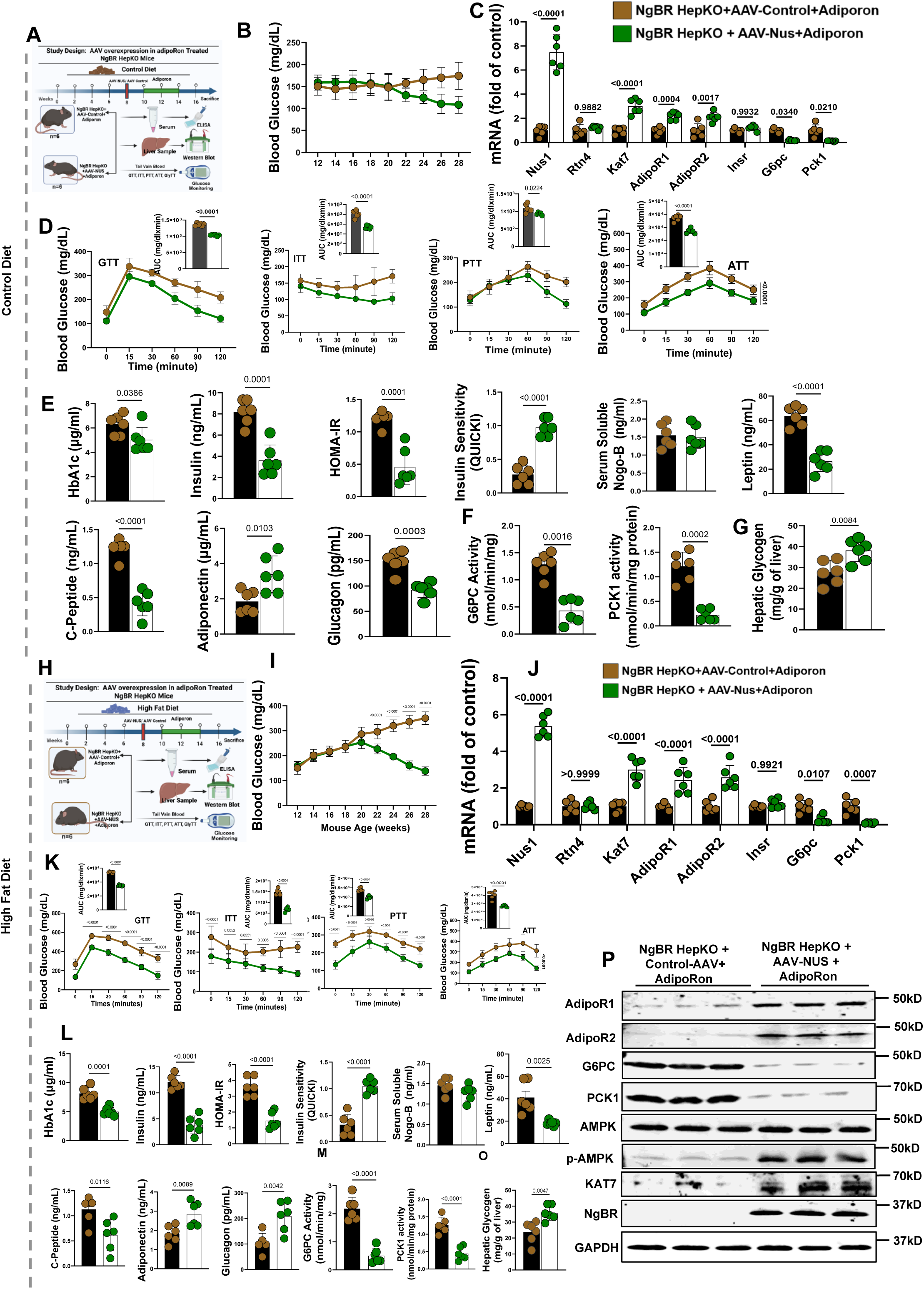
AAV-mediated hepatic NgBR restoration reinstates adiponectin responsiveness and glucose homeostasis in NgBR-deficient mice. **(A)** Schematic of AAV8-mediated hepatic NgBR (NUS1) re-expression in NgBR HepKO mice followed by AdipoRon treatment under control diet (CD) conditions. **(B)** Longitudinal fasting blood glucose measurements showing persistent hyperglycemia in AAV-control–treated NgBR HepKO mice and progressive normalization following AAV–NUS1 re-expression. **(C)** Hepatic qPCR analysis showing restoration of *Nus1* expression together with increased *KAT7*, *AdipoR1*, and *AdipoR2* transcripts and suppression of *G6pc* and *Pck1* in AAV–NUS1–treated mice (n = 5–6 mice per group; exact *P* values shown). **(D)** Dynamic metabolic testing under CD conditions (GTT, ITT, PTT, ATT) showing improved glucose tolerance, insulin sensitivity, and reduced substrate-driven glucose production following NgBR restoration. Insets show AUC quantification (n = 6–8 mice per group; exact *P* values shown). **(E–G)** Metabolic profiling under CD showing reductions in HbA1c, fasting insulin, C-peptide, and HOMA-IR, together with increased insulin sensitivity (QUICKI) and hepatic glycogen following AAV–NUS1 treatment. Additional parameters, including glucagon, leptin, adiponectin, and circulating soluble Nogo-B, are shown (n = 5–6 mice per group; exact *P* values shown). **(H)** Schematic of AAV–AdipoRon intervention under high-fat diet (HFD) conditions. **(I)** Longitudinal fasting glucose measurements under HFD showing persistent hyperglycemia in AAV-control–treated NgBR HepKO mice and improved glycemic control following AAV–NUS1 re-expression. **(J)** Hepatic qPCR analysis under HFD showing restoration of *Nus1*, *KAT7*, *AdipoR1*, and *AdipoR2* expression, with suppression of *G6pc* and *Pck1* (n = 5–6 mice per group; exact *P* values shown). **(K)** Dynamic metabolic testing under HFD (GTT, ITT, PTT, ATT) showing improved glucose tolerance, insulin responsiveness, and reduced gluconeogenic output following NgBR restoration. Insets show AUC quantification (n = 6–8 mice per group; exact *P* values shown). **(L–O)** Metabolic and hormonal parameters under HFD showing reductions in HbA1c, insulin, C-peptide, HOMA-IR, glucagon, leptin, and serum lipids, together with increased adiponectin and hepatic glycogen following AAV–NUS1 treatment. Enzymatic assays show reduced G6PC and PCK1 activity (n = 5–6 mice per group; exact *P* values shown). **(P)** Immunoblot analysis showing that NgBR re-expression restores AdipoR1, AdipoR2, and KAT7 protein levels, enhances AMPK phosphorylation in response to AdipoRon, and reduces G6PC and PCK1 protein abundance in NgBR HepKO mice. Data are presented as mean ± SD denotes biological replicates (mice) as indicated. Statistical tests are described in Methods. For time-course experiments (GTT, ITT, PTT, ATT), two-way repeated-measures ANOVA were used. Exact *P* values are provided in the figures.

Functional metabolic testing demonstrated near-complete rescue of glucose homeostasis. Under CD conditions, NgBR restoration normalized glucose excursions during glucose, insulin, pyruvate, and alanine tolerance tests (**Fig. 7D**). These effects were more pronounced under HFD, where NgBR re-expression reversed glucose intolerance, restored insulin sensitivity, and corrected substrate-driven gluconeogenesis (**Fig. 7K**). Consistent with these findings, systemic metabolic indices were markedly improved following NgBR restoration. HbA1c, fasting insulin, and C-peptide levels were reduced, insulin resistance (HOMA-IR) normalized, and insulin sensitivity (QUICKI) increased (**Fig. 7E, L**). Hormonal regulators of metabolism were also restored, with increased circulating adiponectin, reduced glucagon, stabilization of leptin levels, and recovery of hepatic glycogen stores (**Fig. 7E–G, 7L–O**). Circulating soluble Nogo-B levels remained unchanged, indicating that the observed effects are driven by hepatocyte-intrinsic NgBR restoration. At the level of hepatic glucose production, NgBR re-expression re-established suppression of gluconeogenic enzyme activity. Elevated G6PC and PCK1 activity in NgBR-deficient mice was significantly reduced following AAV-mediated restoration (**Fig. 7F, M**), indicating recovery of adiponectin-mediated metabolic control.

Mechanistically, NgBR restoration reinstated adiponectin signaling. AdipoRon-induced AMPK phosphorylation, which was absent in NgBR HepKO mice, was fully restored following NgBR re-expression (**Fig. 7P**). This was accompanied by normalization of AdipoR1 and AdipoR2 protein levels, restoration of KAT7 expression, and suppression of G6PC and PCK1 protein abundance, demonstrating recovery of both receptor expression and downstream signaling competence. Beyond hepatic signaling, NgBR restoration improved systemic metabolic parameters, including reductions in hepatic and circulating triglycerides, cholesterol, and free fatty acids, as well as improvement in markers of liver, pancreatic, and kidney function **(Extended Data Fig. 12)**. These changes occurred independently of alterations in energy balance, as food intake, body weight, and energy expenditure remained comparable between groups **(Extended Data Fig. 13)**.

These findings reveal that hepatic NgBR restoration reverses multiple metabolic defects associated with NgBR deficiency in this model. Re-expression of NgBR reinstates adiponectin receptor expression, restores KAT7-dependent signaling, reactivates AMPK, suppresses hepatic gluconeogenesis, and normalizes systemic glucose and lipid homeostasis, supporting a role for NgBR as a regulator of adiponectin responsiveness in vivo.

## Discussion

Adiponectin has long been viewed as a promising therapeutic axis in metabolic disease, yet clinical translation has been limited by inconsistent efficacy.^6,19–21^ Here, we identify hepatic NgBR as a regulator of adiponectin signaling competence and provide mechanistic evidence linking NgBR loss to hepatic adiponectin resistance. Our findings demonstrate that failure of adiponectin signaling arises not from ligand insufficiency but from loss of hepatocellular responsiveness.

Across multiple model systems, hepatic NgBR expression is reduced in metabolic disease, whereas ligand availability remains largely unchanged. Hepatocyte-specific NgBR deletion recapitulates hyperglycemia, dyslipidemia, and multi-organ dysfunction without alterations in energy balance, indicating a primary hepatic defect. Mechanistically, NgBR constrains gluconeogenesis by maintaining adiponectin receptor expression and downstream AMPK activation.^10,11^ Loss of NgBR induces a transcriptional program characterized by activation of gluconeogenic genes and suppression of metabolic signaling pathways. Consistent with this, pharmacological activation of adiponectin receptors reveals a strict dependence on NgBR: AdipoRon improves metabolic parameters in control mice but is ineffective in NgBR-deficient animals despite equivalent exposure.^21^ This uncoupling of ligand availability from signaling output supports a state of intrinsic adiponectin resistance and supports a role for NgBR as a regulator of hepatocellular signaling competence required for adiponectin action.

At the molecular level, NgBR sustains a KAT7-dependent chromatin program that supports transcription of *AdipoR1* and *AdipoR2*. Loss of NgBR reduces histone acetylation at these loci, leading to receptor deficiency and impaired AMPK activation. Restoration of KAT7 rescues receptor expression and downstream signaling, supporting a model in which adiponectin resistance can arise at the level of chromatin accessibility. In parallel, NgBR links adiponectin signaling to lipid metabolism. In control livers, adiponectin agonism is associated with reduced ceramide and sphingolipid species, whereas NgBR-deficient livers show persistent enrichment of these lipotoxic metabolites. Increased sensitivity to ceramide-induced gluconeogenesis and partial rescue by PP2A inhibition support a ceramide–PP2A–AMPK axis as a reinforcing mechanism of metabolic dysfunction in the absence of NgBR ^36,37,39,40^.

Importantly, this defect is reversible. AAV-mediated restoration of hepatic NgBR reinstates adiponectin responsiveness, normalizes AMPK signaling, suppresses gluconeogenesis, and corrects systemic metabolic abnormalities without affecting energy balance. These data reveal that NgBR contributes to hepatic metabolic homeostasis and supports the idea that hepatocellular signaling competence, rather than ligand availability, may limit adiponectin action.

In summary, hepatic NgBR functions as a regulator of adiponectin signaling by sustaining a KAT7-dependent epigenetic program that preserves receptor expression and metabolic control. Loss of NgBR is associated with impaired adiponectin signaling, ceramide accumulation, and excessive glucose production, whereas restoration of NgBR reverses these defects in the experimental systems examined here. Together, these findings support a model in which hepatic NgBR regulates adiponectin receptor competence through KAT7-dependent chromatin mechanisms and contributes to hepatic metabolic regulation in metabolic disease.

### Limitations of the study

Several limitations should be considered. Although our findings are supported across human datasets, non-human primates, and mouse models, the mechanistic experiments rely primarily on murine systems and hepatocyte-based assays; thus, direct validation in human liver tissue with functional perturbation will be important to confirm translational applicability. While AdipoRon is widely used as a pharmacologic adiponectin receptor agonist, it may not fully recapitulate endogenous adiponectin signaling dynamics, including receptor isoform specificity and tissue crosstalk. In addition, although we identify a KAT7-dependent chromatin mechanism regulating *AdipoR1* and *AdipoR2* expression, the upstream signals linking NgBR to KAT7 stability and activity remain incompletely understood. Our lipidomic analyses further implicate ceramide–PP2A signaling in reinforcing adiponectin resistance; however, causal dissection of this axis in vivo will require targeted genetic or pharmacologic approaches. Finally, while AAV-mediated NgBR restoration demonstrates reversibility of the phenotype, long-term safety, tissue specificity, and therapeutic feasibility of targeting this pathway in humans remain to be established.

## Resource availability

### Lead contact

Further information and request for resources and reagents should be directed to and will be fulfilled by lead contact, Qing Robert Miao.

#### Materials availability

All unique reagents and materials generated in this study are available from the corresponding author upon reasonable request and completion of a materials transfer agreement (MTA), where applicable. The hepatocyte-specific NgBR knockout (NgBR HepKO) mouse model (NgBR fl/fl crossed with Albumin-Cre) and AAV8-TBG-NgBR-HA constructs generated for this study are available upon request. AdipoRon and other commercially available reagents were obtained from the sources specified in the Key Resources Table.

Human liver tissue microarrays were purchased from Sekisui Xenotech. Publicly available transcriptomic datasets analyzed in this study are accessible through the Gene Expression Omnibus (GEO) under accession numbers GSE15653 and GSE188418. Additional human plasma lipidomic data were obtained from the All of Us Research Program under approved data use agreements.

RNA-seq data generated in this study have been deposited in the Gene Expression Omnibus (GEO) and will be made publicly available upon publication. No new proprietary software or custom algorithms were generated in this study.

### Data and code availability

This study utilized a range of open-source tools for data analysis; however, no new algorithms or tools were developed. Detailed information about the tools and their respective versions can be found in the Methods section.

## Acknowledgments

This work is primarily supported by R01DK132056, National Institute of Diabetes and Digestive and Kidney Diseases (NIDDK).

## Author contributions

M.S.M., M.B.T., W.H., and Q.R.M. conceived and designed the study. M.S.M., M.B.T., and W.H. performed experiments and analyzed the data. M.S.M. and M.B.T. conducted data analysis and interpretation.

R.B., M.C., G.J.S., and M.W.Z. contributed to experimental work and data collection.

Q.R.M. supervised the study. M.S.M. wrote the manuscript with input and revisions from all authors. All authors reviewed and approved the final manuscript.

## Declaration of interests

The authors declare that a patent application related to this work has been filed by their institution (inventors: Qing Robert Miao and Mohammad Sarif Mohiuddin; application pending).

## Declaration of Generative AI and AI-Assisted Technologies in the Writing Process

During the preparation of this manuscript, the authors used ChatGPT (OpenAI) and Grammarly (Grammarly, Inc.) to correct grammatical errors and improve clarity. These tools were used solely for editorial assistance. All content was carefully reviewed and revised by the authors, who take full responsibility for the accuracy and integrity of the final manuscript.

## METHODS

### Human Liver Samples

Human liver tissue microarrays (formalin-fixed, paraffin-embedded) were purchased from Sekisui Xenotech and included samples from lean individuals (**n = 6**), obese individuals without type 2 diabetes (**n = 15**), and obese individuals with type 2 diabetes (**n = 30**). All samples were provided as de-identified specimens without direct identifiers. Available clinical metadata, including BMI and diabetes status, were supplied by the vendor. Use of these commercially available, de-identified specimens was considered not human subjects research and was therefore exempt from institutional review board oversight. Immunohistochemical staining for NgBR was performed on the tissue microarrays, and staining intensity was quantified using a semi-quantitative H-score approach. All image acquisition and scoring were performed in a blind manner with respect to donor group assignment.

### Animals

All animal experiments were approved by the Institutional Animal Care and Use Committee (IACUC) of New York University Long Island School of Medicine (Mineola, NY, USA) and were conducted in accordance with NIH guidelines for the care and use of laboratory animals. Mice were housed in a pathogen-free facility under standard laboratory conditions, including a 12-hour light/12-hour dark cycle (lights on at 7:00 a.m.), a temperature-controlled environment (22–23 °C), and ad libitum access to food and water unless otherwise specified. Animals were acclimated to handling for at least one week prior to metabolic testing. C57BL/6J wild-type mice (JAX:000664) and db/db mice (JAX:000697) were obtained from The Jackson Laboratory. Hepatocyte-specific NgBR knockout mice (NgBR HepKO) were generated by crossing NgBR floxed mice with Albumin-Cre transgenic mice (JAX:003574) as described in our previous publication.^41^ Littermate NgBR floxed mice lacking Cre recombinase were used as controls (NgBR fl/fl). Only male mice were used for all experiments to minimize variability due to sex-dependent differences in metabolic phenotypes. Unless otherwise indicated, mice were 12 weeks of age at the start of dietary interventions. Animals were fed either a control diet (CD; D12450J, 10% kcal from fat) or a high-fat diet (HFD; D12492, 60% kcal from fat; Research Diets) for up to 16 weeks.

## METHOD DETAILS

### Study Design and Randomization

Mice were randomly assigned to experimental groups based on genotype and diet using simple randomization, with littermates distributed across groups whenever possible. Investigators were blinded to group allocation during metabolic phenotyping, biochemical analyses, histological assessments, and data quantification, and blinding was maintained until completion of data analysis. Sample sizes were determined based on prior experience with metabolic phenotyping studies and were sufficient to detect biologically meaningful differences in glucose homeostasis. Humane endpoints were defined in advance, and mice were euthanized at study completion by CO_₂_ inhalation followed by cervical dislocation in accordance with institutional guidelines. No animals were excluded from analysis unless predefined technical failure occurred.

### AdipoRon Administration

AdipoRon, a small-molecule adiponectin receptor agonist, was used to mimic adiponectin signaling. AdipoRon was prepared fresh on the day of injection and administered intraperitoneally (i.p.) at a dose of 50 mg/kg/day. Treatment was initiated after 10 weeks of Control diet (CD) or high-fat diet (HFD) feeding and continued once daily for 4 consecutive weeks. Vehicle-treated mice received intraperitoneal injections on the same schedule. In AAV rescue experiments, AdipoRon treatment was initiated after hepatic viral expression had stabilized. All metabolic phenotyping was performed after completion of the AdipoRon treatment period.

### AAV-Mediated Hepatic NgBR Rescue

Recombinant AAV8 vectors expressing HA-tagged NgBR under the hepatocyte-specific thyroxine-binding globulin (TBG) promoter (AAV8-TBG-NgBR-HA) or empty control vectors were used. Adult mice received a single tail vein injection of AAV at a dose of 0.5 × 10¹¹ genome copies (gc) per mouse following 8 weeks of control diet or high-fat diet feeding. Viral expression was allowed to proceed for 6 weeks prior to metabolic phenotyping. Hepatic specificity and expression efficiency were verified by quantitative PCR and immunoblotting of liver tissue.

### Longitudinal Metabolic Monitoring

Body weight, food intake, and fasting blood glucose levels were measured biweekly throughout the experimental period. Fasting blood glucose was assessed after a 12-hour overnight fast using tail vein sampling and a handheld glucometer.

### Metabolic Tolerance Tests

All metabolic tolerance tests were conducted after a 12-hour overnight fast (food removed at 7:00 p.m.; testing performed the following morning). Mice were allowed free access to water during fasting.

### Glucose Tolerance Test (GTT)

Following a 12-hour overnight fast, D-glucose was dissolved in sterile saline and administered intraperitoneally at a dose of 2 g/kg body weight. Blood glucose was measured from the tail vein at 0, 15, 30, 60, 90, and 120-minutes post-injection.

### Insulin Tolerance Test (ITT)

Following a 12-hour overnight fast, recombinant human insulin was administered intraperitoneally at a dose of 0.75 U/kg body weight. Blood glucose was measured from the tail vein at 0, 15, 30, 60, 90, and 120-minute post-injection.

### Pyruvate Tolerance Test (PTT)

Following a 12-hour overnight fast, sodium pyruvate was administered intraperitoneally at a dose of 2 g/kg body weight. Blood glucose was measured from the tail vein at 0, 15, 30, 60, 90, and 120 minutes post-injection.

### Alanine Tolerance Test (ATT)

Following a 12-hour overnight fast, L-alanine was administered intraperitoneally at a dose of 1.5 g/kg body weight. Blood glucose was measured from the tail vein at 0, 15, 30, 60, 90, and 120 minutes post-injection.

For all tolerance tests, area under the curve (AUC) was calculated using the trapezoidal method.

### DEXA Body Composition Analysis

Whole-body composition was assessed by dual-energy X-ray absorptiometry (DEXA). All body composition and metabolic measurements were performed at ambient room temperature (23 °C). Mice were anesthetized and maintained under isoflurane inhalation anesthesia throughout the scanning procedure to minimize movement. Anesthetized mice were positioned prone on the specimen tray and scanned for approximately 4–5 min per animal. The head region was excluded from analysis.

Parameters assessed included total body weight, fat mass, lean mass, percent body fat, bone mineral density (BMD), and bone mineral content (BMC). Animals were returned to their home cages and monitored during recovery from anesthesia.

### CLAMS Metabolic Cage Analysis

Locomotor activity, oxygen consumption, and carbon dioxide production were monitored using a Comprehensive Lab Animal Monitoring System (Oxymax®-CLAMS, Columbus Instruments, Columbus, OH). Mice were singly housed in metabolic cages and acclimated for 48 hours prior to data acquisition to allow adaptation to the cages and drinking system. Data collected during the initial acclimation period were excluded from analysis. Following acclimation, energy expenditure, food intake, locomotor activity, oxygen consumption (VO_₂_), carbon dioxide production (VCO_₂_), and respiratory exchange ratio (RER) were continuously recorded. Measurements were analyzed over total 24-hour periods as well as separately for light and dark cycles.

### Blood Biochemistry and Hormone Measurements

Blood samples were collected from fasted mice in the morning and centrifuged at 8,000 × g for 5 minutes to obtain serum. Serum samples were analyzed for HbA1c, insulin, C-peptide, adiponectin, glucagon, leptin, free fatty acids, triglycerides, cholesterol, alanine aminotransferase (ALT), aspartate aminotransferase (AST), alkaline phosphatase (ALP), total bilirubin, albumin, blood urea nitrogen (BUN), creatinine, amylase, and lipase using commercially available ELISA or colorimetric assay kits according to the manufacturers’ instructions. All the kit details are available on the **Key Resources table.**

Insulin resistance and insulin sensitivity were estimated using the homeostatic model assessment of insulin resistance (HOMA-IR) and the quantitative insulin sensitivity check index (QUICKI), calculated from fasting glucose and insulin values.

### Primary Hepatocyte Isolation

Mice were anesthetized and euthanized by cervical dislocation in accordance with institutional guidelines. The inferior vena cava was cannulated using a 24 G butterfly needle connected to a peristaltic pump. A small incision was made in the portal vein to allow outflow, and the liver was perfused in situ with 35 mL phosphate-buffered saline (PBS) containing 1% penicillin/streptomycin to remove blood. The liver was subsequently perfused with 35 mL liver perfusion medium (Gibco #17701038) supplemented with 1% penicillin/streptomycin, followed by 35 mL liver digestion medium (Gibco, #17703034) containing 0.25% collagenase II (Thermo Fisher Scientific #17101015). Following perfusion, the liver was excised, minced in digestion medium, and incubated on a rotor at 37 °C until complete tissue dissociation was achieved. An equal volume of hepatocyte culture medium (Gibco #17705-021) containing 10% fetal bovine serum (FBS) was added to terminate collagenase activity. The cell suspension was centrifuged at 300 × g for 10 minutes, and the resulting pellet was washed with hepatocyte wash buffer and centrifuged again at 300 × g for 10 minutes. The pellet was resuspended in hepatocyte culture medium and filtered through a 100 μm sterile cell strainer (Fisher Scientific, #431752). The filtrate was centrifuged at 300 × g for 10 minutes, and the final hepatocyte pellet was resuspended in hepatocyte culture medium and seeded onto collagen-coated six-well plates (Corning, #356400).

### Hepatic Glucose Production Assay

Primary hepatocytes were incubated in glucose-free DMEM supplemented with lactate (20 mM) and pyruvate (2 mM). Cells were maintained under serum-free conditions for the indicated incubation period. At the end of the incubation, culture supernatants were collected, and glucose concentrations were quantified using an enzymatic glucose assay according to the manufacturer’s instructions. Glucose production was normalized to total cellular protein content, determined from parallel cell lysates.

### si-RNA-Mediated Knockdown in HepG2 Cells

HepG2 cells were cultured in Dulbecco’s modified Eagle’s medium (DMEM) supplemented with 10% fetal bovine serum and penicillin/streptomycin and maintained at 37 °C in a humidified incubator with 5% CO_₂_. Cells were seeded one day prior to transfection to achieve approximately 50–70% confluence at the time of siRNA delivery. For siRNA-mediated gene silencing, HepG2 cells were transfected using Lipofectamine according to the manufacturer’s instructions. siRNA duplexes were used at a final concentration of 20 nM. siRNA–Lipofectamine complexes were prepared by diluting siRNA and Lipofectamine separately in Opti-MEM, combining the solutions, and incubating for 20 minutes at room temperature prior to addition to cells. Complexes were added dropwise to cells maintained in serum-containing medium, and plates were gently rocked to ensure even distribution. Control conditions included non-targeting siRNA and transfection reagent–only (mock) controls. Culture medium was replaced with fresh complete medium 6–8 hours post-transfection or the following day if signs of cytotoxicity were observed. Gene silencing efficiency was assessed at the mRNA level 24 hours post-transfection by quantitative real-time PCR and at the protein level 48 hours post-transfection by immunoblotting. Functional assays were performed at time points corresponding to maximal knockdown, as shown in the figure legends.

### si-RNA-Mediated Knockdown in Human Hepatocyte

Cryopreserved human hepatocytes were thawed and plated according to the manufacturer’s instructions in hepatocyte medium and maintained at 37 °C in a humidified incubator with 5% CO_₂_. Cells were seeded onto poly-L-lysine–coated culture plates at the recommended density and allowed to attach for 16–24 hours prior to transfection. Medium was refreshed the following day to remove residual DMSO and unattached cells. For gene silencing, hepatocytes were transfected with gene-specific siRNA duplexes at a final concentration of 20 nM using Lipofectamine RNAiMAX according to the manufacturer’s protocol optimized for primary cells. Briefly, siRNA and Lipofectamine RNAiMAX were diluted separately in Opti-MEM, combined, and incubated for 15–20 minutes at room temperature to allow complex formation. The siRNA–lipid complexes were added dropwise to cells maintained in serum-containing medium. Non-targeting siRNA was used as a negative control. Culture medium was replaced with fresh hepatocyte medium 6–8 hours after transfection to minimize cytotoxicity. Knockdown efficiency was assessed at the mRNA level 24 hours post-transfection by quantitative real-time PCR and at the protein level 48 hours post-transfection by immunoblotting. Functional assays were performed at time points corresponding to maximal gene suppression, as indicated in the figure legends.

### Lentiviral Overexpression

Lentiviral vectors expressing KAT7 or a control lentivirus lacking a transgene were generated using a second-generation lentiviral packaging system. Lentiviral expression plasmids were co-transfected with the packaging plasmid psPAX2 and the envelope plasmid pVSVG into Lenti-X 293T cells using Lipofectamine 3000, according to the manufacturer’s instructions. Twelve hours after transfection, the culture medium was replaced with high–fetal bovine serum (FBS)-containing medium without antibiotics. Lentiviral supernatants were collected at 24- and 48-hours post-transfection, centrifuged at 12,000 rpm for 5 minutes to remove cellular debris, and filtered through 0.45-µm filters. Filtered viral supernatants were concentrated using the Lenti-X Concentrator according to the manufacturer’s protocol. HepG2 cells were infected with the concentrated lentiviral preparations. Twenty-four hours after infection, cells were selected with puromycin (1 µg/mL) for 3–4 days. Successfully transduced cell populations were subsequently expanded and allowed to recover to confluency prior to downstream analyses.

### Western Blotting

Cellular protein was extracted using radioimmunoprecipitation assay (RIPA) lysis buffer (Thermo Scientific) supplemented with protease and phosphatase inhibitor tablets (Pierce–Thermo Scientific). Lysates were cleared by centrifugation, and protein concentrations were determined using a detergent-compatible colorimetric protein assay (Bio-Rad, CA). Equal amounts of protein were mixed with Laemmli sample buffer supplemented with 10% β-mercaptoethanol, denatured by heating at 65 °C for 10 minutes, and resolved on 8–12% or 10% SDS–PAGE gels as indicated. Proteins were transferred onto nitrocellulose membranes using a Bio-Rad Trans-Blot® Turbo™ Transfer System for 30 minutes. Membranes were blocked in Tris-buffered saline (TBS) containing 1% (wt/vol) casein buffer (Bio-Rad) and incubated with primary antibodies overnight at 4 °C. Membranes were washed with TBS containing 0.1% Tween-20 (TBST) prior to incubation with secondary antibodies. Primary antibodies were detected using highly cross-adsorbed IRDye-conjugated secondary antibodies. All the details are available on the key **resources table.** Blots were imaged wet using the Odyssey CLx Imaging System (LI-COR Biosciences) with detection in the 680-nm and 780-nm channels. Densitometric analysis was performed using Image Studio software (LI-COR Biosciences).

### Quantitative Real-Time PCR

Total RNA was isolated from cells or tissues using TRIzol reagent or the RNeasy Mini Kit (QIAGEN, Hilden, Germany), according to the manufacturers’ instructions. RNA concentration and purity were assessed spectrophotometrically. One microgram of total RNA was reverse transcribed into complementary DNA (cDNA) using the iScript cDNA Synthesis Kit (Bio-Rad, Los Angeles, CA). Quantitative real-time PCR (qPCR) was performed on a Bio-Rad CFX Real-Time PCR Detection System using iTaq Universal SYBR Green Supermix (Bio-Rad, Los Angeles, CA). All reactions were run under standard cycling conditions as recommended by the manufacturer. Relative messenger RNA (mRNA) expression levels were calculated using the ΔΔCt method and normalized to the housekeeping gene GAPDH. Primer sequences used for qPCR are provided in **(Supplementary Table 1).**

### Chromatin Immunoprecipitation–qPCR (ChIP–qPCR) Assay

Chromatin immunoprecipitation (ChIP) assays were performed in HepG2 cells under the indicated treatment conditions using the SimpleChIP® Plus Enzymatic Chromatin IP Kits (Cell Signaling Technology, #9004 and #9005), according to the manufacturer’s instructions. Briefly, cells were cross-linked with formaldehyde, quenched with glycine, and nuclei were isolated. chromatin was digested enzymatically and sheared to an average fragment size suitable for qPCR analysis. Immunoprecipitation was carried out using antibodies specific to the indicated targets, as listed in **key resources table.** An anti-histone H3 antibody was used as a positive control, and a normal IgG antibody was used as negative control. Following immunoprecipitation, cross-links were reversed, and DNA was purified according to the kit protocol. Recovered DNA was analyzed by quantitative PCR (qPCR) using gene promoter–specific primers, which are listed in **Supplementary Table 1.** ChIP–qPCR data were calculated as percent input or fold enrichment relative to IgG controls, as indicated in the figure legends.

### Histology, Immunohistochemistry

Liver tissues were fixed in 10% neutral-buffered formalin, embedded in paraffin, and sectioned at 5 μm thickness. Sections were stained with hematoxylin and eosin (H&E) following standard histological procedures. For immunohistochemistry (IHC) paraffin sections were deparaffinized, rehydrated, and subjected to heat-induced antigen retrieval. Endogenous peroxidase activity was quenched for IHC analyses, and nonspecific binding was blocked prior to antibody incubation. Sections were incubated with validated primary antibodies against NgBR, G6PC, PCK1 overnight at 4 °C, followed by appropriate secondary antibodies for chromogenic detection (IHC). Images were acquired using a fluorescence or bright-field microscope under identical exposure settings. Quantification, where applicable, was performed in a blinded manner using standardized image analysis criteria.

### RNA sequencing and Library preparation

The amount and quality of the total isolated RNA from the liver tissue were evaluated. To guarantee high-quality RNA for sequencing, RNA integrity numbers (RIN) were calculated. After that, the mRNA was separated from the total RNA using magnetic beads coupled to poly-T oligos, and it was broken into short fragments using divalent cations at a high temperature. Reverse transcriptase and random hexamer primers were used for first-strand cDNA synthesis, whereas DNA Polymerase I and RNase H were used for second-strand cDNA synthesis. To maintain strand orientation for directional libraries, dUTP was added to the second-strand synthesis. After that, the cDNA was subjected to adapter ligation, A-tailing, and end repair. To choose cDNA fragments of the ideal lengths for sequencing, size selection was used. A Qubit Fluorometer and an Agilent Bioanalyzer were used to assess the quality of the prepared libraries. Real-time PCR was used to quantify the libraries, and the Bioanalyzer was used to verify their size distribution. Libraries were pooled based on their effective concentration and data amount and sequenced on an Illumina platform using a paired-end 150 bp (PE150) strategy (Parkhomchuk et al. 2009). All these procedures were carried out by Novogene in Sacramento, CA.

### Data Processing

Raw sequencing reads were initially processed to eliminate adapters, poly-N sequences, and low-quality reads using Trimmomatic v0.36, applying the default parameters ^42^. Subsequently, each sample’s quality was evaluated with standard parameters implemented in FastQC program. The paired-end alignment of pre-processed reads to the reference mouse genome (mm10) was conducted using the HISAT2 v.2.1.0 splice-mapping technique with default parameters^43^. Furthermore, FeatureCounts v1.5.0-p3 was used to determine the number of reads mapped to each gene ^44^. Each gene’s FPKM (Fragments Per Kilobase of Transcript Sequence per Million Base Pairs) was determined using the gene’s length and the read count mapped to it. The most often used variables for estimating gene expression levels are read count and FPKM.

### Differential expression analysis

Differential expression analysis of any two groups with three biological replicates per group was performed using DESeq2 (v.1.37.4) implemented in Bioconductor, R package (v.4.2.1). DESeq2 examines the dependence of variance on mean counts in the count data from two sample groups and tests for differential gene expression using the negative binomial distribution.^45^ It computes statistical metrics such as False Discovery Rate (FDR), log2 fold change, and p-value following Benjamini and Hochberg’s technique. Genes having an adjusted p-value (padj) less than 0.05 were considered significantly expressed. The differentially expressed genes (DEGs) were visualized through complex heat maps and volcano plots.

### KEGG Pathway analysis

The KEGG (Kyoto Encyclopedia of Gene and Genome) is a database for higher-level of metabolic functions of biological systems and their annotations based on large-scale molecular datasets generated by high-throughput sequencing.^46,47^ KEGG pathway enrichment analysis of differentially expressed genes was conducted using clusterProfiler (v. 4.5.1) implemented in Bioconductor, R package., with the p-value (< 0.05) is considered as significant.^48^

### Lipidomics Analysis

Hepatic lipidomic profiling was performed by MetWareBio (8A Henshaw Street, Woburn, MA 01801). Fresh-frozen liver tissues were submitted directly to MetWareBio, where all subsequent procedures—including lipid extraction, mass spectrometry–based lipidomics acquisition, lipid identification, and primary quantification—were conducted by the service provider using their standardized platforms. Lipid species were annotated and grouped into major lipid classes, including fatty acids, glycerolipids, glycerophospholipids, sphingolipids, prenol lipids, and sterol lipids. The processed lipidomics datasets provided by MetWareBio were subsequently analyzed in our laboratory. Quantitative data were normalized to tissue weight prior to downstream analysis. Differential lipid abundance between experimental groups was determined based on log_₂_ fold-change thresholds with statistical significance assessed using adjusted p-values to control for multiple testing.

### Quantification and Statistical Analysis

All data are presented as mean ± SD, unless otherwise indicated. Sample size (n) refers to biological replicates and is defined in the corresponding figure legends. No statistical methods were used to predetermine sample size. Statistical analyses were performed using GraphPad Prism (v10.0). For comparisons between two groups, unpaired two-tailed Student’s *T*-tests were used. For comparisons involving more than two groups, one-way or two-way ANOVA followed by Tukey’s multiple comparisons test was applied, as appropriate. For repeated measurements (e.g., GTT, ITT, PTT, ATT time-course data), two-way repeated measures ANOVA was used. RNA-seq differential expression analysis was performed using DESeq2, with significance defined as adjusted *P* < 0.05 (Benjamini–Hochberg correction). Differences were considered statistically significant at *P* < 0.05. Exact *P* values are reported where appropriate.

## KEY RESOURCES TABLE

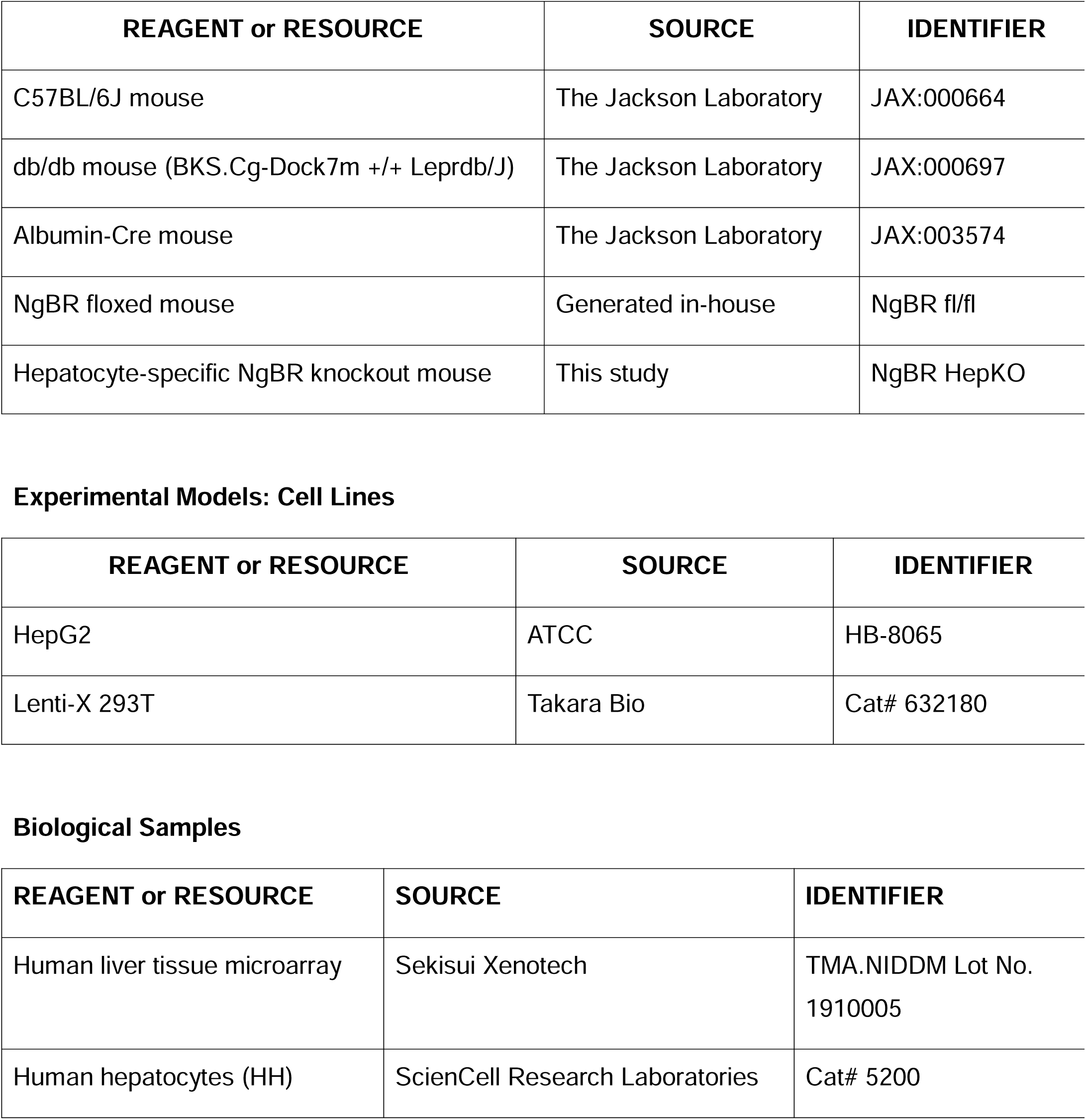
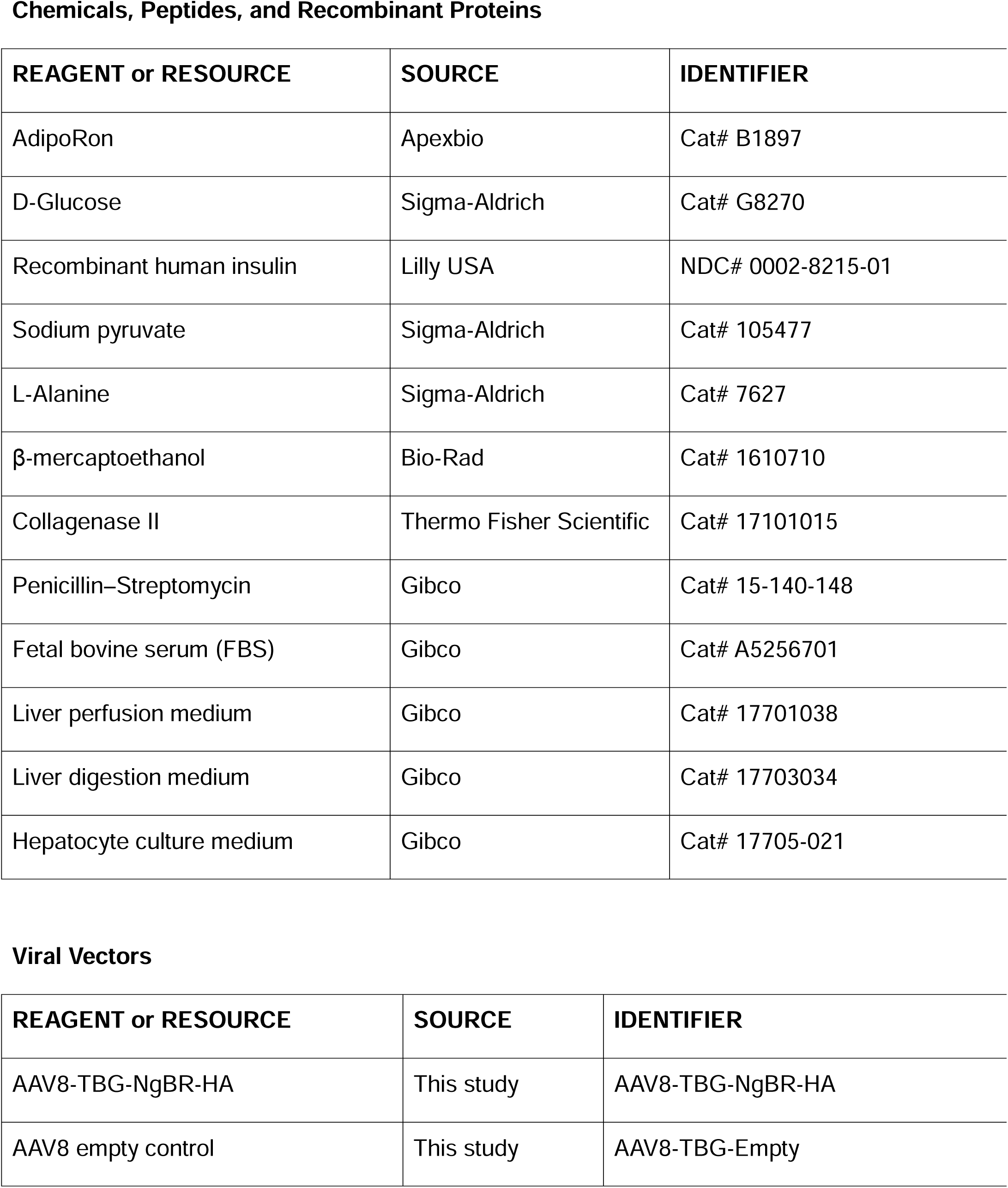
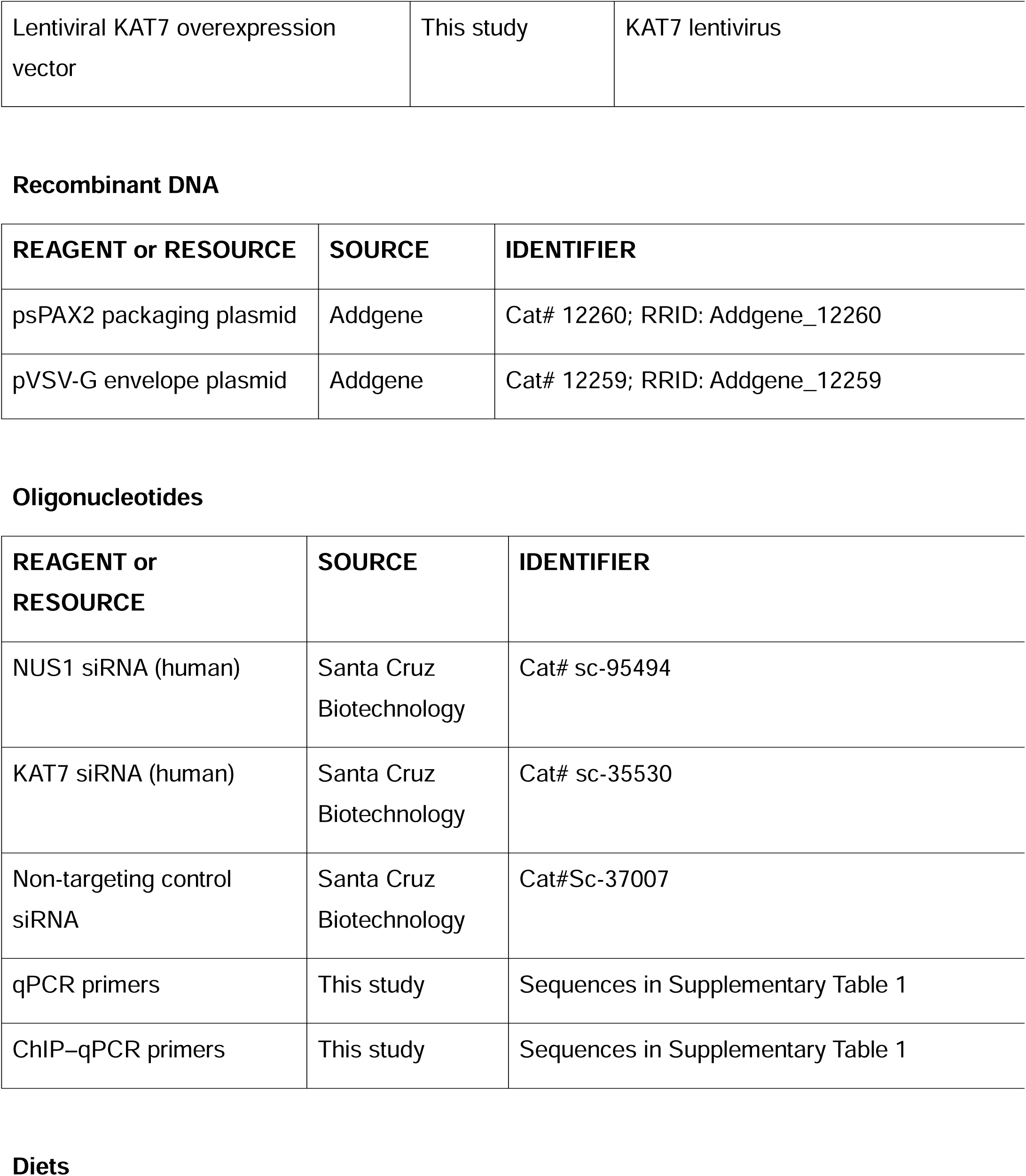
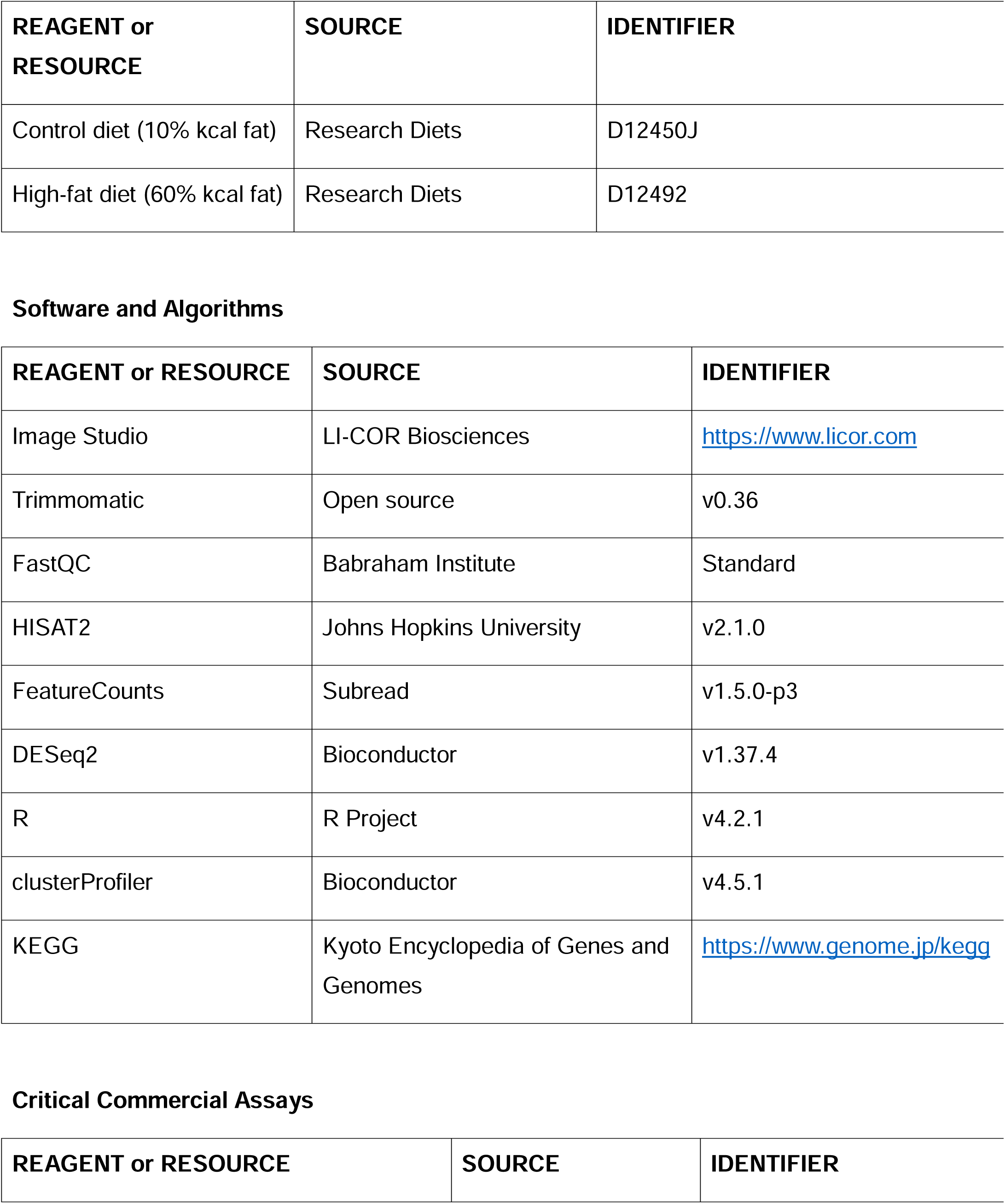
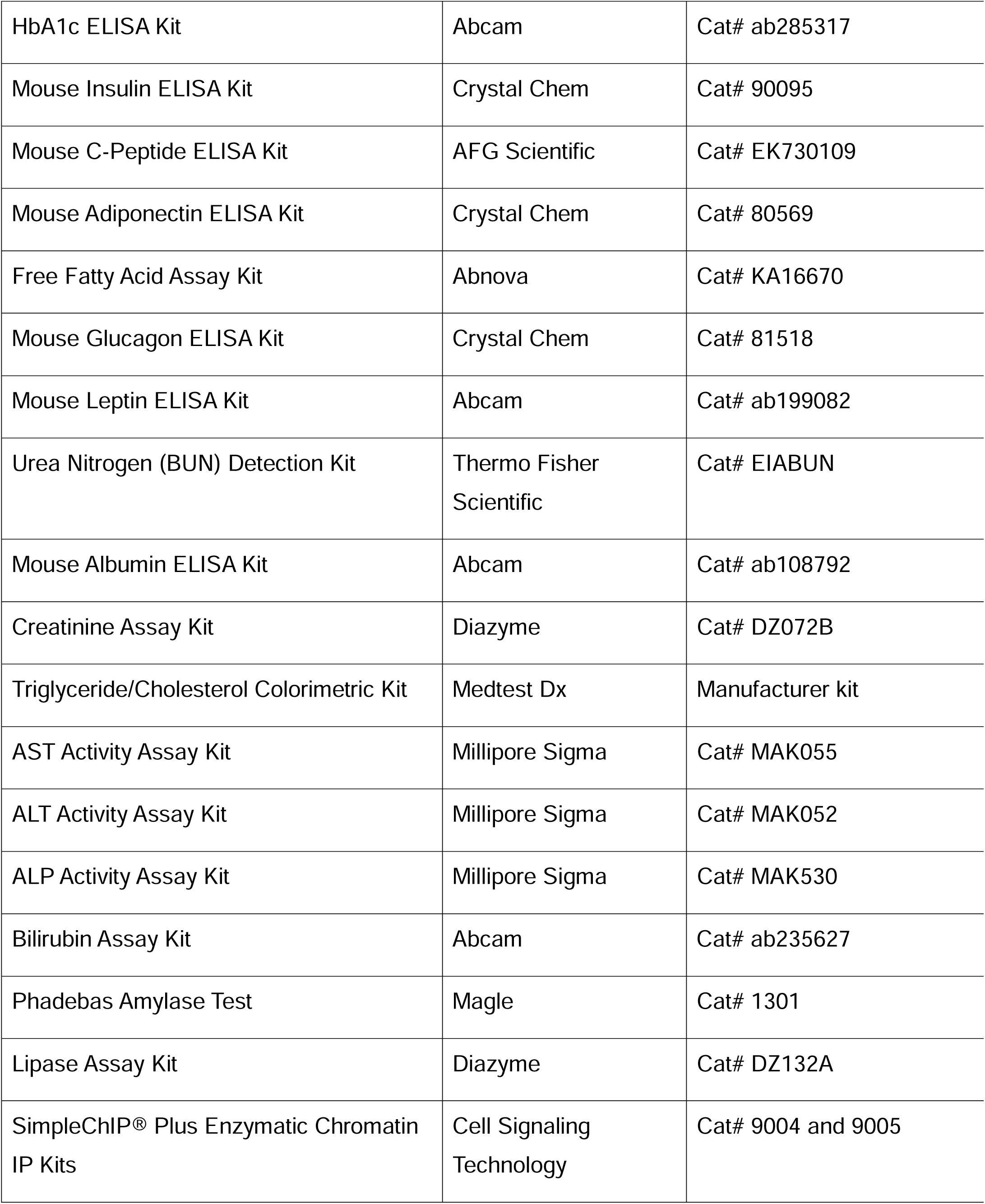
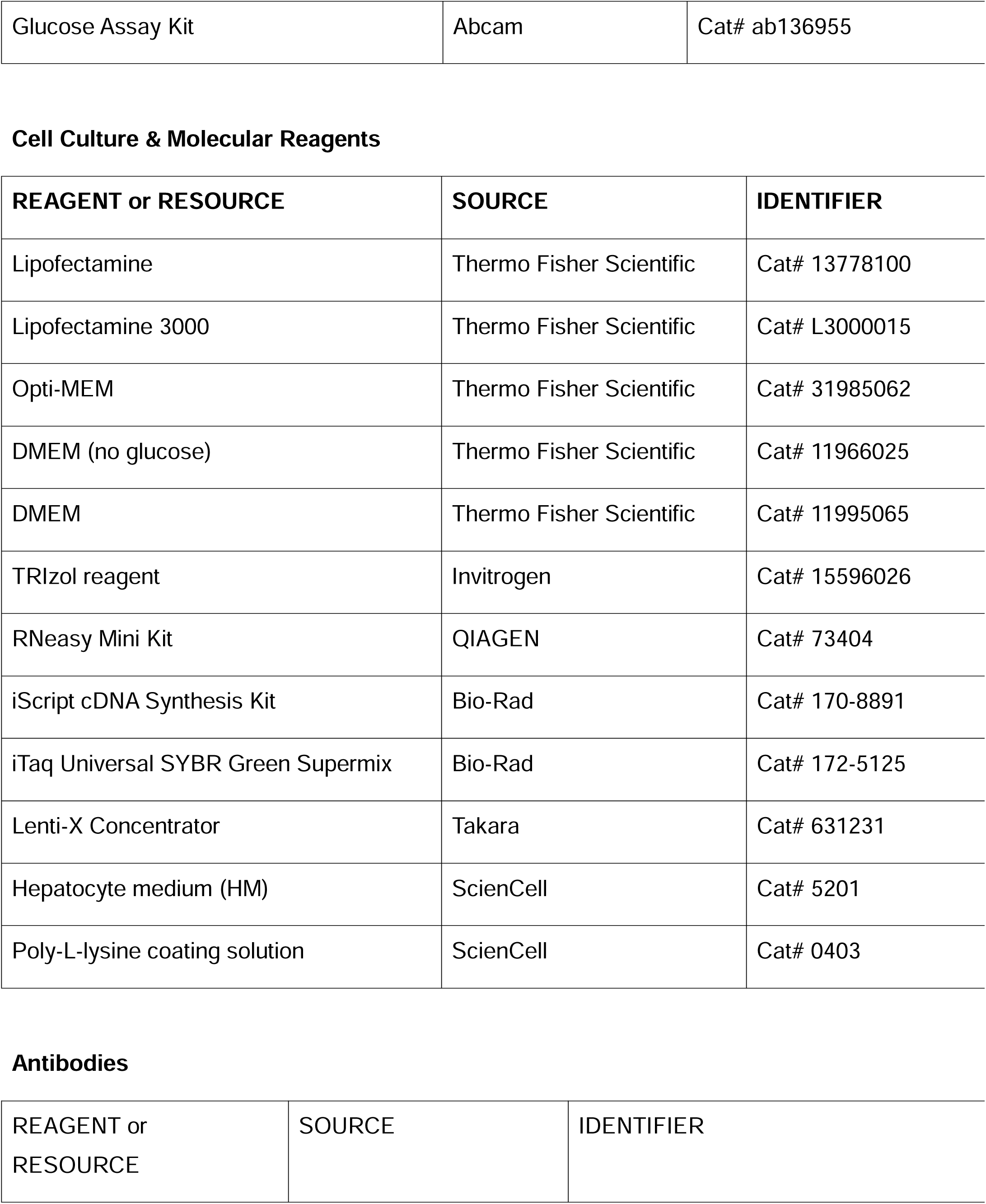
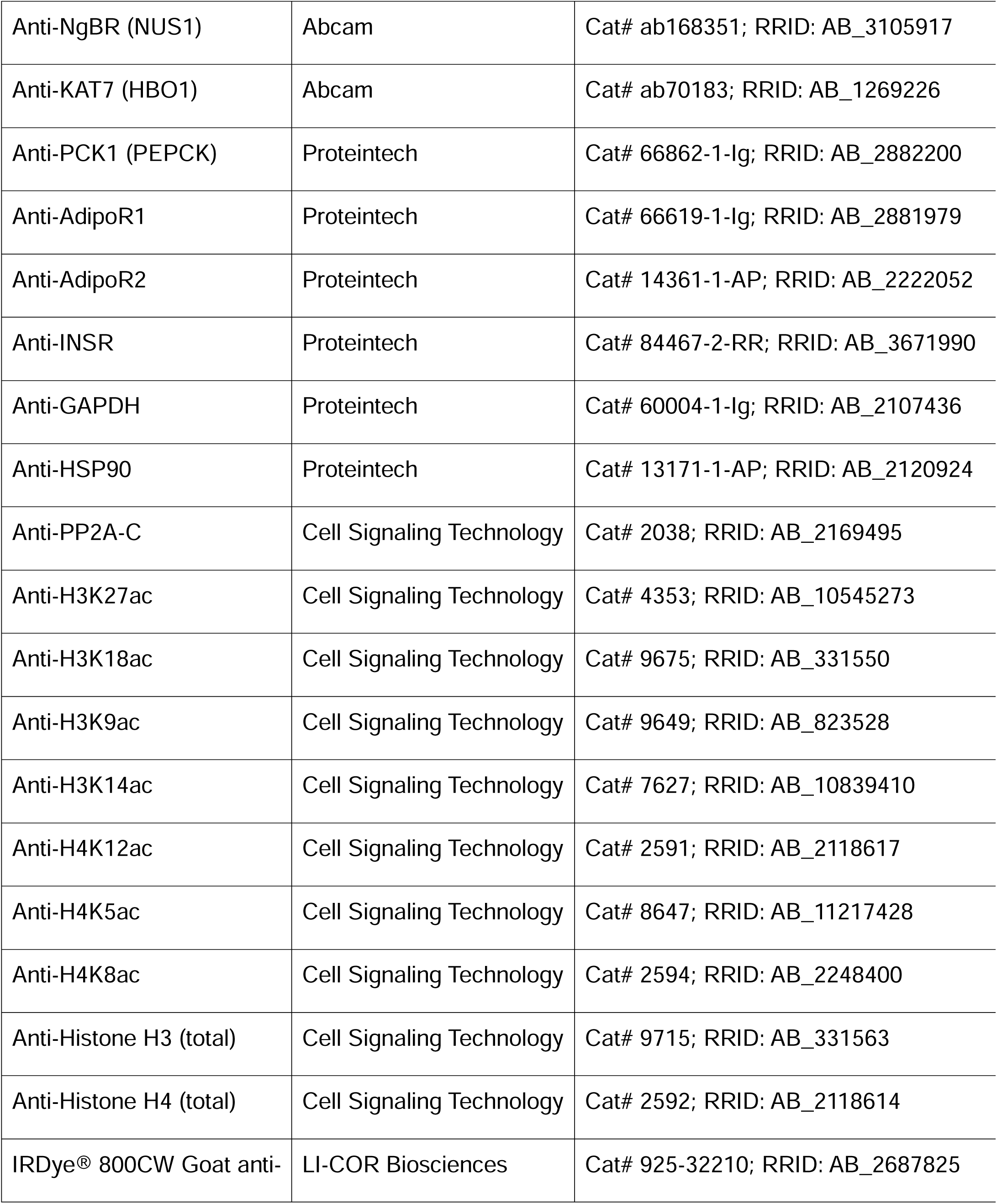
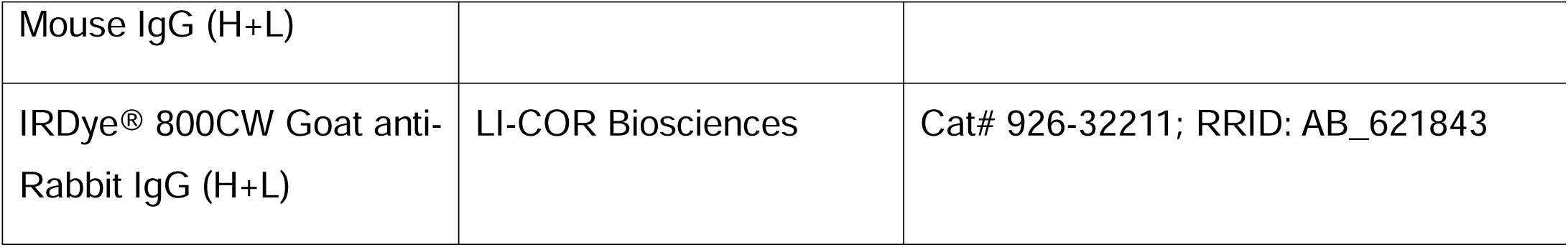

**Supplementary Table 1.**
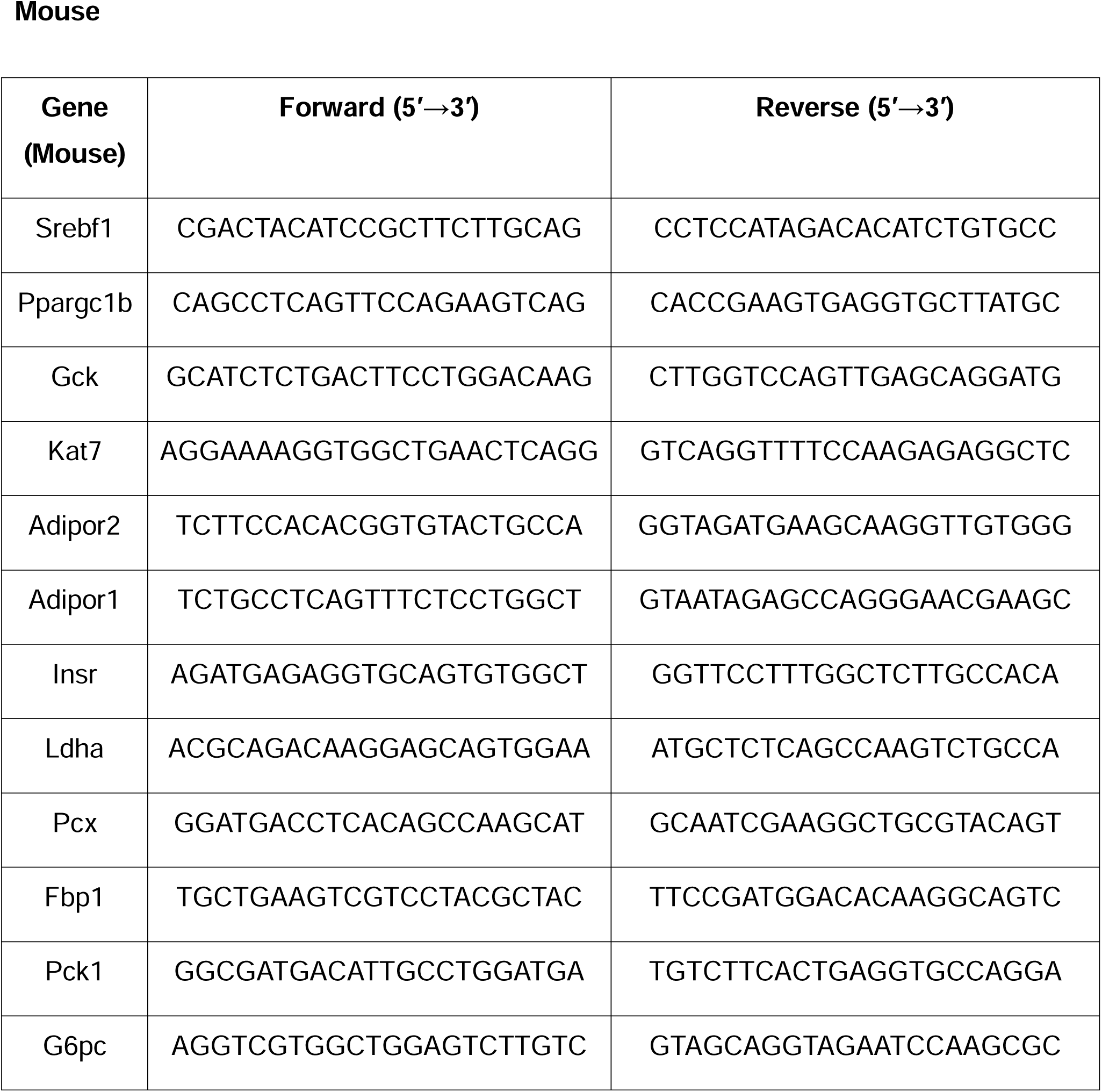
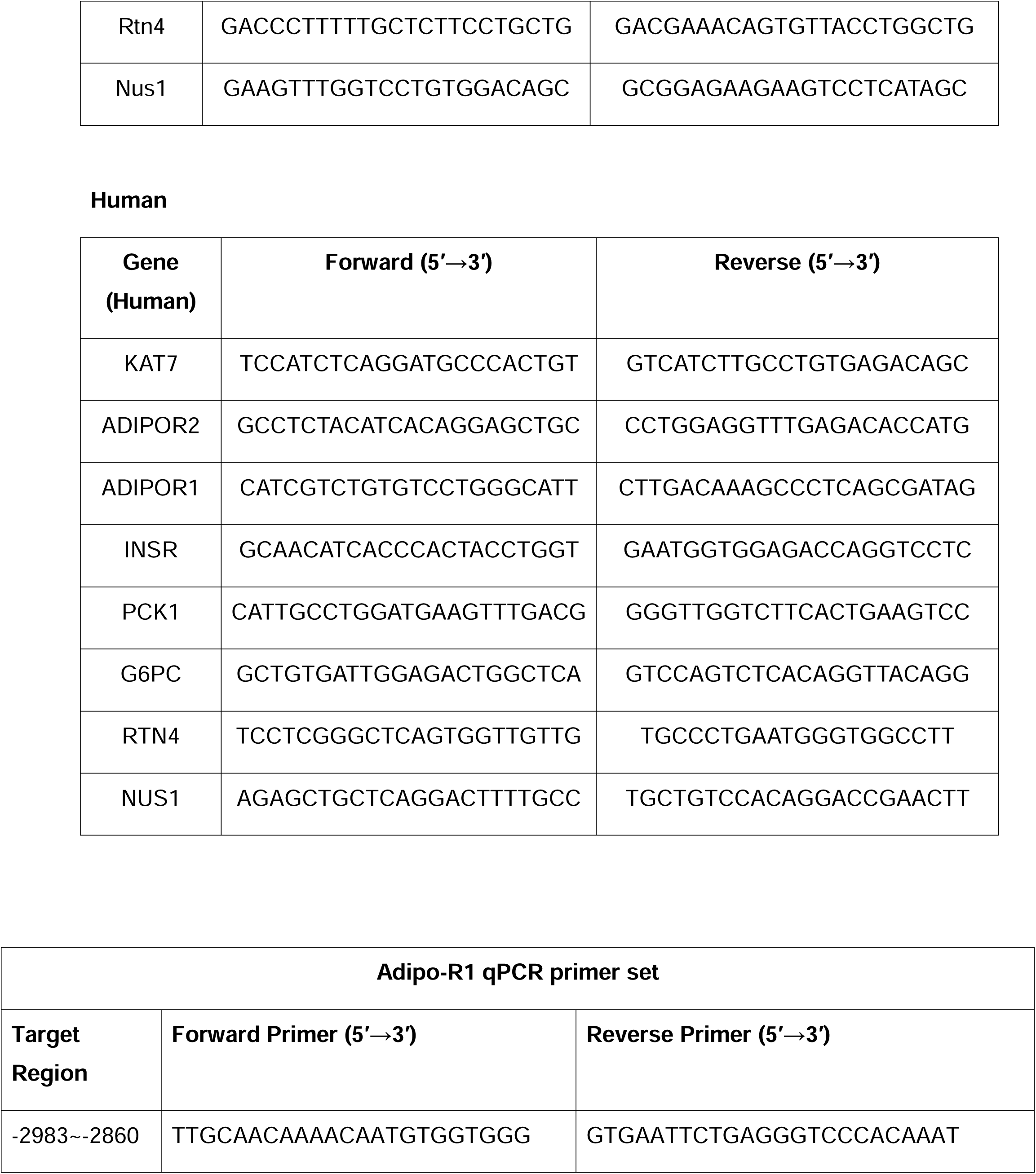
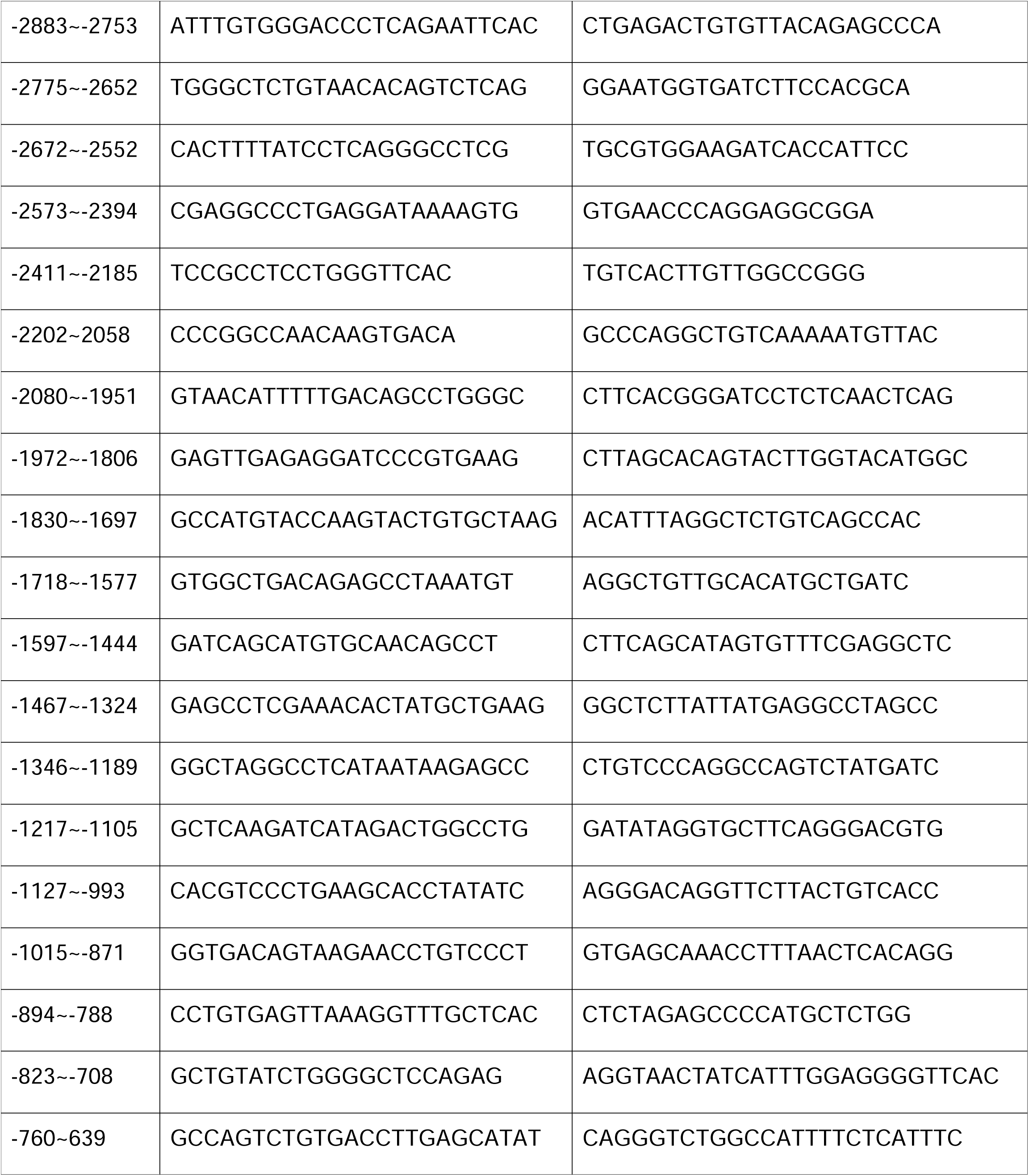
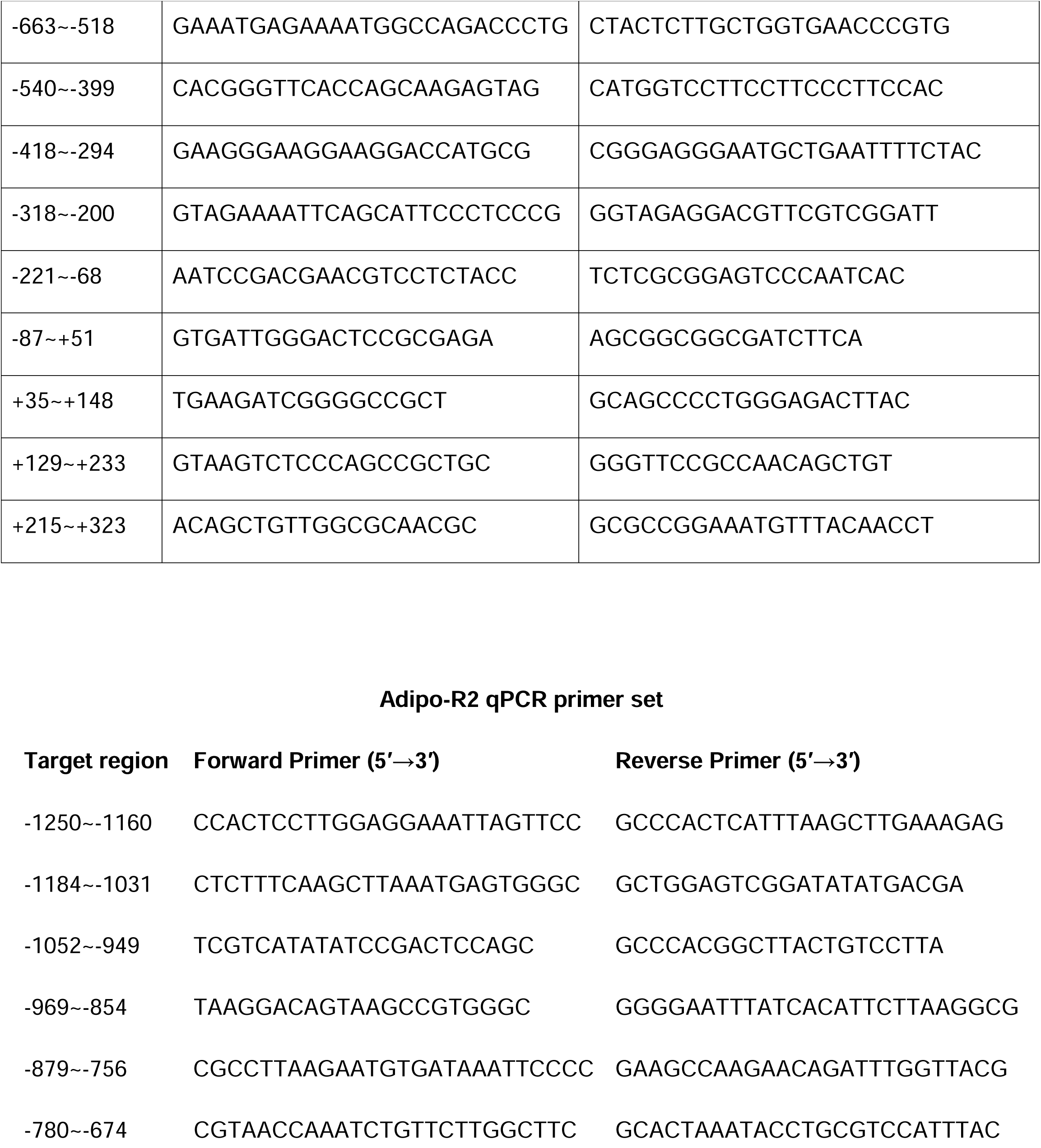
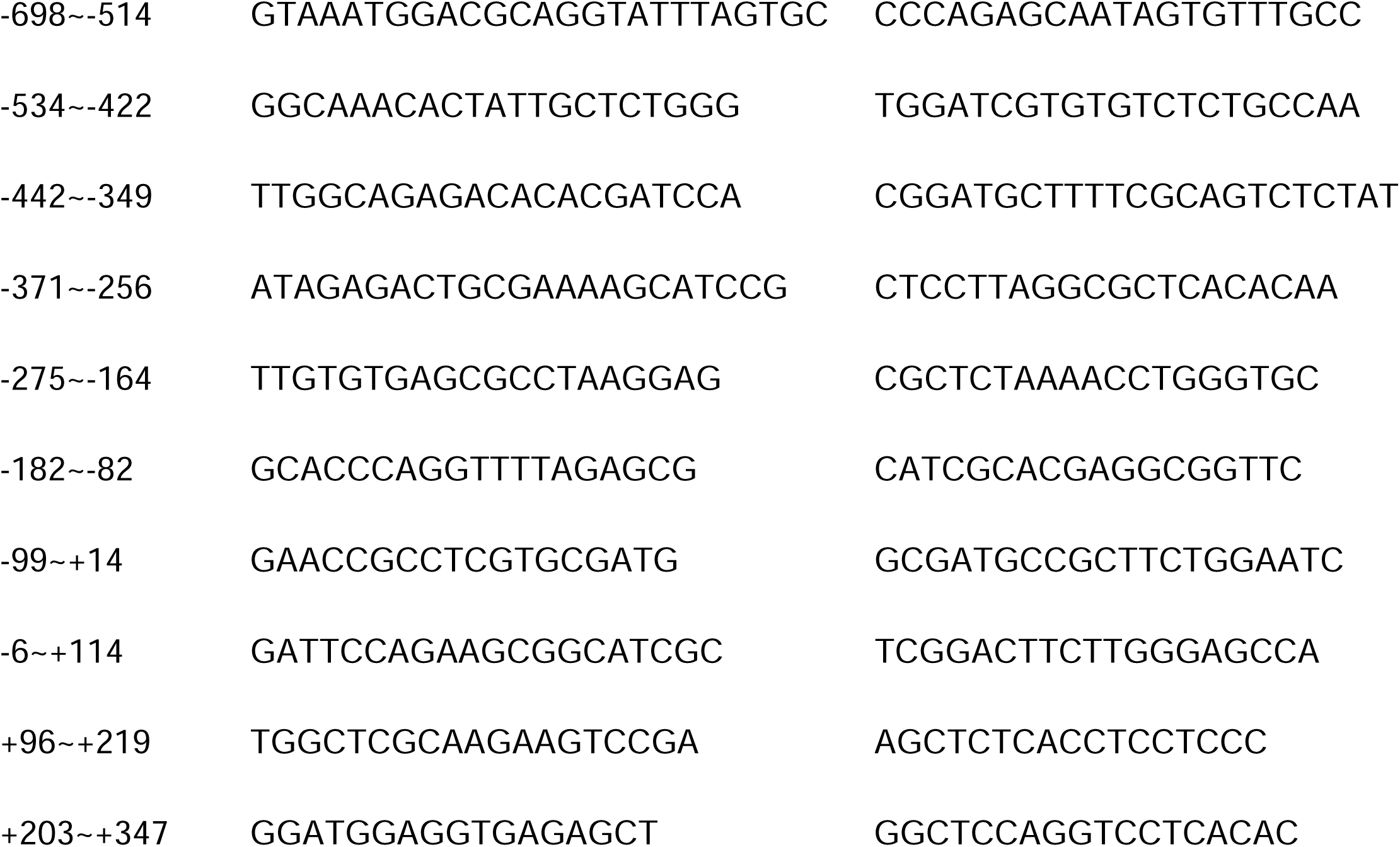
qPCR Primers Used in This Study.

**Extended Data Fig. 1:**
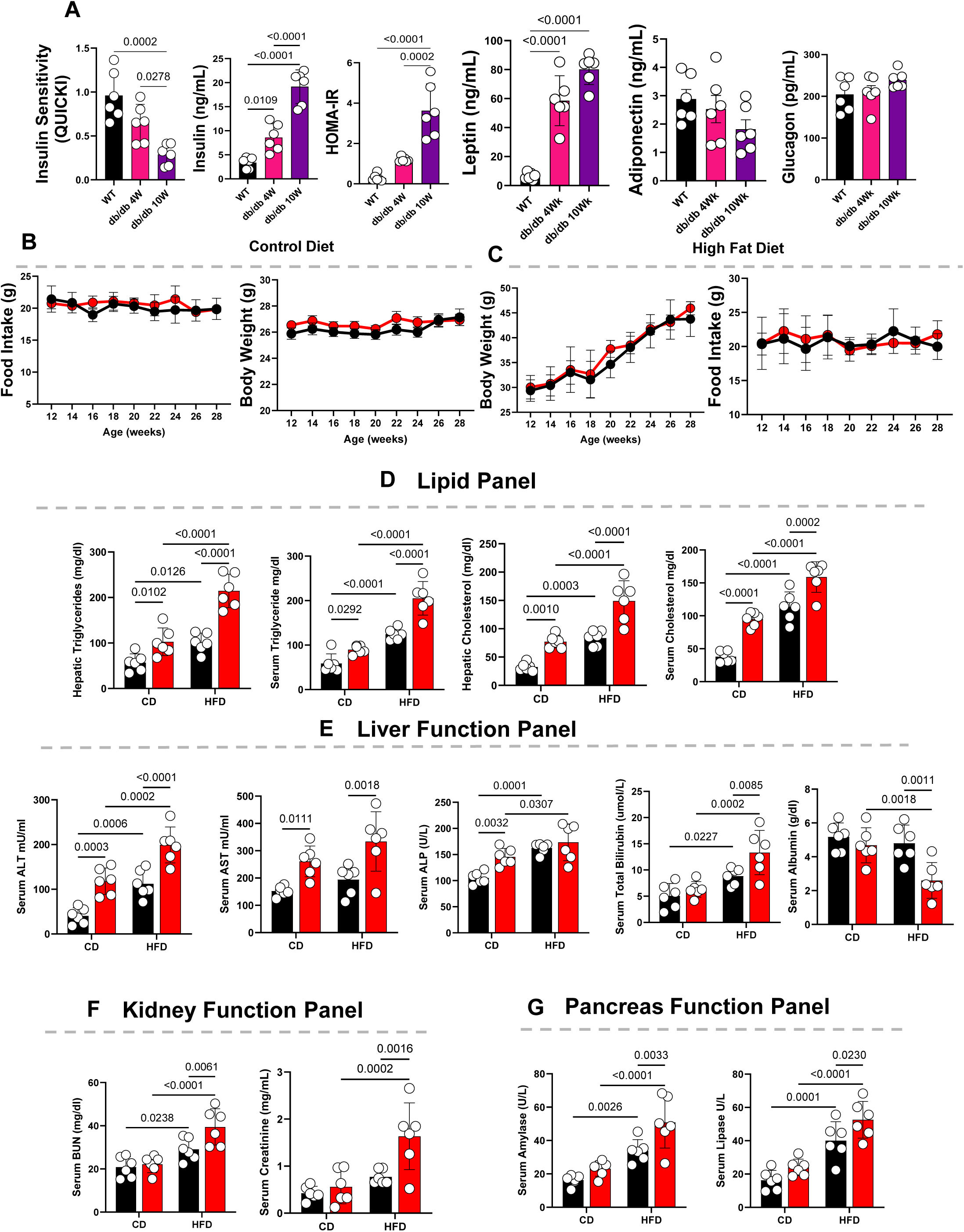
Hepatic NgBR Deficiency Disrupts Liver and Pancreatic Function in Metabolic Disease. **(A)** Metabolic hormone and insulin sensitivity profiling in wild-type (WT), early-stage diabetic (db/db, 4 weeks), and advanced diabetic (db/db, 10 weeks) mice showing reduced insulin sensitivity (QUICKI), increased fasting insulin, elevated HOMA-IR, increased leptin, reduced adiponectin, and increased glucagon with disease progression (n = 5–6 mice per group; exact *P* values shown). **(B–C)** Food intake and body weight in NgBR fl/fl and NgBR HepKO mice under control diet (CD; B) and high-fat diet (HFD; C), showing no major differences in caloric intake or body weight between genotypes. **(D)** Lipid profiling showing increased hepatic triglycerides, serum triglycerides, hepatic cholesterol, and serum cholesterol in NgBR HepKO mice, with greater elevations under HFD conditions (n = 5–6 mice per group; exact *P* values shown). **(E)** Liver function panel showing increased serum ALT, AST, ALP, and total bilirubin, together with reduced serum albumin in NgBR HepKO mice, consistent with impaired hepatic function (n = 5–6 mice per group; exact *P* values shown). **(F)** Kidney function panel showing increased serum BUN and creatinine in NgBR HepKO mice, indicating renal stress (n = 5–6 mice per group; exact *P* values shown). **(G)** Pancreatic enzyme analysis showing increased serum amylase and lipase levels in NgBR HepKO mice under both CD and HFD conditions (n = 5–6 mice per group; exact *P* values shown). Data are presented as mean ± SD denotes biological replicates (mice) as indicated. Statistical tests are described in Methods. Exact *P* values are provided in the figures.

**Extended Data Fig. 2:**
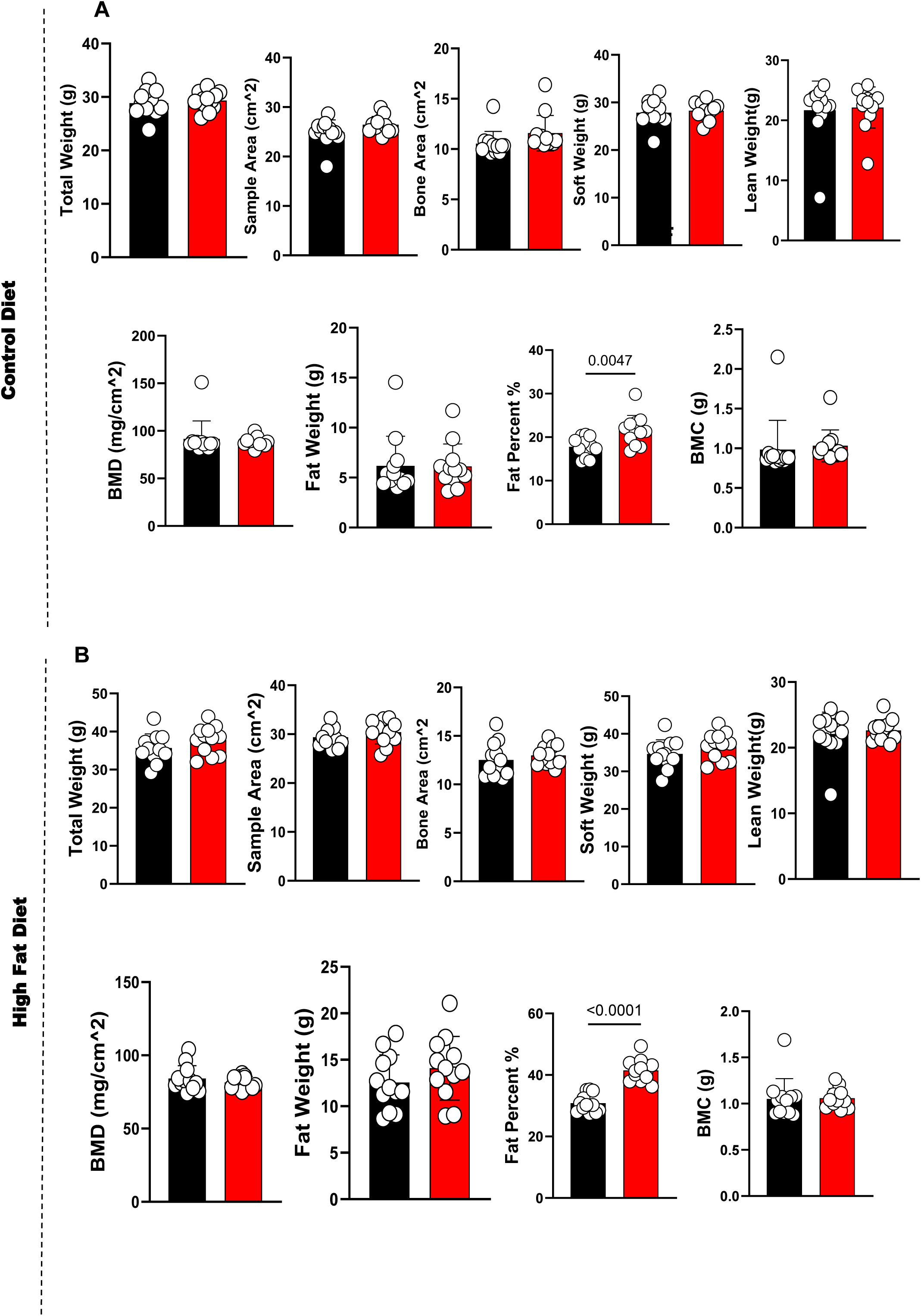
DEXA analysis reveals no significant differences between NgBRfl/fl and NgBR HepKO. **(A)** Dual-energy X-ray absorptiometry (DEXA) measurements in NgBR fl/fl and NgBR HepKO mice under control diet (CD) showing total body weight, sample area, bone area, soft tissue mass, lean mass, bone mineral density (BMD), fat mass, fat percentage, and bone mineral content (BMC). Most parameters are comparable between genotypes, with a modest increase in fat percentage observed in NgBR HepKO mice (n = 8–10 mice per group; exact *P* values shown). **(B)** DEXA analysis under high-fat diet (HFD) conditions showing similar total body weight, skeletal parameters (bone area, BMD, BMC), soft tissue mass, lean mass, and fat mass between NgBR fl/fl and NgBR HepKO mice. Fat percentage is modestly increased in NgBR HepKO mice under HFD (n = 8–10 mice per group; exact *P* values shown). Data are presented as mean ± SD denotes biological replicates (mice) as indicated. Statistical tests are described in Methods. Exact *P* values are provided in the figures.

**Extended Data Fig. 3:**
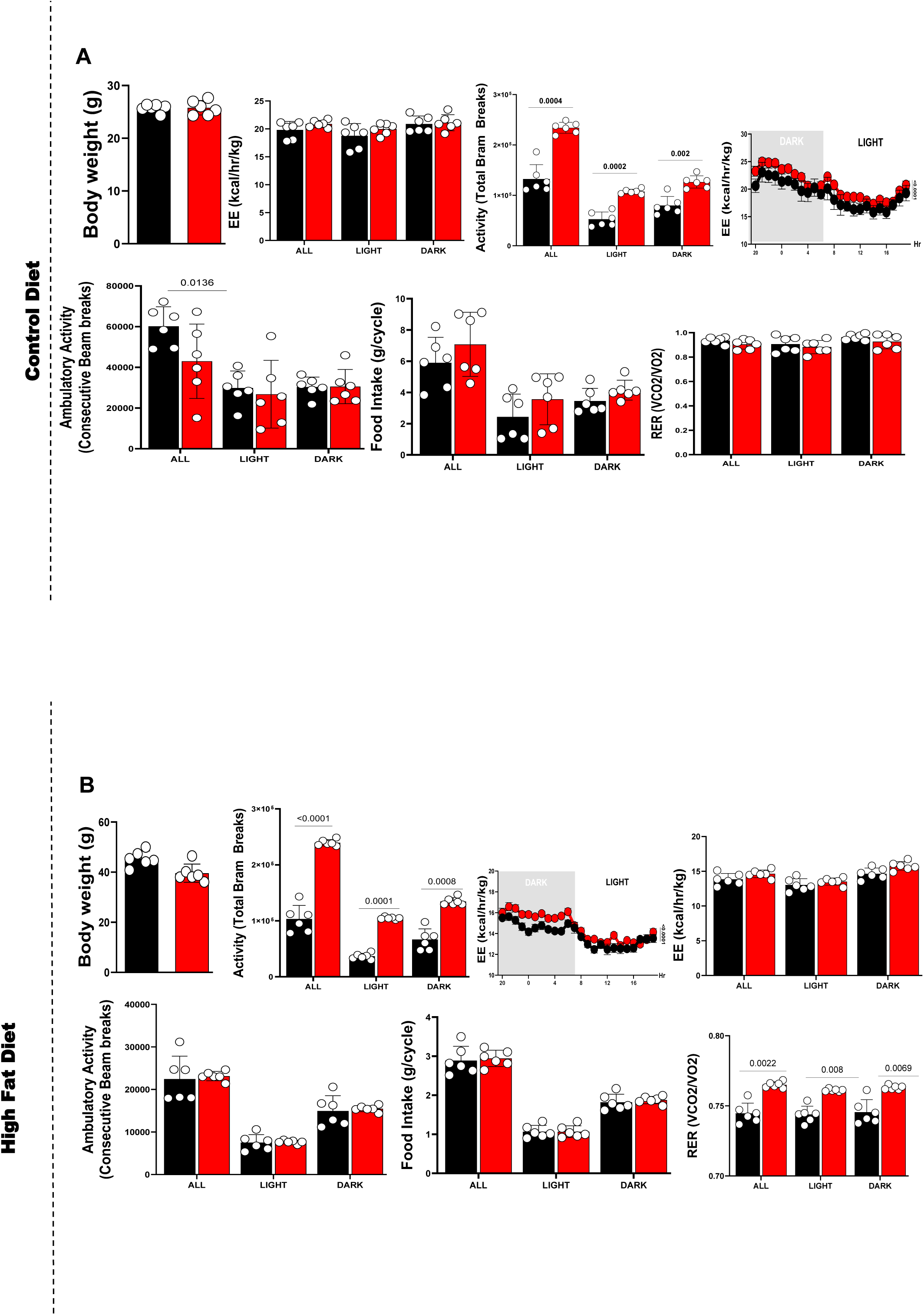
CLAMS analysis shows no significant differences between NgBR fl/fl and NgBR HepKO mice. **(A)** Comprehensive Lab Animal Monitoring System (CLAMS) analysis in NgBR fl/fl and NgBR HepKO mice under control diet (CD) conditions. Body weight, energy expenditure (EE), locomotor activity (beam breaks), food intake, and respiratory exchange ratio (RER) are shown across light and dark cycles. Most parameters are comparable between genotypes, with a modest increase in ambulatory activity in NgBR HepKO mice (n = 5–6 mice per group; exact *P* values shown). **(B)** CLAMS analysis under high-fat diet (HFD) conditions showing body weight, locomotor activity, energy expenditure, food intake, and RER across light and dark cycles. NgBR HepKO mice display increased locomotor activity, while EE, food intake, and RER remain comparable between genotypes (n = 5–6 mice per group; exact *P* values shown). Data are presented as mean ± SD denotes biological replicates (mice) as indicated. Statistical tests are described in Methods. For repeated measurements across light/dark cycles, two-way repeated-measures ANOVA was used. Exact *P* values are provided in the figures.

**Extended Data Fig. 4:**
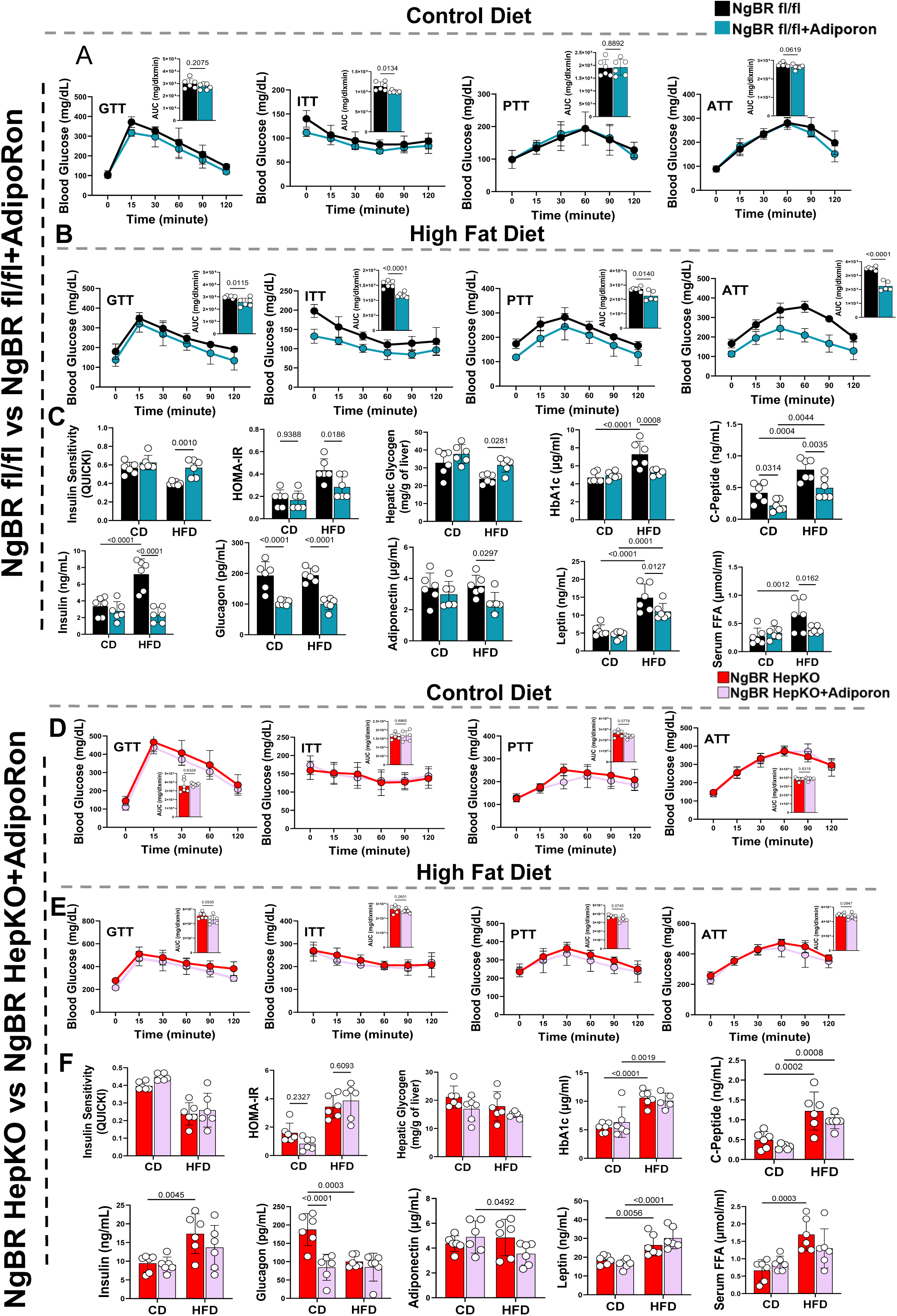
Loss of hepatic NgBR abolishes AdipoRon-induced improvements in glucose homeostasis and insulin sensitivity under control and high-fat diets. **(A–C)** NgBR fl/fl mice ± AdipoRon. **(A)** Control diet (CD). Glucose tolerance test (GTT), insulin tolerance test (ITT), pyruvate tolerance test (PTT), and alanine tolerance test (ATT) showed improved glucose handling and insulin sensitivity in AdipoRon-treated NgBR fl/fl mice. Insets show area under the curve (AUC) quantification (n = 6–8 mice per group; exact *P* values shown). **(B)** High-fat diet (HFD). AdipoRon treatment improves GTT, ITT, PTT, and ATT responses in NgBR fl/fl mice under HFD conditions, with corresponding reductions in AUC values (n = 6–8 mice per group; exact *P* values shown). **(C)** Metabolic profiling under CD and HFD showing improved insulin sensitivity (QUICKI), reduced HOMA-IR and HbA1c, increased hepatic glycogen, and modulation of circulating metabolic parameters, including insulin, glucagon, adiponectin, leptin, C-peptide, and serum free fatty acids (FFAs) in AdipoRon-treated NgBR fl/fl mice (n = 5–6 mice per group; exact *P* values shown). **(D–F)** NgBR HepKO mice ± AdipoRon. **(D)** Control diet (CD). GTT, ITT, PTT, and ATT showing minimal changes in glucose tolerance and insulin sensitivity following AdipoRon treatment in NgBR HepKO mice. Insets show AUC quantification (n = 6–8 mice per group; exact *P* values shown). **(E)** High-fat diet (HFD). AdipoRon treatment does not substantially improve glucose tolerance, insulin responsiveness, or substrate-driven glucose production in NgBR HepKO mice, as shown by GTT, ITT, PTT, and ATT curves and AUC analyses (n = 6–8 mice per group; exact *P* values shown). **(F)** Metabolic profiling under CD and HFD showing limited or absent improvement in insulin sensitivity (QUICKI), HOMA-IR, HbA1c, hepatic glycogen, and circulating metabolic markers in NgBR HepKO mice following AdipoRon treatment (n = 5–6 mice per group; exact *P* values shown). Data are presented as mean ± SD denotes biological replicates (mice) as indicated. Statistical tests are described in Methods. For time-course experiments (GTT, ITT, PTT, ATT), two-way repeated-measures ANOVA were used. Exact *P* values are provided in the figures.

**Extended Data Fig. 5:**
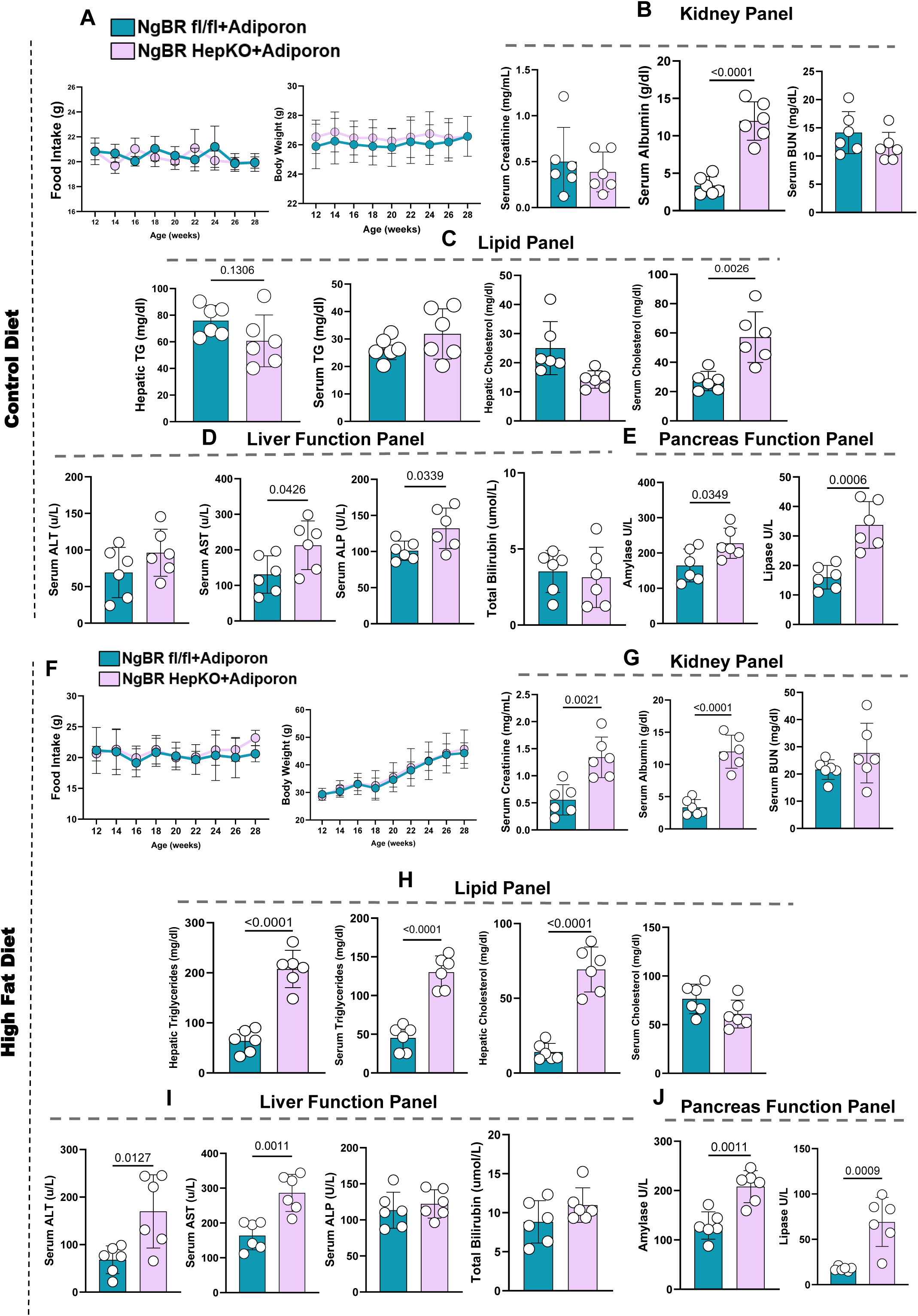
Adiponectin treatment improves metabolic and hepatic function in NgBR fl/fl mice but has no effect in NgBR HepKO mice. **(A, F)** Food intake and body weight during AdipoRon treatment in NgBR fl/fl and NgBR HepKO mice under control diet (CD; A) and high-fat diet (HFD; F) conditions. No major differences in feeding behavior or body weight trajectories are observed between groups. **(B, G)** Kidney function panels under CD (B) and HFD (G). AdipoRon treatment is associated with improved serum albumin and modest changes in BUN and creatinine in NgBR fl/fl mice, whereas NgBR HepKO mice show limited or no improvement (n = 5–6 mice per group; exact *P* values shown). **(C, H)** Lipid profiling under CD (C) and HFD (H). AdipoRon reduces hepatic triglycerides and circulating lipid parameters in NgBR fl/fl mice, while NgBR HepKO mice show minimal or no changes in lipid levels (n = 5–6 mice per group; exact *P* values shown). **(D, I)** Liver function panels under CD (D) and HFD (I). AdipoRon treatment is associated with reduced serum ALT, AST, ALP, and total bilirubin in NgBR fl/fl mice, whereas NgBR HepKO mice show persistent elevation of liver injury markers (n = 5–6 mice per group; exact *P* values shown). **(E, J)** Pancreatic enzyme analysis under CD (E) and HFD (J). AdipoRon reduces serum amylase and lipase levels in NgBR fl/fl mice, while NgBR HepKO mice show limited or no reduction in these markers (n = 5–6 mice per group; exact *P* values shown). Data are presented as mean ± SD denotes biological replicates (mice) as indicated. Statistical tests are described in Methods. Exact *P* values are provided in the figures.

**Extended Data Fig. 6:**
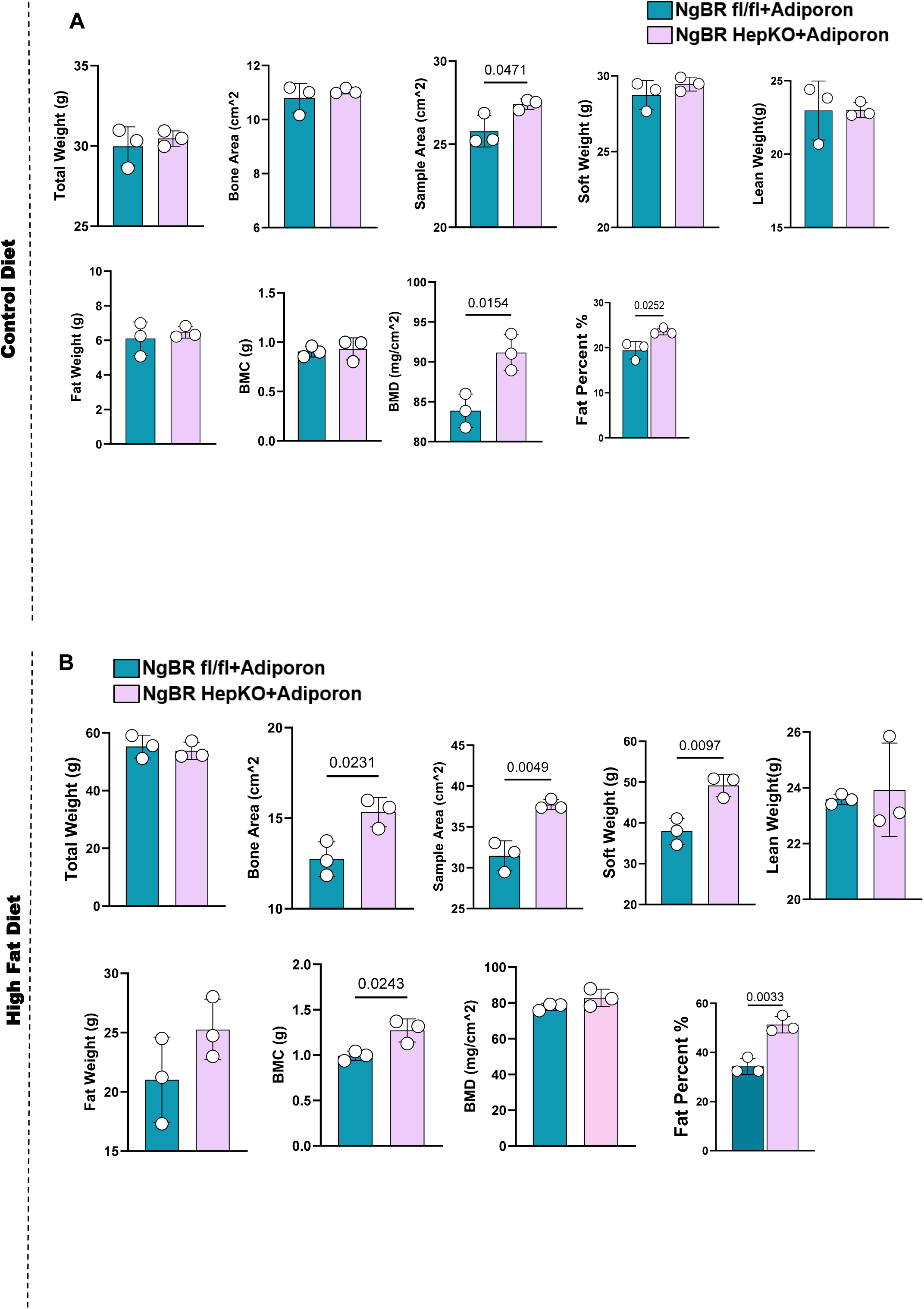
DEXA analysis shows no significant changes following adiponectin treatment in either NgBR fl/fl or NgBR HepKO mice. **(A)** Control diet (CD). Dual-energy X-ray absorptiometry (DEXA) measurements in NgBR fl/fl and NgBR HepKO mice treated with AdipoRon showing total body weight, bone area, sample area, soft tissue mass, lean mass, fat mass, bone mineral content (BMC), bone mineral density (BMD), and fat percentage. Most parameters are comparable between groups, with modest differences observed in sample area, BMD, and fat percentage (n = 5–6 mice per group; exact *P* values shown). **(B)** High-fat diet (HFD). DEXA analysis following AdipoRon treatment showing total body weight, bone area, sample area, soft tissue mass, lean mass, fat mass, BMC, BMD, and fat percentage. Overall body composition remains broadly comparable, with modest increases in selected parameters, including bone area, sample area, soft tissue mass, and fat percentage in NgBR HepKO mice (n = 5–6 mice per group; exact *P* values shown). Data are presented as mean ± SD denotes biological replicates (mice) as indicated. Statistical tests are described in Methods. Exact *P* values are provided in the figures.

**Extended Data Fig. 7:**
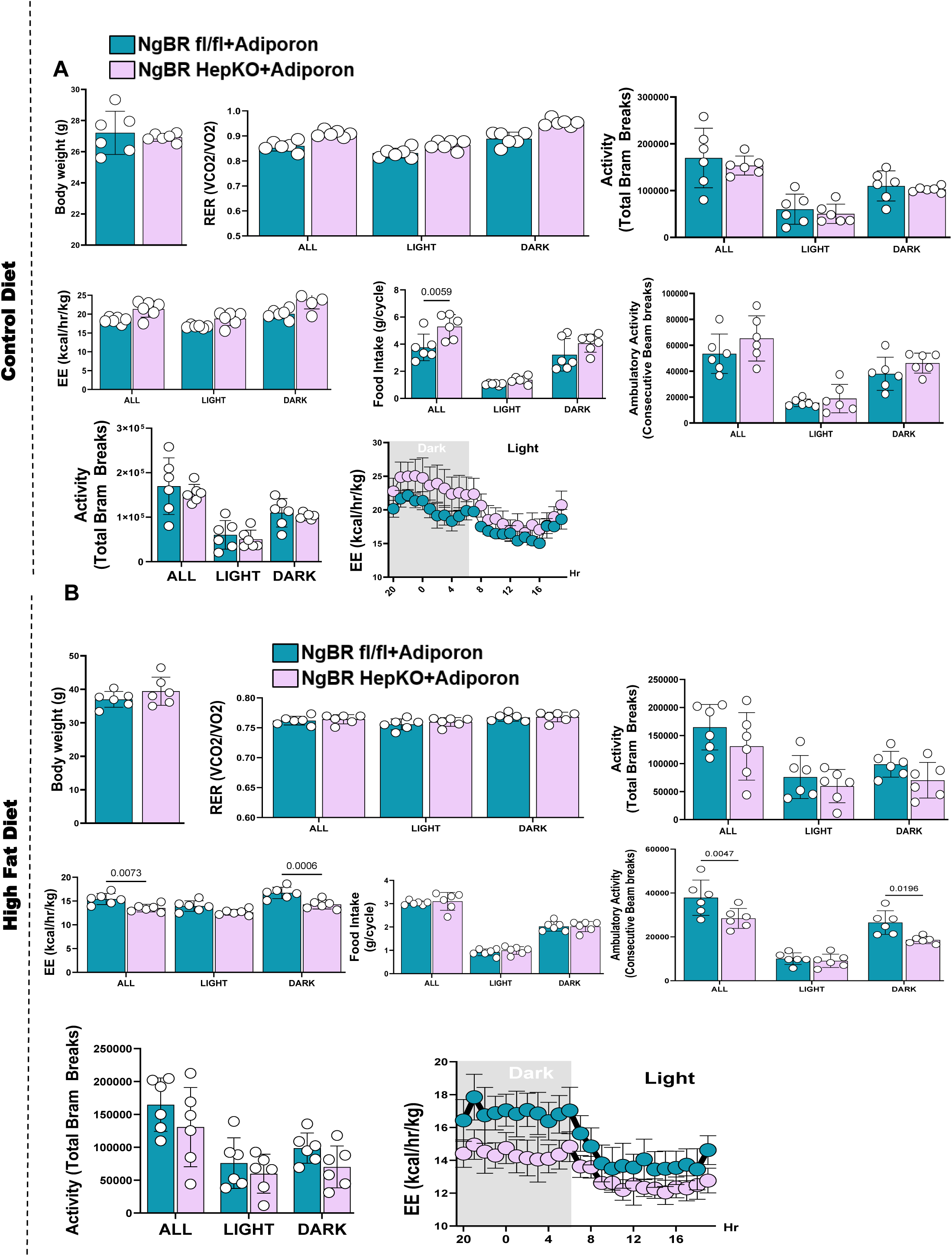
CLAMS analysis shows no significant effect on energy expenditure following AdipoRon treatment. **(A)** Control diet (CD). Comprehensive Lab Animal Monitoring System (CLAMS) analysis in NgBR fl/fl and NgBR HepKO mice following AdipoRon treatment showing body weight, respiratory exchange ratio (RER), energy expenditure (EE), locomotor activity (beam breaks), and food intake across 24-hour cycles. Light- and dark-phase analyses demonstrate comparable RER and EE profiles between groups, with similar diurnal patterns of energy expenditure (n = 5–6 mice per group; exact *P* values shown). **(B)** High-fat diet (HFD). CLAMS analysis under HFD conditions showing RER, EE, locomotor activity, and food intake in NgBR fl/fl and NgBR HepKO mice following AdipoRon treatment. Overall energy balance parameters remain comparable between groups, with only minor variations in activity that do not translate into consistent changes in energy expenditure or substrate utilization (n = 5–6 mice per group; exact *P* values shown). Data are presented as mean ± SD denotes biological replicates (mice) as indicated. Statistical tests are described in Methods. For repeated measurements across time (light/dark cycles and hourly traces), two-way repeated-measures ANOVA was used. Exact *P* values are provided in the figures.

**Extended Data Fig. 8:**
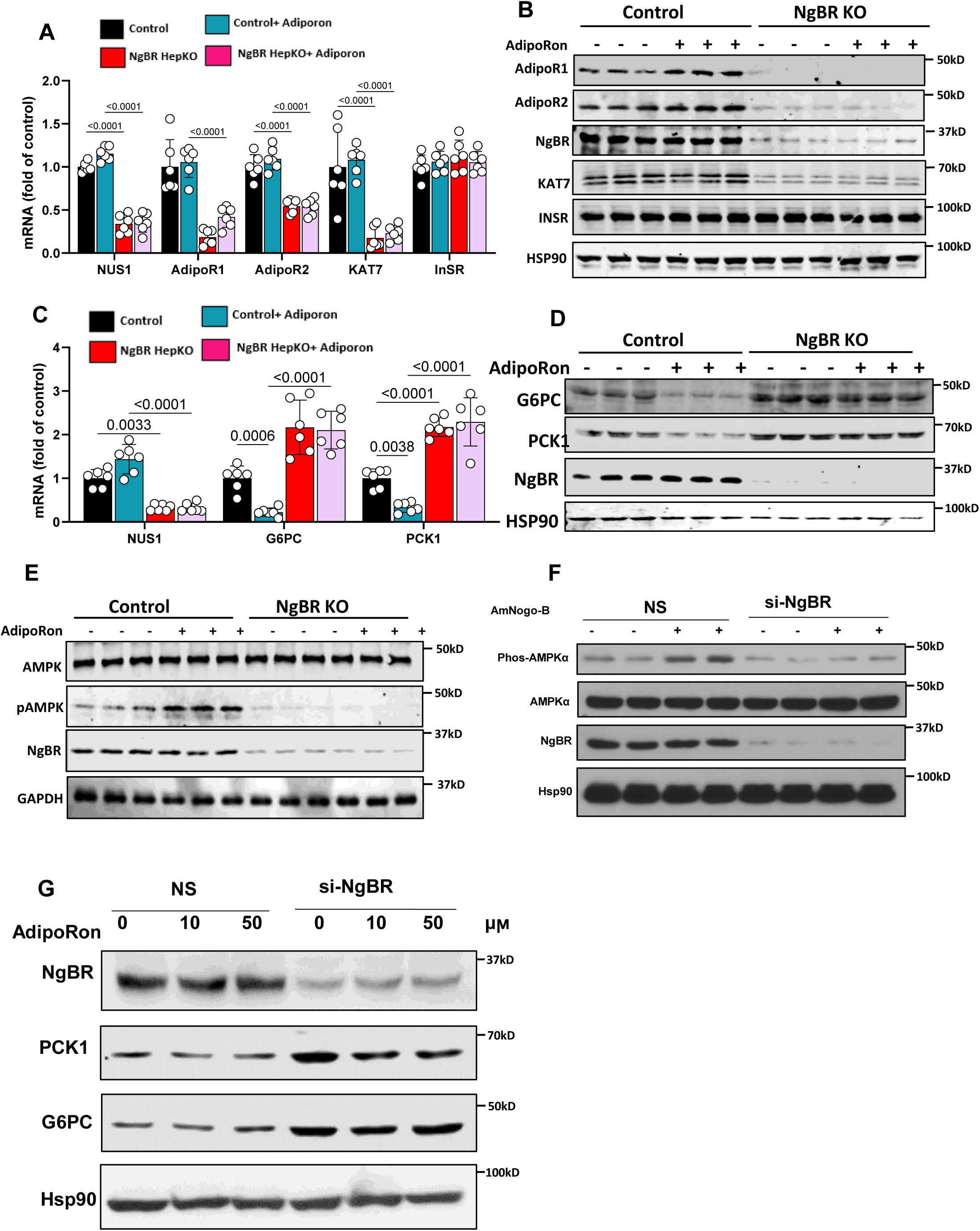
NgBR Is Essential for Adiponectin-Induced AMPK Activation and Suppression of Hepatic Gluconeogenesis in HepG2 cells. **(A)** qPCR analysis in HepG2 cells showing that AdipoRon treatment increases expression of **NUS1, ADIPOR1, ADIPOR2, and KAT7** in control cells, whereas NgBR knockout markedly attenuates this response. INSR expression remains largely unchanged (n = 4 independent experiments; exact *P* values shown). **(B)** Immunoblot analysis confirming increased AdipoR1, AdipoR2, and KAT7 protein levels following AdipoRon treatment in control cells, with reduced or absent induction in NgBR knockout cells. **(C)** qPCR analysis showing that AdipoRon suppresses **G6PC** and **PCK1** expression in control HepG2 cells, whereas this repression is markedly blunted in NgBR-deficient cells (n = 4 independent experiments; exact *P* values shown). **(D)** Immunoblot validation demonstrates reduced G6PC and PCK1 protein levels following AdipoRon treatment in control cells, with diminished suppression in NgBR knockout cells. **(E)** Immunoblot analysis showing that AdipoRon induces AMPK phosphorylation in control HepG2 cells, whereas this response is markedly reduced in NgBR knockout cells. **(F)** siRNA-mediated NgBR silencing (si-NgBR) confirms reduced AMPK phosphorylation following stimulation with recombinant Nogo-B, indicating that NgBR is required for efficient downstream signaling. **(G)** Dose–response analysis showing that increasing concentrations of AdipoRon (0–50 μM) progressively suppress PCK1 and G6PC protein expression in control cells, whereas NgBR-deficient cells exhibit attenuated responsiveness. Data are presented as mean ± SD denotes independent biological replicates as indicated. Statistical tests are described in Methods. Exact *P* values are provided in the figures.

**Extended Data Fig. 9:**
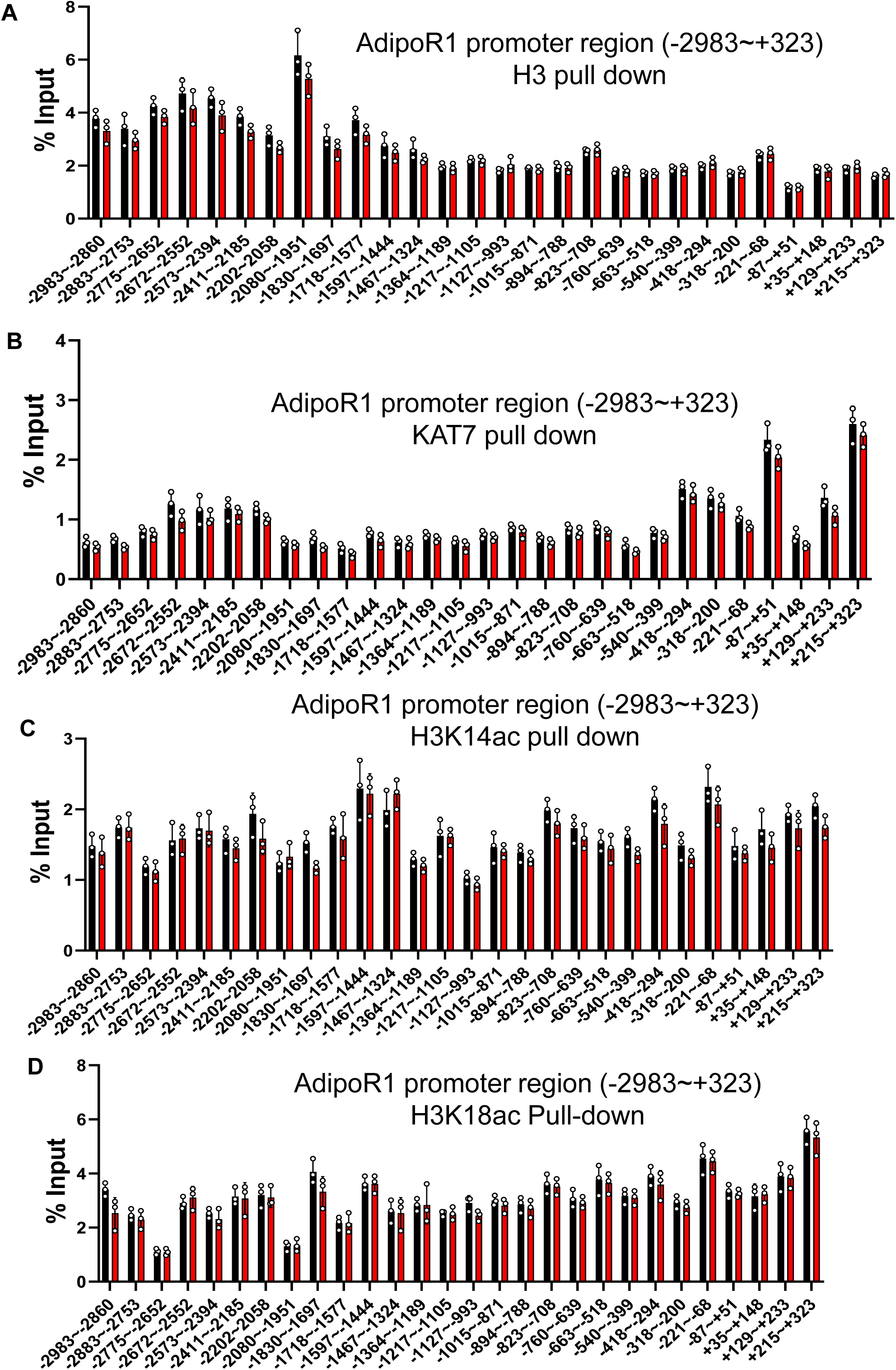
KAT7-dependent histone acetylation is associated with a permissive chromatin landscape at the AdipoR1 promoter. Chromatin immunoprecipitation followed by quantitative PCR (ChIP–qPCR) was performed across the **AdipoR1 promoter** (−2983 to +323 relative to the transcription start site) using tiled amplicons to assess chromatin occupancy and histone acetylation. Enrichment is expressed as percentage of input DNA. **(A)** Total histone H3 occupancy showing nucleosome distribution across the AdipoR1 promoter. **(B)** KAT7 (HBO1) ChIP demonstrates selective occupancy at discrete regulatory regions. **(C–D)** ChIP–qPCR analysis of activating histone marks **H3K14ac** and **H3K18ac**, showing enrichment across multiple promoter regions. KAT7 occupancy overlaps with regions enriched for H3K14ac and H3K18ac, consistent with coordinated histone acetyltransferase activity at the AdipoR1 locus. These data support a model in which KAT7-dependent histone acetylation contributes to the maintenance of a transcriptionally permissive chromatin state. Data is presented as mean ± SD with individual biological replicates shown. n denotes independent experiments as indicated. Statistical tests are described in Methods. Exact *P* values are provided where applicable.

**Extended Data Fig. 10:**
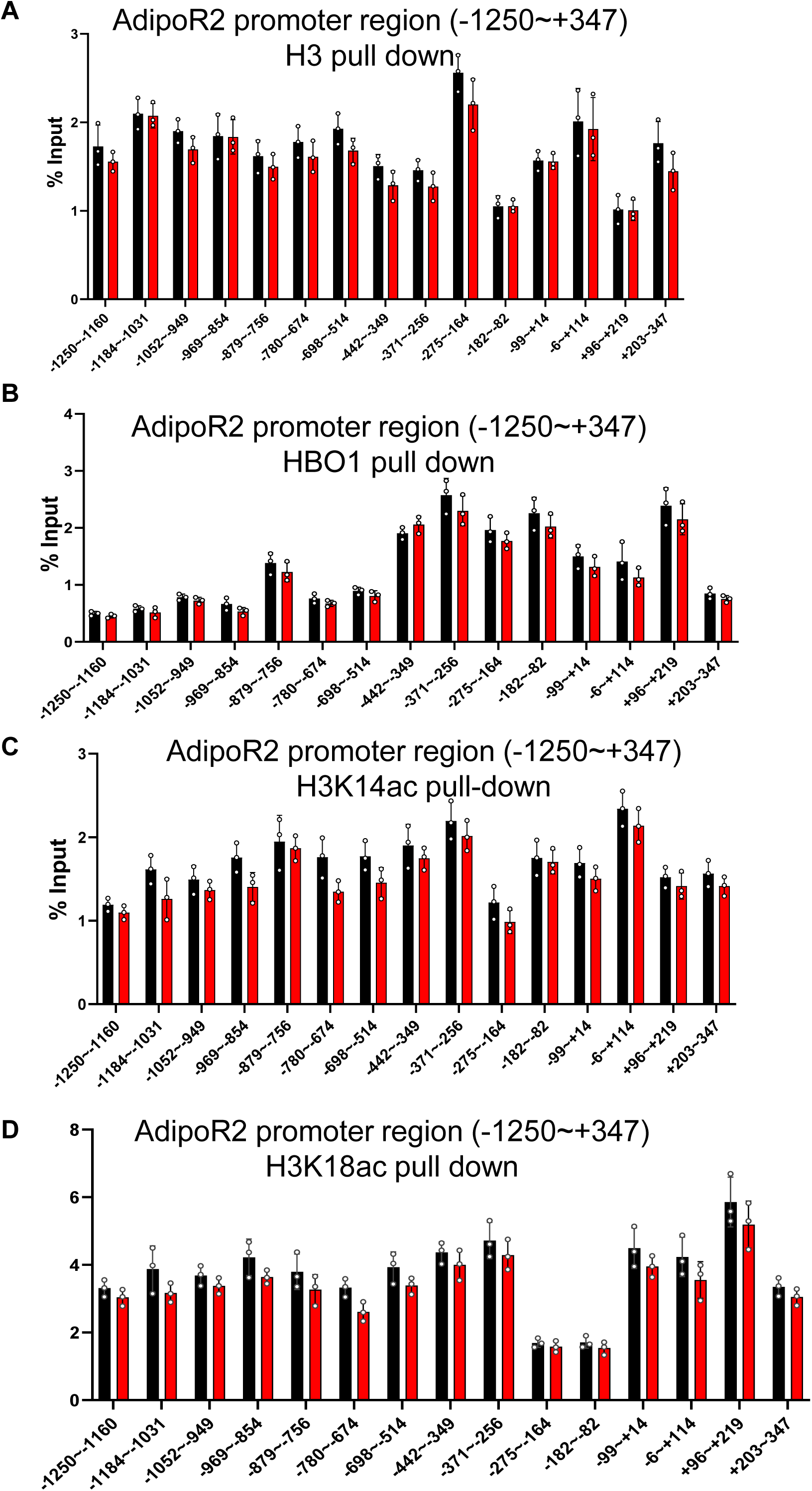
Epigenetic profiling of the AdipoR2 promoter shows coordinated KAT7 occupancy and histone acetylation. Chromatin immunoprecipitation followed by quantitative PCR (ChIP–qPCR) was performed across the **AdipoR2 promoter** (−1250 to +347 relative to the transcription start site) using tiled amplicons to assess chromatin occupancy and histone acetylation. Enrichment is expressed as percentage of input DNA. **(A)** Total histone H3 occupancy showing nucleosome distribution across the AdipoR2 promoter. **(B)** KAT7 (HBO1) ChIP demonstrates selective occupancy at discrete regulatory regions. **(C–D)** ChIP–qPCR analysis of activating histone marks **H3K14ac** and **H3K18ac**, showing enrichment across multiple promoter regions. KAT7 occupancy overlaps with regions enriched for H3K14ac and H3K18ac, consistent with coordinated histone acetyltransferase activity at the AdipoR2 locus. These data support a model in which KAT7-dependent histone acetylation contributes to maintenance of a transcriptionally permissive chromatin state at the AdipoR2 promoter. Data are presented as mean ± SD with individual biological replicates shown. n denotes independent experiments as indicated. Statistical tests are described in Methods. Exact *P* values are provided where applicable.

**Extended Data Fig. 11:**
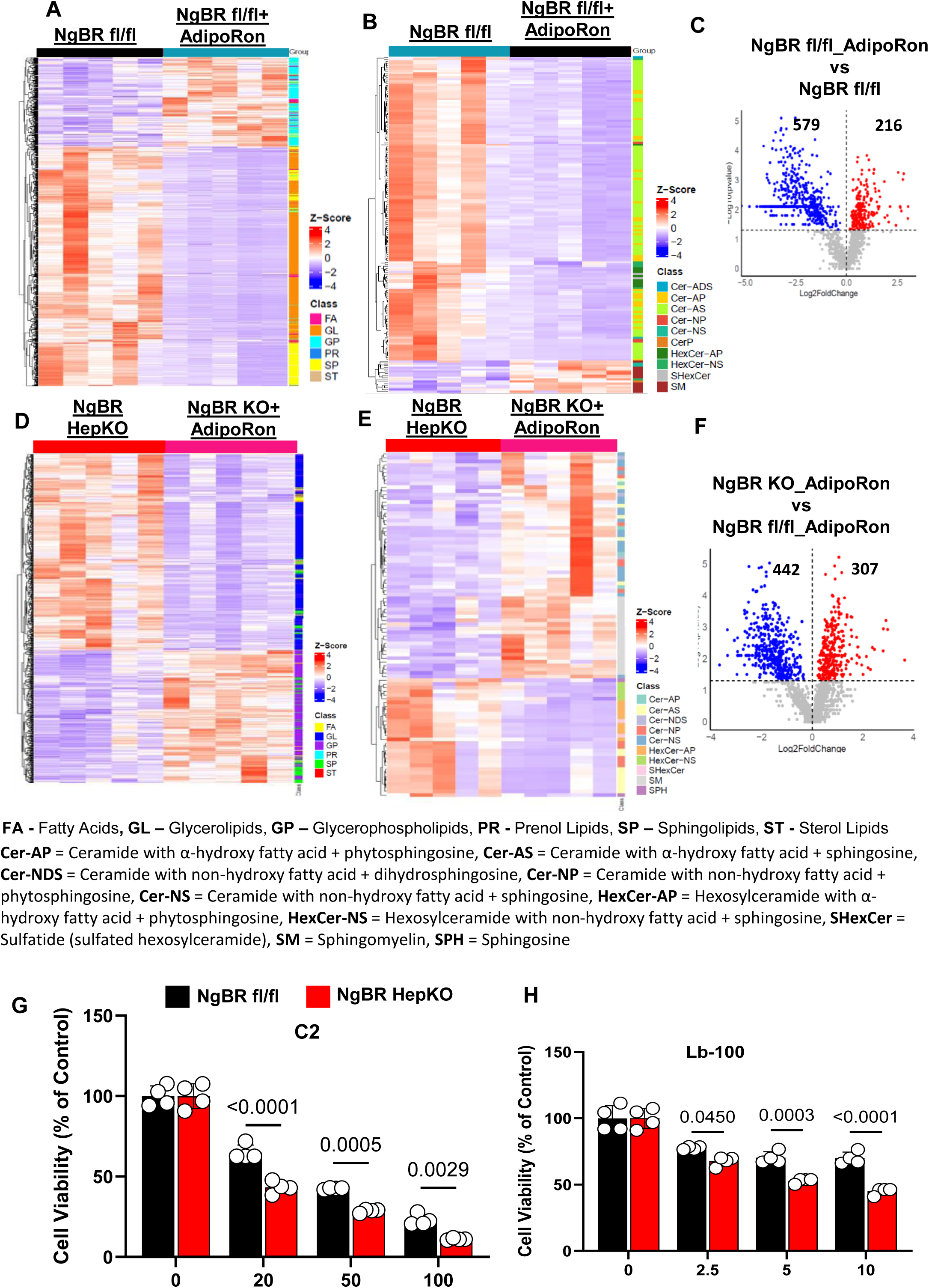
Adiponectin Remodels Hepatic Lipid Profiles in NgBR fl/fl Mice but Not in NgBR HepKO Mice. **(A–B)** Unsupervised hierarchical clustering heatmaps showing global hepatic lipidomic profiles in NgBR fl/fl mice before and after AdipoRon treatment. AdipoRon induces broad remodeling across multiple lipid subclasses, including fatty acids (FA), glycerolipids (GL), glycerophospholipids (GP), prenol lipids (PR), sphingolipids (SP), and sterol lipids (ST). **(C)** Volcano plot comparing AdipoRon-treated and untreated NgBR fl/fl livers, showing differential abundance of lipid species with a predominance of decreased lipids following treatment. **(D–E)** Heatmaps of hepatic lipidomic profiles in NgBR HepKO mice before and after AdipoRon treatment. In contrast to NgBR fl/fl mice, NgBR HepKO livers exhibit limited lipid remodeling, with persistence of multiple sphingolipid and ceramide-associated species. **(F)** Volcano plot comparing AdipoRon-treated NgBR HepKO and NgBR fl/fl livers, demonstrating a distinct lipidomic response in NgBR-deficient mice. **(G)** Cell viability assay showing reduced survival of NgBR HepKO hepatocytes compared with NgBR fl/fl controls following treatment with the membrane-permeable ceramide analog C2 across increasing concentrations. **(H)** Cell viability assay showing decreased survival of NgBR HepKO hepatocytes compared with controls following treatment with the PP2A inhibitor LB-100, indicating increased sensitivity to PP2A-associated stress. Data are presented as mean ± SD, and biological replicates are indicated. Statistical tests are described in Methods. Exact *P* values are provided in the figures. **Abbreviations** FA, fatty acids; GL, glycerolipids; GP, glycerophospholipids; PR, prenol lipids; SP, sphingolipids; ST, sterol lipids; Cer, ceramide; HexCer, hexosylceramide; SHexCer, sulfatide; SM, sphingomyelin; SPH, sphingosine.

**Extended Data Fig. 12:**
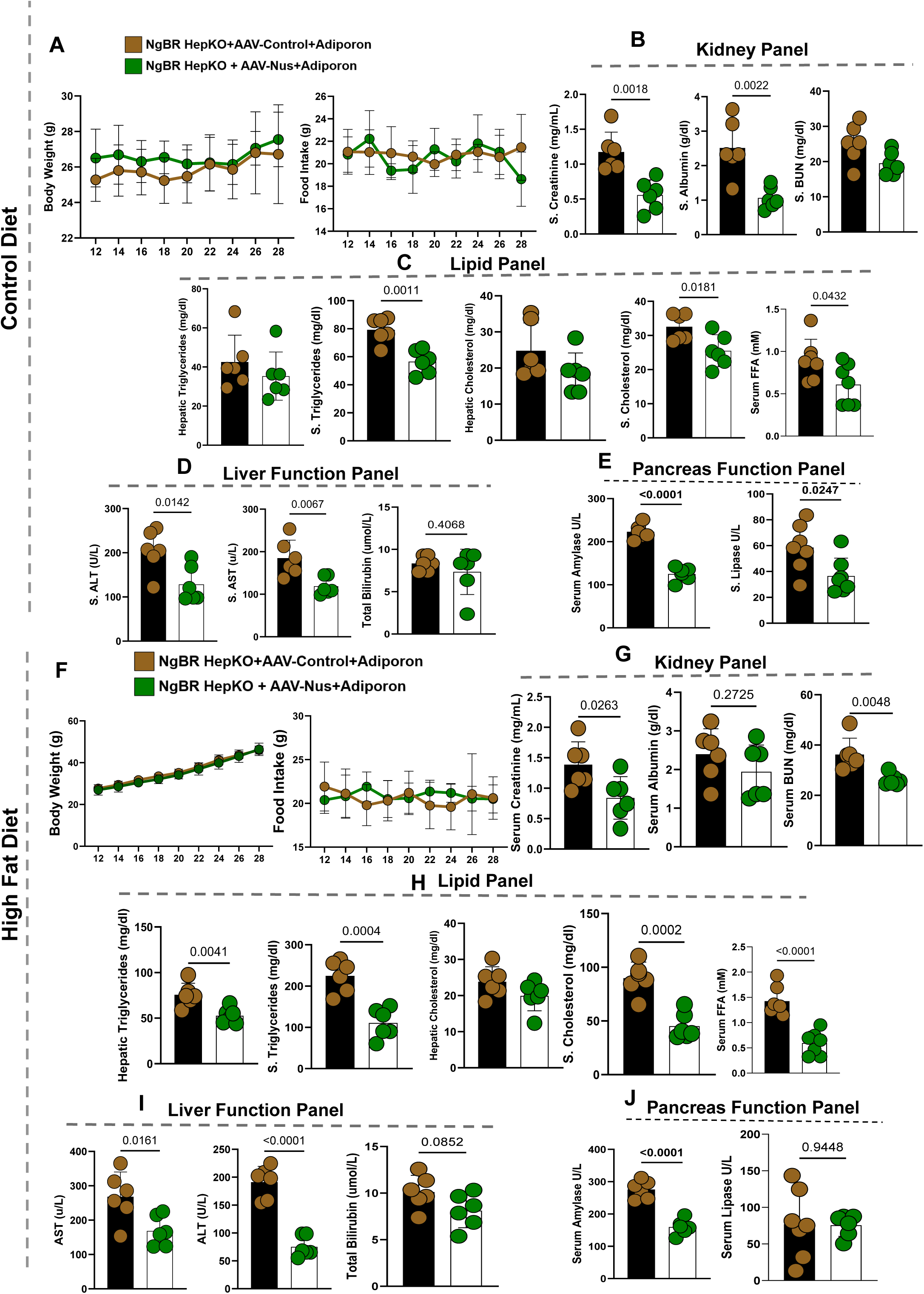
AAV-Mediated NgBR Overexpression Restores Insulin Sensitivity and Glucose Homeostasis in AdipoRon-Treated NgBR HepKO Mice. **(A–E)** Control diet (CD). Metabolic and biochemical assessment of NgBR HepKO mice treated with AAV-Control or AAV-NUS1 during AdipoRon administration. **(A)** Body weight and food intake over the study period, showing comparable trajectories between groups. **(B)** Kidney function panel showing differences in serum creatinine, albumin, and BUN between groups. **(C)** Lipid panel showing reduced hepatic triglycerides and changes in circulating triglycerides, cholesterol, and free fatty acids (FFAs) in AAV-NUS1–treated mice. **(D)** Liver function panel showing differences in serum ALT, AST, and total bilirubin between groups. **(E)** Pancreatic enzyme panel showing reduced serum amylase and lipase levels in AAV-NUS1–treated mice. **(F–J)** High-fat diet (HFD). Metabolic assessment of AAV-mediated NgBR re-expression under diet-induced stress. **(F)** Body weight and food intake over time, showing comparable patterns between groups. **(G)** Kidney function panel showing differences in serum creatinine, albumin, and BUN following AAV-NUS1 treatment. **(H)** Lipid panel showing changes in hepatic triglycerides, circulating triglycerides, cholesterol, and FFAs. **(I)** Liver function panel showing reduced ALT and AST levels with modest changes in total bilirubin. **(J)** Pancreatic enzyme panel showing reduced serum amylase, with limited change in lipase levels. Data are presented as mean ± SD denotes biological replicates (mice) as indicated. Statistical tests are described in Methods. Exact *P* values are provided in the figures.

**Extended Data Fig. 13:**
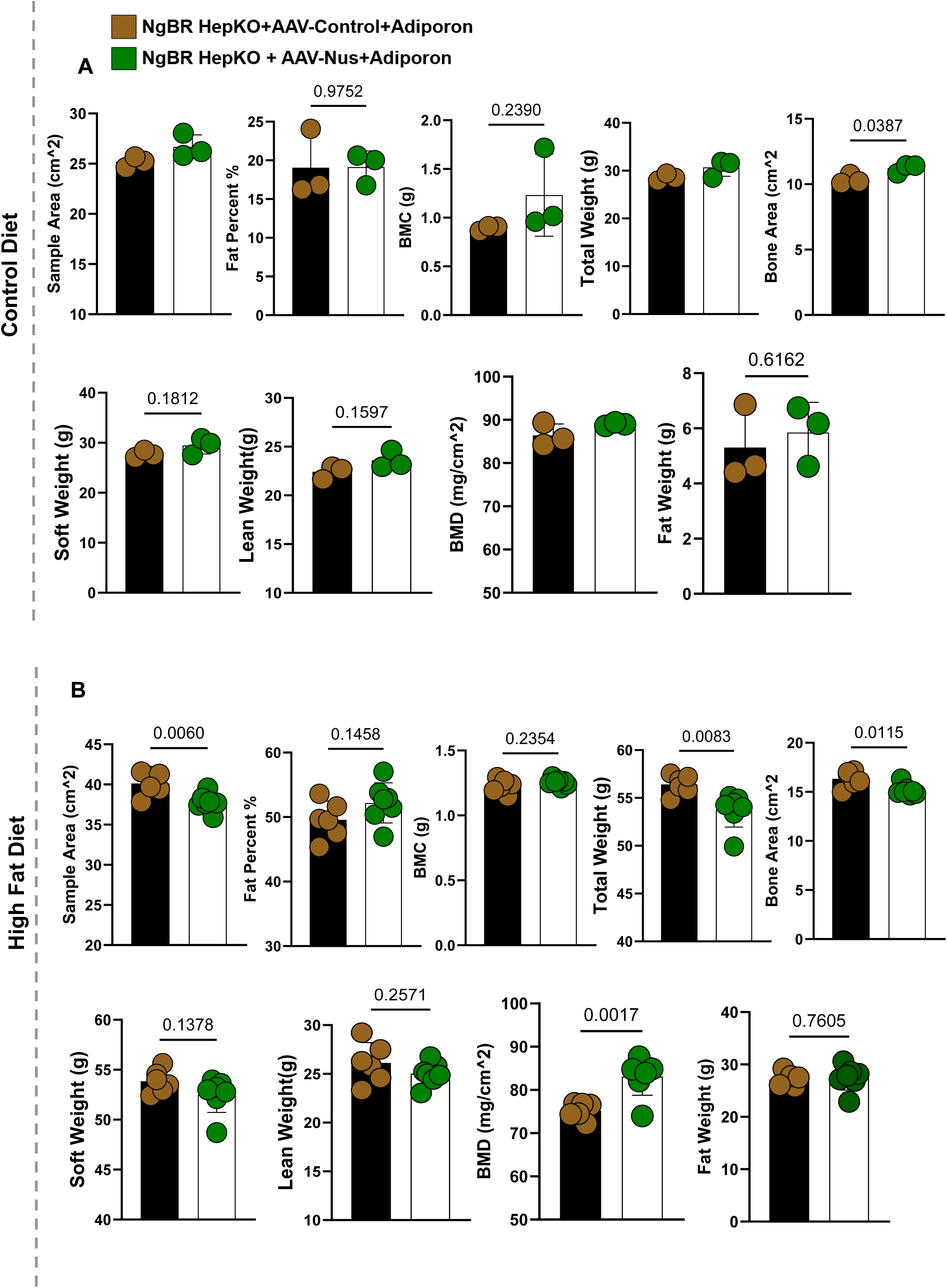
DEXA analysis demonstrates improved body composition and mass following AAV-mediated NgBR restoration. **(A)** Control diet (CD). Dual-energy X-ray absorptiometry (DEXA) measurements comparing NgBR HepKO mice treated with AAV-Control + AdipoRon versus AAV-NUS1 + AdipoRon. Body composition parameters, including total weight, lean mass, fat mass, fat percentage, soft tissue mass, bone mineral density (BMD), bone mineral content (BMC), and sample area, were largely comparable between groups. A modest difference in bone area was observed, while other parameters showed no consistent changes. **(B)** High-fat diet (HFD). DEXA measurements under metabolic stress conditions. Differences were observed in selected parameters, including sample area, total body weight, bone area, and BMD. Lean mass, soft tissue mass, fat mass, fat percentage, and BMC were not consistently altered between groups. Data are presented as mean ± SD denotes biological replicates (mice) as indicated. Statistical tests are described in Methods. Exact *P* values are provided in the figure panels.

